# A missense in HSF2BP causing Primary Ovarian Insufficiency affects meiotic recombination by its novel interactor C19ORF57/MIDAP

**DOI:** 10.1101/2020.03.05.978007

**Authors:** Natalia Felipe-Medina, Sandrine Caburet, Fernando Sánchez-Sáez, Yazmine B. Condezo, Dirk de Rooij, Laura Gómez-H, Rodrigo García-Valiente, Anne-Laure Todeschini, Paloma Duque, Manuel Sánchez-Martín, Stavit A. Shalev, Elena Llano, Reiner Veitia, Alberto M. Pendás

**Author notes:** Authors for Correspondence Alberto M. Pendas (ORCID: 0000-0001-9264-3721), Reiner Veitia (ORCID: 0000-0002-4100-2681). These authors contributed equally (Natalia Felipe-Medina, Sandrine Caburet).

## Abstract

Primary Ovarian Insufficiency (POI) is a major cause of infertility, but its etiology remains poorly understood. Using whole-exome sequencing in a family with 3 cases of POI, we identified the candidate missense variant S167L in *HSF2BP*, an essential meiotic gene. Functional analysis of the HSF2BP-S167L variant in mouse, compared to a new *HSF2BP* knock-out mouse showed that it behaves as a hypomorphic allele. HSF2BP-S167L females show reduced fertility with small litter sizes. To obtain mechanistic insights, we identified C19ORF57/MIDAP as a strong interactor and stabilizer of HSF2BP by forming a higher-order macromolecular structure involving BRCA2, RAD51, RPA and PALB2. Meiocytes bearing the HSF2BP-S167L mutation showed a strongly decreased expression of both MIDAP and HSF2BP at the recombination nodules. Although HSF2BP-S167L does not affect heterodimerization between HSF2BP and MIDAP, it promotes a lower expression of both proteins and a less proficient activity in replacing RPA by the recombinases RAD51/DMC1, thus leading to a lower frequency of cross-overs. Our results provide insights into the molecular mechanism of two novel actors of meiosis underlying non-syndromic ovarian insufficiency.

**Summary:** Felipe-Medina et al. describe a missense variant in the meiotic gene *HSF2BP* in a consanguineous family with Premature Ovarian Insufficiency, and characterize it as an hypormorphic allele, that *in vivo* impairs its dimerization with a novel meiotic actor, MIDAP/ C19ORF57, and affect recombination at double-strand DNA breaks.

## Introduction

The process of gametogenesis is one of the most complex and highly regulated differentiation programs. It involves a unique reductional cell division, known as meiosis, to generate highly specialized gametes. The outcome of meiosis is the production of oocytes and spermatozoa, which are the most distinctive cells of an adult organism and are essential for the faithful transmission of the genome across generations.

The meiotic division is an orderly process that results in the pairing and synapsis of homologous chromosomes and cross-over (CO) formation, which ultimately enable homologous chromosomes segregation (Hunter, 2015; Loidl, 2016; Zickler and Kleckner, 2015). In mammals, pairing of homologs is dependent on the repair of self-induced double-strand breaks (DSBs) during prophase I by homologous recombination (Handel and Schimenti, 2010) and it precedes the intimate alignment of homologous chromosomes (synapsis) through the zipper-like structure that is the synaptonemal complex (SC) (Cahoon and Hawley, 2016). The SC proteinaceous tripartite assembly also provides the structural framework for DSBs repair (Baudat et al., 2013), as epitomized by the tight association of the recombination nodules (RN, multicomponent recombinogenic factories) and the axial elements of the SC (Zickler and Kleckner, 2015).

Meiotic DSBs repair is an evolutionarily conserved pathway that is highly regulated to promote the formation of at least one crossover per bivalent. Chromosome junction together with cohesion ensure tension between sister kinetochores during the first reductional division. As other DNA repair processes, proper meiotic recombination is essential for genome stability and alterations can result in infertility, miscarriage and birth defects (Geisinger and Benavente, 2016; Handel and Schimenti, 2010; Webster and Schuh, 2017).

Infertility refers to failure of a couple to conceive and affects 10–15% of couples (Isaksson and Tiitinen, 2004). Infertility can be due to female factors, male factors, a combination of both or to unknown causes, each category representing approximately 25% of cases (Isaksson and Tiitinen, 2004; Matzuk and Lamb, 2008). There are several underlying causes and physiological, genetic and even environmental and social factors can play a role. Forward and reverse genetic analyses in model organisms have identified multiple molecular pathways that regulate fertility and have allowed inferring reasonable estimates of the number of protein-coding genes essential for fertility (de Rooij and de Boer, 2003; Schimenti and Handel, 2018).

Primary ovarian insufficiency (POI) is a major cause of female infertility and affects about 1–3% of women under 40 years of age. It is characterized by cessation of menstruation before the age of 40 years. POI results from a depletion of the ovarian follicle pool and can be isolated or syndromic. Genetic causes of POI account for approximately 20% of cases (Rossetti et al., 2017). Whereas individual infertility-causing pathogenic variants are inherently unlikely to spread in a population, they can be observed within families, especially in the event of consanguinity. They provide crucial insights into the function of the genes and molecular mechanisms that they disrupt. Over the last decade, several causative genes have been found using whole exome sequencing (WES) in POI pedigrees. In particular, pathogenic variants in genes involved in DNA replication, recombination or repair, such as *STAG3, SYCE1, HFM1, MSH5* and *MEIOB* have been formally implicated in this condition by ourselves and others (Caburet et al., 2014; Caburet et al., 2019b; de Vries et al., 2014; Guo et al., 2017; Wang et al., 2014).

In this study, we identified in a consanguineous family with POI a candidate missense variant in *HSF2BP*, an essential yet poorly-studied meiotic gene. *HSF2BP* gene encodes a binding protein of the heat shock response transcription factor HSF2 (Yoshima et al., 1998). During the course of this work and in agreement with our results, two independent groups showed that HSF2BP is essential for meiotic recombination through its ability to interact with BRCA2 (Brandsma et al., 2019; Zhang et al., 2019). Here, we report that the introduction of the missense variant HSF2BP-S167L in the mouse leads to subfertility and DNA repair defects during prophase I. In addition, we identified a protein complex composed of BRCA2, HSF2BP, and the as yet unexplored C19ORF57/MIDAP (meiotic double-stranded break BRCA2/HSF2BP complex associated protein) as a key component of the meiotic recombination machinery. Our studies show that a single substitution (S167L) in HSF2BP leads to a reduction in the loading of both MIDAP and HSF2BP at the recombination nodules. Furthermore, our results suggest that meiotic progression requires a critical threshold level of MIDAP for the ulterior loading of recombinases to the RN.

## Results

### Clinical cases

The parents are first-degree cousins of Moslem-Arab origin. Of the five daughters, three are affected with POI and presented with early secondary amenorrhea. They had menarche at normal age (at 13-14) but with irregular menstruation that stopped around 25. Only one of the POI patients could conceive with the help of a fertility treatment (see pedigree in Figure S1). In order to identify the genetic basis of this familial POI case, we performed WES on genomic DNA from two POI patients, III-2 and III-3, and their fertile sister III-10 (Table S1). Variants were filtered on the basis of i) their homozygosity in the patients, ii) their heterozygosity or absence in the fertile sister, iii) their absence in unrelated fertile in-house controls and iv) a minor allele frequency (MAF) below 0.01 in all available databases (Table S2). The identified variant was a missense substitution located in the *HSF2BP* gene, rs200655253 (21:43630396 G>A, GRCh38). The variant lies within the sixth exon of the reference transcript ENST00000291560.7 (NM_007031.2:c.500C>T) and changes a TCG codon into a TTG (NP_008962.1:p.Ser167Leu). It is very rare (VAF 0.0001845 in the GnomAD database and 0.0005 in the GME Variome dedicated to Middle-East populations) and absent at the homozygous state in all databases. The variant was verified by Sanger sequencing and was found to segregate in a Mendelian fashion within the family: the affected twin III-1 was homozygous for the variant and both parents and fertile siblings were heterozygous carriers (Figure S2). Serine 167 is a highly conserved position and the S167L variant is predicted to be pathogenic or deleterious by 11 out of the 18 pathogenicity predictors available in dbNSFP 3.5. (Table S3, Figures S3 and S4).

HSF2BP was first identified as a testis-specific interactor of HSF2, able to sequester this transcription factor in the cytoplasm (Yoshima et al., 1998). Another partner, basonuclin 1 (BCN1), has been recently implicated in POI (Tsuiko et al., 2016; Wu et al., 2013; Zhang et al., 2018). We therefore tested a possible impact of the S167L variant in HSF2BP on its interaction with HSF2 and BCN1 by a two-hybrid system in cell culture, but did not observe any difference with the wild-type (WT) protein.

### Mice with the HSF2BP S167L variant show a partial reduction of fertility

In order to confirm the causality of the S167L variant in this POI family, we generated a knock-in mouse *Hsf2bp^S167L/S167L^* by genome editing (Figure S5a). We also generated a loss of function model (*Hsf2bp^-/-^*) for direct comparison (Figure S5b-e). *Hsf2bp^S167L/S167L^* male and female mice were able to reproduce but females showed a significant reduction in the number and size of litters (Figure 1a) whilst males only showed a slight non-significant reduction in fertility (Figure 1a). This is in agreement with the fertility defects observed in *Hsf2bp^-/-^* mice (Figure 1c, e-f and (Brandsma et al., 2019; Zhang et al., 2019)), and suggests that the S167L variant might impact more specifically female fertility.

**Figure 1.**
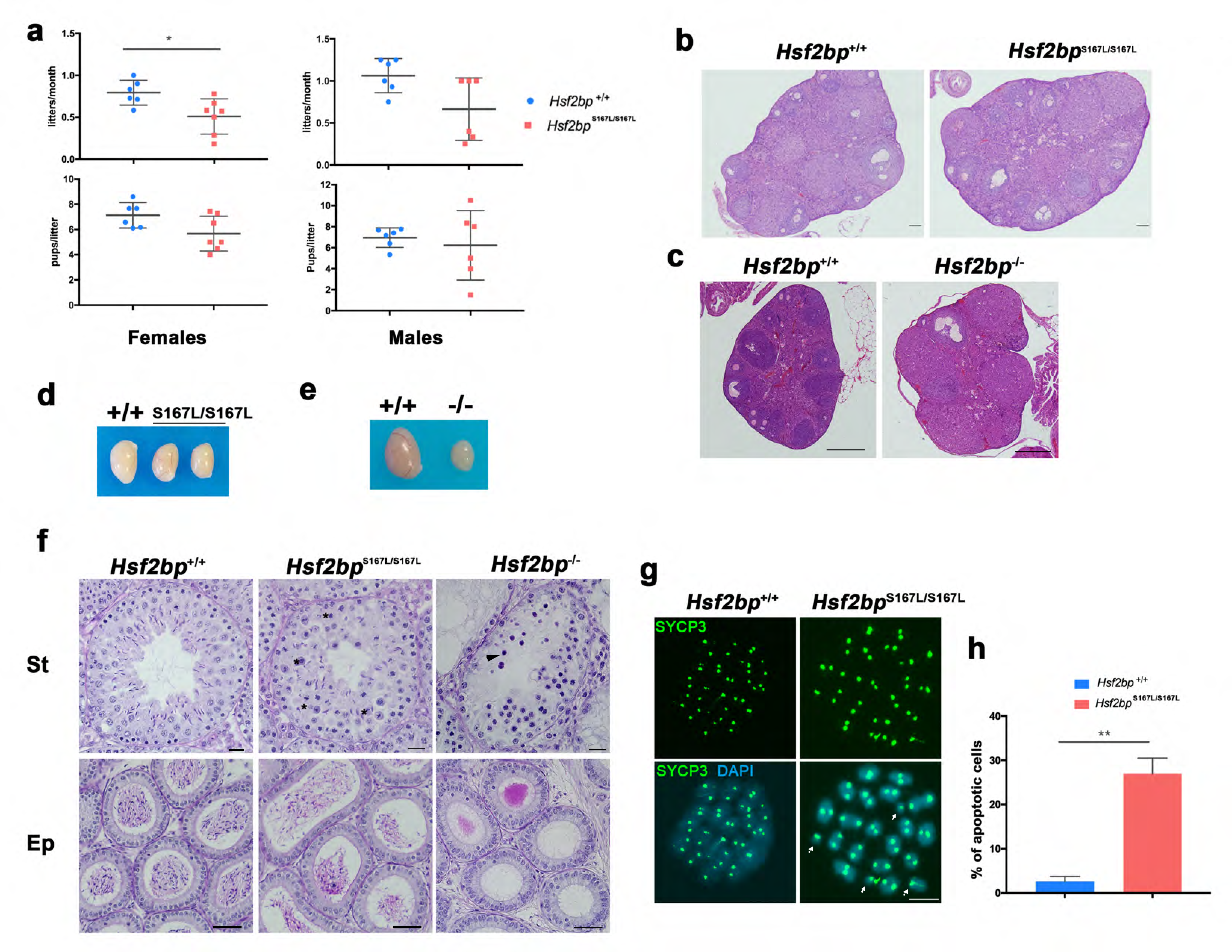
Mice carrying the HSF2BP S167L variant show a partial reduction of fertility. (a) Fertility assessment of *Hsf2bp*^S167L/S167L^ mice were performed by crossing *Hsf2bp*^+/+^ and *Hsf2bp*^S167L/S167L^ males and females (8 weeks old) with wild type females and males, respectively. The number of pups per litter was recorded. Both mutant males (right plots) and females (left plots) were able to reproduce but females showed a reduction in fertility whereas males only show a trend towards a reduction of the fertility. Welch’s t-test analysis: *p<0.05; (b) Ovaries from adult (8 weeks) *Hsf2bp*^S167L/S167L^ females looked apparently normal and do not show a significant reduction in the number of follicles as shown in hematoxylin+eosin stained sections. Bar in panels 50μm. (c) Ovaries from adult (8 weeks) *Hsf2bp*^-/-^ females show a strong depletion of follicles that leads to a premature ovarian insufficiency (POI). Bar in panels 200μm. (d-e) Testis size of *Hsf2bp*^S167L/S167L^ (d, 25% reduction) and *Hsf2bp*^-/-^ mice (e, 75% reduction) in comparison with their WT counterparts. (f) The S167L variant in HSF2BP leads to a partial arrest of spermatogenesis with all the cell types present but an elevated number of apoptotic meiotic divisions (asterisks) as shown in PAS+hematoxylin stained testis sections. The spermatogenic partial arrest leads to a reduction in the number of spermatozoa in the epididymides. The *Hsf2bp*^-/-^ showed a more exacerbated phenotype with a complete arrest of spermatogenesis in epithelial stage IV and a total absence of spermatozoa. Massive apoptosis of spermatocytes is indicated (black arrowheads). Bar in upper panels 10μm and in lower panels 20μm. (St) Seminiferous tubules, (Ep) Epididymides. (g) Labeling of spermatocyte spread preparations with SYCP3 (green), showing the presence of univalent chromosomes (arrows) in metaphases I (MI) of *Hsf2bp*^S167L/S167L^ males. Bar in panel, 10 μm. (h) Quantification of the percentage of apoptotic cells referred to the total number of spermatocytes in spread preparations from *Hsf2bp*^-/-^ and *Hsf2bp*^S167L/S167L^ mice. Welch’s t-test analysis: ** p<0.01.

Histological analysis of *Hsf2bp^S167L/S167L^* ovaries revealed no apparent difference in the number of follicles in comparison with WT animals, in contrast with the drastic reduction of the follicle pool in *Hsf2bp^-/-^* ovaries (Figure 1b-c). Testes from *Hsf2bp^S167L/S167L^* mice displayed a reduced size (only 70% of WT mice: S167L 67,1mg ±10,4 vs 95,6 mg ±10,1 for WT controls; n=6, Figure 1d) and this reduction was higher in *Hsf2bp^-/-^* testes (25% of the wild-type, Figure 1e). Accordingly, histological analysis of adult *Hsf2bp^S167L/S167L^* testes revealed seminiferous tubules with a partial arrest with apoptotic spermatocytes (meiotic divisions) and their epididymis exhibited scarcer spermatozoa (Figure 1f). *Hsf2bp^-/-^* males show a total meiotic arrest at epithelial stage IV with massive apoptosis (Figure 1f). The presence of spermatogonia, spermatocytes, Sertoli and Leydig cells was not altered in any of the mutants (Figure 1f). These results suggest that mice bearing the POI-causing variant only partially phenocopy the human disease and fits well with the exacerbated sexual phenotypic dimorphism observed in the *Hsf2bp^-/-^* mice which might be less pronounced or absent in humans. However, the existence of a mild subfertility in females and of an overt spermatogenic phenotype prompted us to analyse their meiotic progression in more detail.

To characterize the meiotic defect in detail, *Hsf2bp^S167L/S167L^* meiocytes were first analyzed for the assembly/disassembly of the SC by monitoring the distribution of SYCP1 and SYCP3. We did not observe any difference in synapsis and desynapsis from leptotene to diplotene in both oocytes and spermatocytes (Figure S6a-b). However, we observed an elevated number of apoptotic meiotic divisions in *Hsf2bp^S167L/S167L^* males (30% in *Hsf2bp^S167L/S167L^* vs 5% in WT) as well as some metaphases I with free univalents (Figure 1g-h and S7). These results are consistent with the partial arrest observed in the histological analysis (Figure 1f). As expected, this phenotype was exacerbated in *Hsf2bp^-/-^* spermatocytes that were arrested at a zygotene-like stage (Figure S6c) and in the *Hsf2bp*-deficient oocytes that showed a delay in prophase I progression with the majority of cells at zygotene stage in 17,5 dpp females (whilst the WT oocytes are mainly at pachytene stage). Additionally, we observed increased numbers of oocytes showing synapsis defects in the *Hsf2bp^-/-^* oocytes (45,5%±1,5 vs 7,5%±1,5 in WT, Figure S6d). These results strongly suggest that the POI variant S167L is a hypomorphic allele.

We next analyzed if the POI-inducing variant affects the loading/stability of HSF2BP by immunolabelling meiocytes from *Hsf2bp^S167L/S167L^* mice. We observed a striking reduction of HSF2BP staining at the axes during prophase I in both oocytes and spermatocytes (Figure 2a-b). This reduction and the specificity of the antibodies used were validated by Western blot of whole testis extracts from wild-type, *Hsf2bp*^S167L/S167L^ and *Hsf2bp^-/-^* animals (Figure 2c) and by immunofluorescence (IF) of *Hsf2bp^-/-^* spermatocytes (Figure S5e).

**Figure 2.**
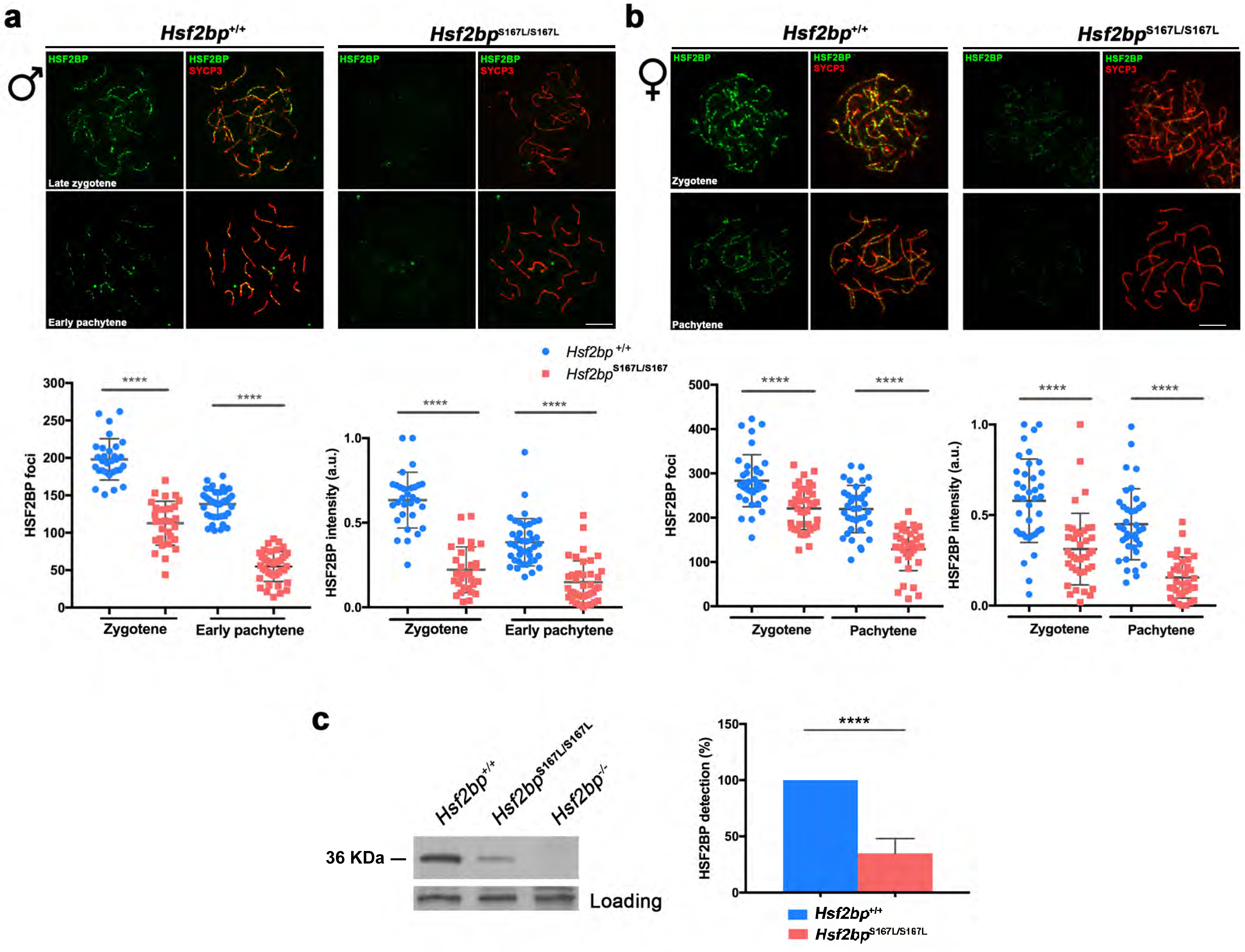
Meiocytes from *Hsf2bp*^S167L/S167L^ mice show a decrease in the expression of HSF2BP. (a-b) Double immunofluorescence of HSF2BP (green) and SYCP3 (red) in *Hsf2bp^+/+^*and *Hsf2bp*^S167L/S167L^ (a) spermatocyte and (b) oocyte spreads showing a strong reduction in the labelling of HSF2BP at the chromosome axis. Plots under the panels show the quantification of the number of HSF2BP foci and intensity on each genotype. Bar in panels, 10 μm. (c) Western blot analysis of protein extracts from adult wild type, *Hsf2bp*^S167L/S167L^ and *Hsf2bp*^-/-^ testes with a specific antibody against the whole recombinant HSF2BP protein. An unspecific band was used as loading control. Graph on the right represents the quantification of the immunoblotting detection of HSF2BP in *Hsf2bp*^S167L/S167L^ testis extracts in comparison with WT (100%). Welch’s t-test analysis: **** p<0.0001.

Given that HSF2BP is essential for DNA repair in spermatocytes and that its deficiency provokes the accumulation of DNA repair proteins such as H2AX (‘tagging’ the DSBs), the single stranded-DNA binding protein RPA, a strong reduction of the recombinases DMC1 and RAD51 at the RNs, and the lack of cross-overs (COs) by labelling MLH1 (Baker et al., 1996) (Figures S8, 3, 4 and 5), we carried out a comparative analysis of these proteins in meiocytes from *Hsf2bp^S167L/S167L^, Hsf2bp^-/-^* and WT animals. Our results reveal that *Hsf2bp^S167L/S167L^* spermatocytes also showed an increased labelling of H2AX at pachytene (Figure 3a), an accumulation of RPA at the chromosome axis (Figure 3b), a reduction of the recombinases DMC1 and RAD51 staining (Figure 4a-b), and a decreased number of COs (measured as MLH1 or interstitial CDK2 foci, Figure 5a) in comparison with the WT. Consequently with the reduction of COs, we observed the presence of free XY univalents at pachytene in *Hsf2bp^S167L/S167L^* males (Figure 5a), which explains the univalents observed at metaphase I and the elevated number of apoptotic cells observed (Figure 1g-h and S7). However, our female analysis showed no accumulation in RPA labelling in *Hsf2bp^S167L/S167L^* oocytes (Figure 5b), whereas RPA accumulates in the synapsis-defective cells from KO oocytes (Figure S8b) and DMC1 shows lower staining in both *Hsf2bp^S167L/S167L^* and *Hsf2bp^-/-^* (Figures 5c and S8c). MLH1 foci show a strong reduction and no differences (though with a clear trend towards a reduction) in the number of COs in oocytes from *Hsf2bp^-/-^ and Hsf2bp^S167L/S167L^,* respectively (Figure 5d). Overall, male and female *Hsf2bp^S167L/S167L^* mice show similar molecular alterations although with different severity. In humans, and given the absence of men with the homozygous variant in the POI family (Figure S1), a similar difference between the sexes cannot be ruled out.

**Figure 3.**
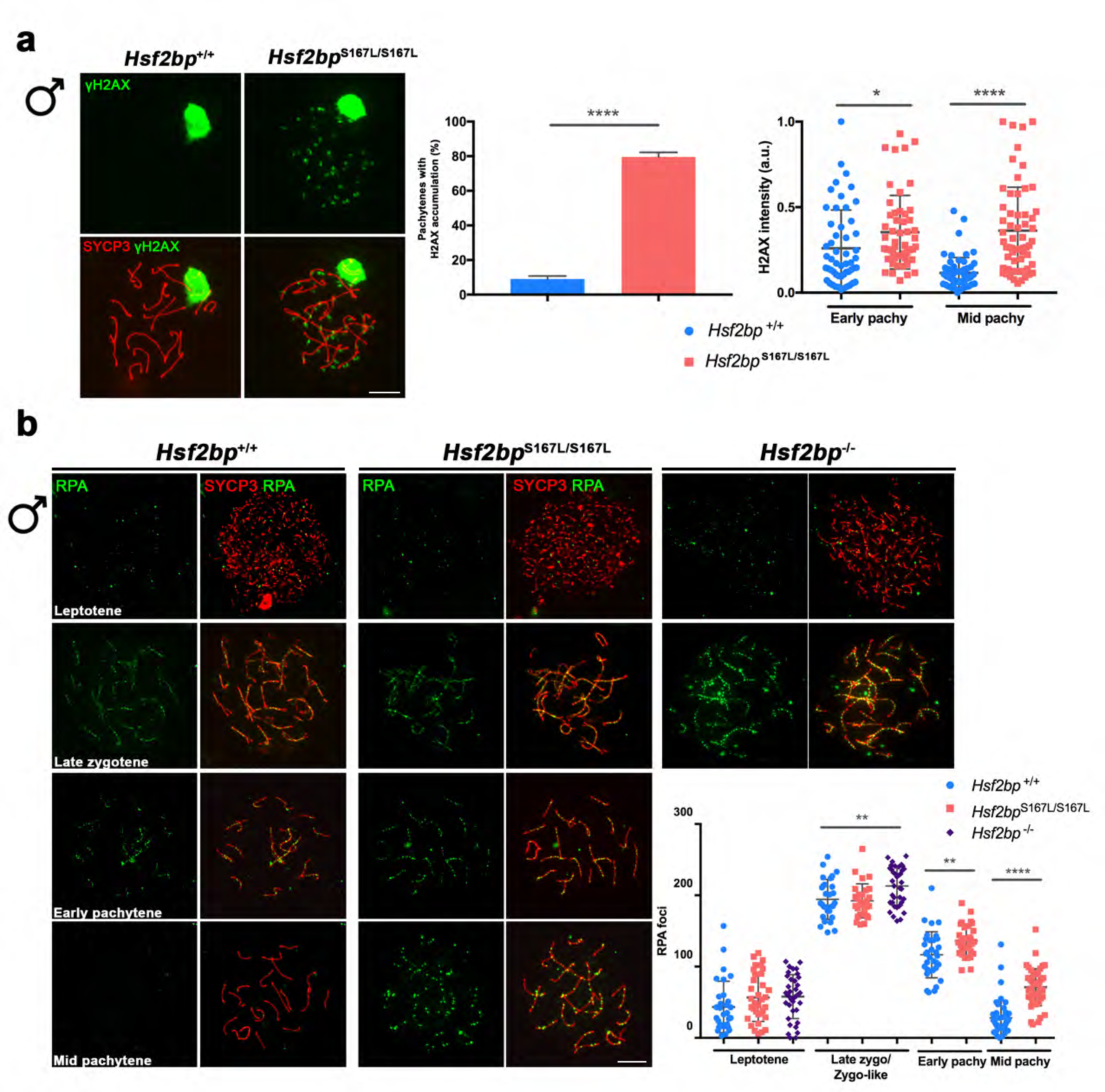
*Hsf2bp* ^S167L/S167L^ males show defects in DNA repair. (a) Double labeling of γH2AX (green) and SYCP3 (red) in spermatocyte spreads from WT and S167L mice showing the accumulation of γH2AX patches in the mutant pachynemas. Plots on the right of the panel represent the percentage of pachynemas that showed γH2AX accumulation and the quantification of γH2AX intensity on autosomes at early and mid pachytene. (b) Double immunolabeling of RPA (green) and SYCP3 (red) in spermatocyte spreads from *Hsf2bp^+/+^*, *Hsf2bp* ^S167L/S167L^ and *Hsf2bp^-/-^*. RPA accumulates at early and mid pachytene in S167L spermatocytes and in the zygotene-like arrested cells from *Hsf2bp^-/-^*. Welch’s t-test analysis: no asterisks, no significant differences; *p<0.05, ** p<0.01, **** p<0.0001. Bar in panels, 10 μm.

**Figure 4.**
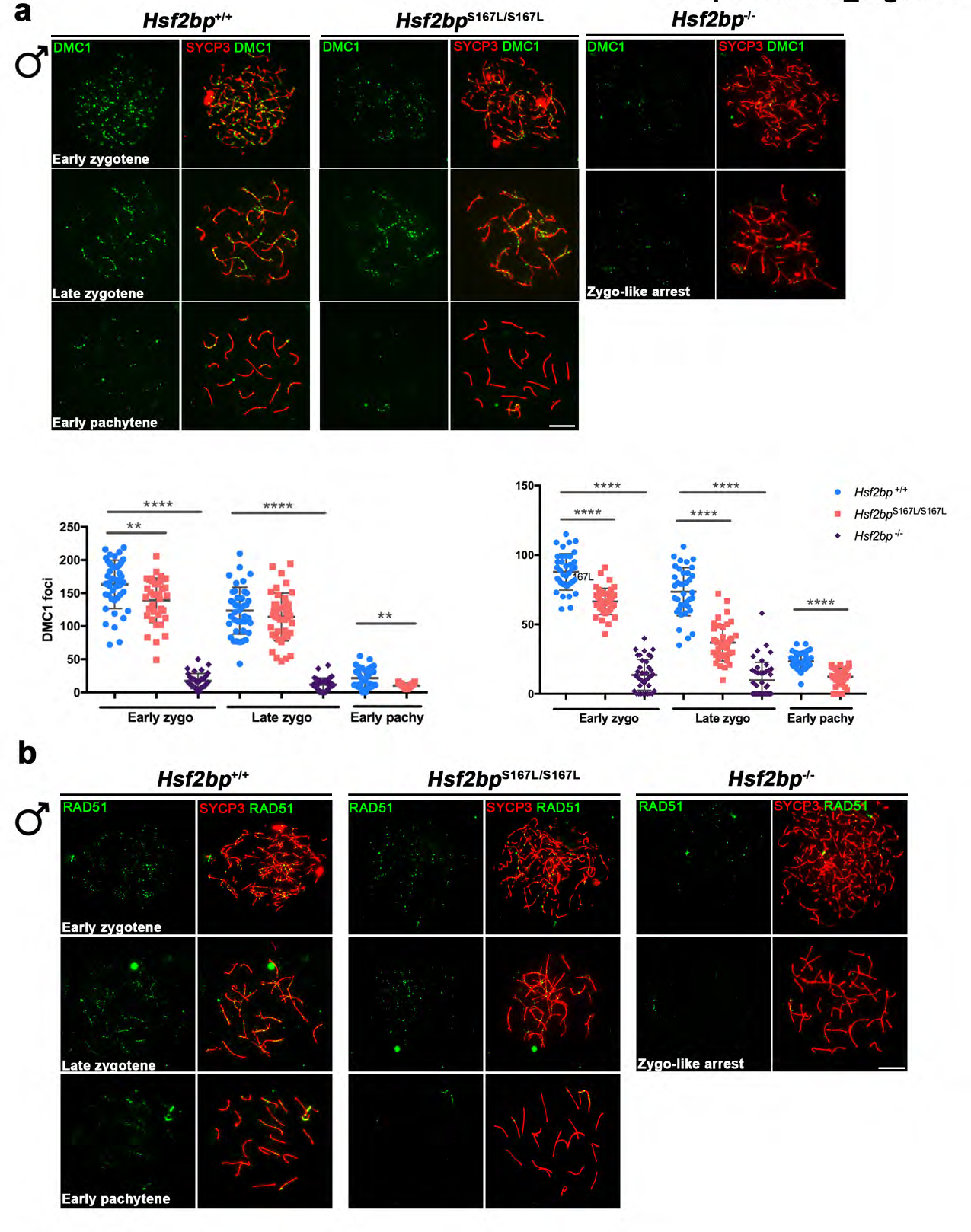
The loading of recombinases is compromised in *Hsf2bp*^S167L/S167L^ males. (a) Double immunolabeling of DMC1 (green) and SYCP3 (red) in *Hsf2bp^+/+^*, *Hsf2bp* ^S167L/S167L^ and *Hsf2bp^-/-^* spermatocytes showing a strong reduction (*Hsf2bp^-/-^*) and mild reduction (*Hsf2bp* ^S167L/S167L^) in comparison with their WT counterparts. Plot under the panel represents the quantification of DMC1 foci on each genotype and stage. (b) Double immunolabeling of RAD51 (green) and SYCP3 (red) in *Hsf2bp^+/+^*, *Hsf2bp* ^S167L/S167L^ and *Hsf2bp^-/-^* spermatocytes. RAD51 staining is reduced in the knock-out and S167L mutants in comparison with the wild-type. Plot above the panel represents the quantification of RAD51 foci on each genotype and stage. Bar in panels, 10 μm. Welch’s t-test analysis: ** p<0.01; **** p<0.0001.

**Figure 5.**
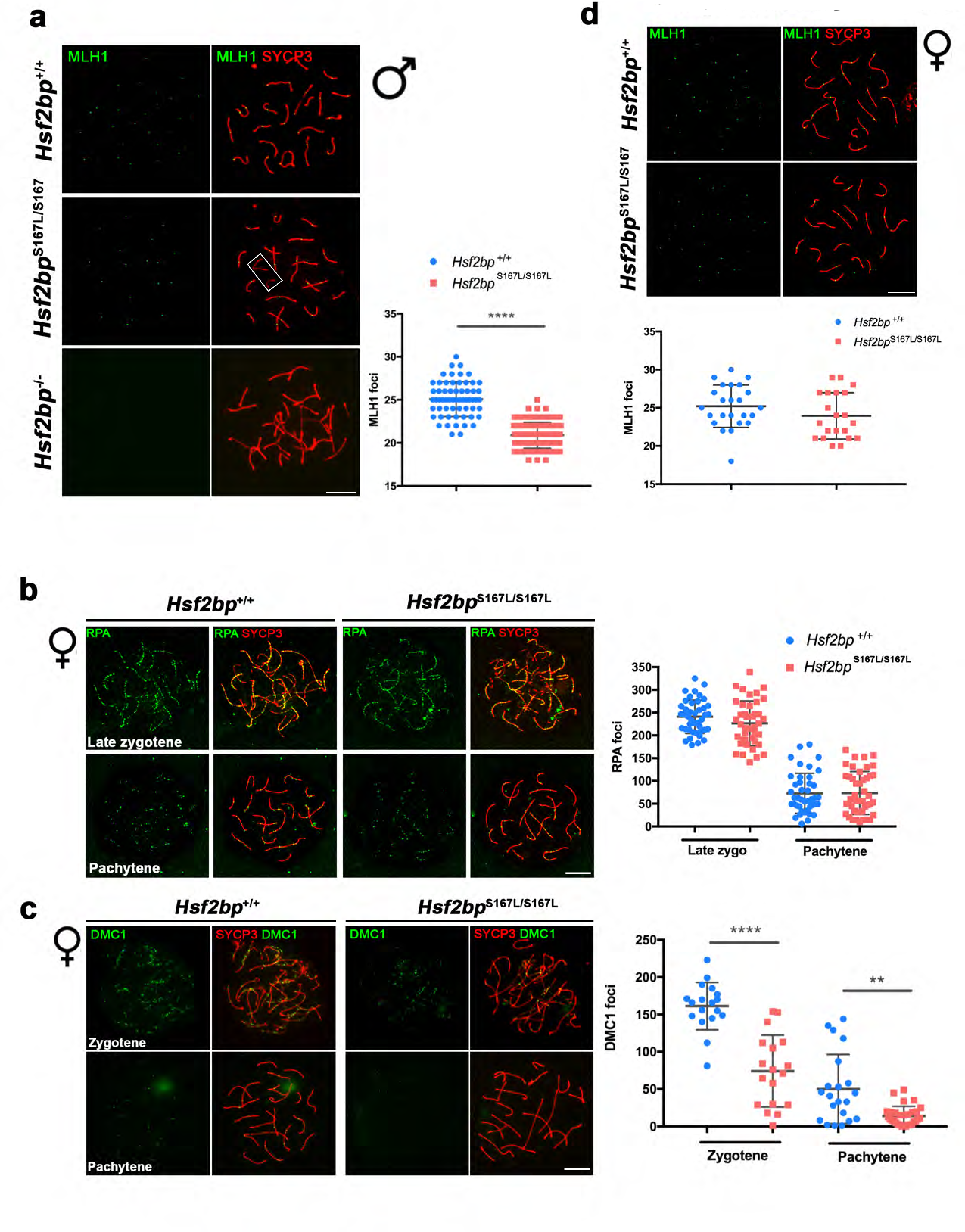
Recombination is clearly affected in *Hsf2bp*^S167L/S167L^ males and partially affected in females. (a) Double immunofluorescence of MLH1 (green) and SYCP3 (red) in spermatocyte spreads from *Hsf2bp*^S167L/S167L^ and *Hsf2bp^+/+^*. MLH1 foci are significantly reduced in the *Hsf2bp*^S167L/S167L^ spermatocytes and showed XY univalents at pachytene stage (white rectangle, see also figure 1g). The plot on the right shows the quantification of MLH1 foci in both genotypes. (b) Double immunolabeling of RPA (green) and SYCP3 (red) in oocyte spreads from *Hsf2bp^+/+^* and *Hsf2bp* ^S167L/S167L^ females showing normal RPA localization and number of foci. Plot on the right of the panel represents the quantification of RPA foci on each genotype and stage. (c) Double immunofluorescence of DMC1 (green) and SYCP3 (red) in *Hsf2bp^+/+^* and *Hsf2bp* ^S167L/S167L^ oocytes showing a reduction in the number of DMC1 foci. Plot on the right of the panel represents the quantification of DMC1 foci on each genotype and stage. (d) Double immunolabeling of MLH1 (green) and SYCP3 (red) in oocyte spreads from *Hsf2bp^+/+^* and *Hsf2bp* ^S167L/S167L^ mice (19.5 dpp) showing no significant differences between both genotypes in the number of cross-overs (see plot under the panel for quantification). Bar in panels, 10 μm. Welch’s t-test analysis: ** p<0.01, **** p<0.0001.

We next sought to understand how the HSF2BP pathogenic variant is mediating the observed meiotic alteration. HSF2BP has been shown to bind BRCA2, an essential protein for homologous meiotic recombination (Martinez et al., 2016; Sharan et al., 2004), in a direct manner tnatahrough its tryptophane at position 200 and the C-term (Gly2270-Thr2337) (Brandsma PhD thesis 2016; (Brandsma et al., 2019)). Given the impossibility to detect endogenous BRCA2 by IF in mouse spermatocytes, we carried out co-localization/interaction assays in a heterologous system by transfecting BRCA2-C (C-term) and HSF2BP in U2OS/HEK293T. The results showed that BRCA2-C co-immunoprecipitates with both HSF2BP-WT and HSF2BP-S167L in similar ways (Figure S9a). When transfected alone, HSF2BP localization was cytoplasmic and BRCA2-C was nuclear. This pattern changed drastically to a nuclear dotted pattern when co-transfected (Figure S9b). This re-localization is independent of the HSF2BP variant, strongly suggesting that the HSF2BP variant effects are not directly mediated by BRCA2 delocalization.

### MIDAP is a novel interactor of HSF2BP

In order to further understand the mechanism underlying the pathogenicity of the HSF2BP-S167L variant, we searched for proteins that interact with the murine HSF2BP through a yeast two hybrid (Y2H) screening. The analysis of the clones with putative interactors revealed that 19 contained an uncharacterized protein named C19ORF57, herein renamed MIDAP for meiotic double-stranded break BRCA2/HSF2BP complex associated protein. This interactor consists of 600 amino acids with a high content of acidic residues, has no recognizable functional domains and is intrinsically disordered. The interaction was validated by transiently transfecting HSF2BP-and MIDAP-encoding expression plasmids. Both HSF2BP-S167L and WT interacted with MIDAP (Figure 6a). To identify the regions required for this interaction, we splitted the MIDAP protein into three fragments (N-terminal, central region and C-terminal). We mapped the HSF2BP/MIDAP-interacting domain to the C-term fragment of MIDAP (spanning residues 475-600 of the murine protein, Figure 6b-c).

**Figure 6.**
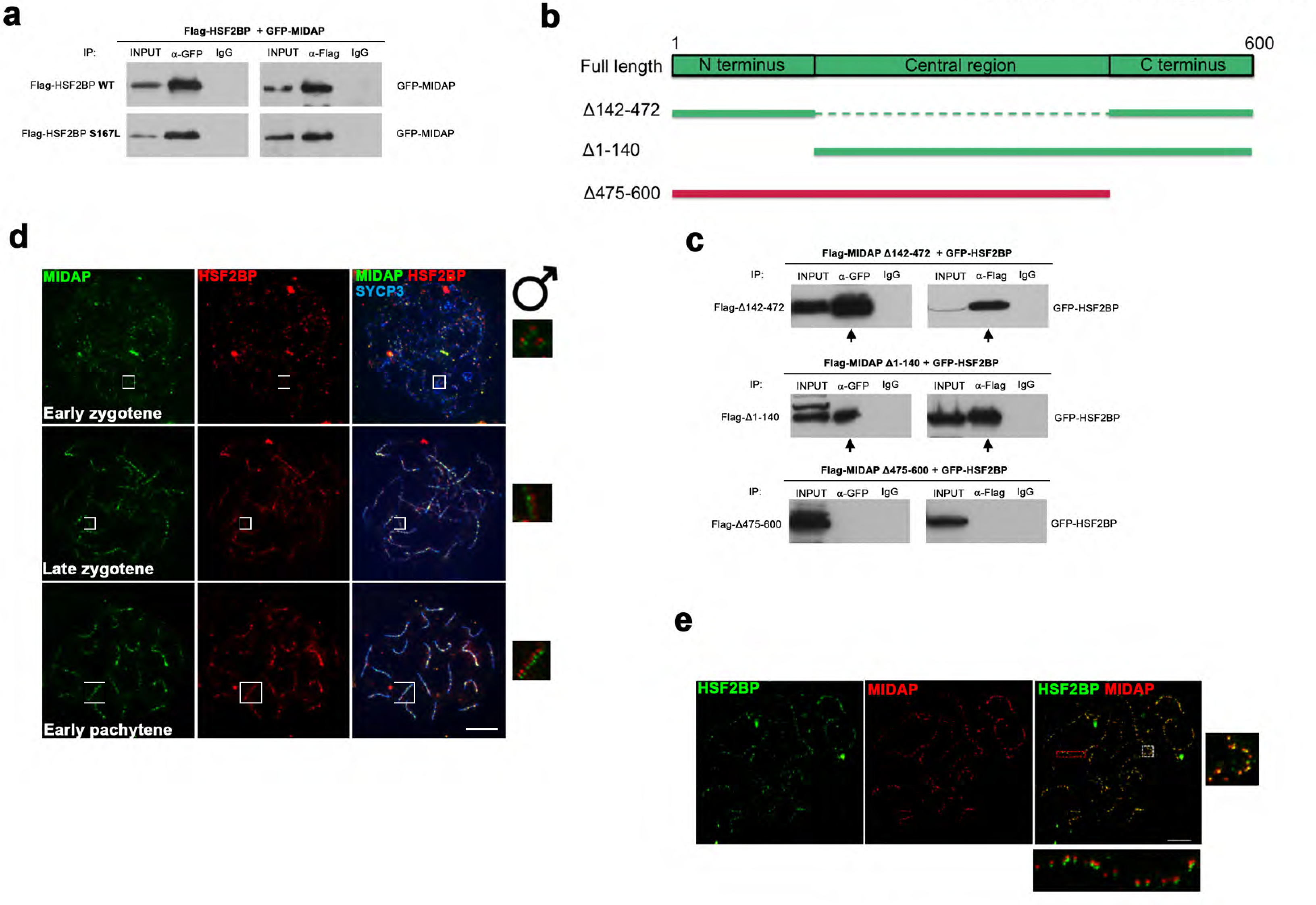
Analysis of the HSF2BP binding region of MIDAP and *in vivo* colocalization of HSF2BP and MIDAP to RNs. (a) HEK293T cells were transfected with Flag-HSF2BP (WT, upper panel; S167L, lower panel) and its novel interactor GFP-MIDAP. Protein complexes were immunoprecipitated overnight with either an anti-Flag or anti-EGFP or IgGs (negative control), and were analysed by immunoblotting with the indicated antibody. Both HSF2BP variants (WT and S167L) co-immunorecipitate (co-IP) with MIDAP similarly. (b) Schematic representation of full-length MIDAP protein and the corresponding Delta (Δ) constructs (filled boxes) generated and subcloned in expression vectors to decipher the essential MIDAP region for interacting with HSF2BP. Constructs in green indicate positive interaction, the construct in red indicates no interaction. (c) HEK293T cells were transfected with GFP-HSF2BP and the different delta constructs of Flag-MIDAP. Immunoprecipitation and analysis were done as in Figure 6a. The Δ475-600 abolishes the interaction indicating that the C terminus of MIDAP is the essential region of interaction with HSF2BP. (d) Triple immunofluorescence of MIDAP (green), HSF2BP (red) and SYCP3 (blue) in wild type spermatocyte spreads showing a perfect colocalization between MIDAP and HSF2BP at late zygotene and pachytene. Bar in panel, 10μm. (e) Double immunolabeling of spermatocyte spread preparations with HSF2BP (green) and MIDAP (red) by Stimulated emission depletion (STED) microscopy. Bar in panel, 5μm.

Next, we asked whether this interacting domain was sufficient for HSF2BP function *in vivo*. We generated the corresponding mutant mice (*Midap^Δ/Δ^*), which expressed the MIDAPΔ142-472 (Figure S10a-d). *Midap^Δ/Δ^* mice loaded HSF2BP to the axes similar to the WT, did not show defects in chromosome synapsis and were fertile (Figure S10e-f). These results indicate that a large fraction of the coding protein of MIDAP is not essential for HSF2BP function *in vivo*.

We next sought to analyse its function in meiosis through IF by using specific antibodies against MIDAP. MIDAP localized on the chromosome axes of meiocytes from zygotene to pachytene with a pattern of discrete foci that mimics RNs (Figure S11a-b). In agreement with the Y2H and co-IP results, MIDAP perfectly co-localized with HSF2BP on the chromosome axes (Figure 6d). This co-localization was verified by super-resolution microscopy (Figure 6e). To further delineate if HSF2BP and/or MIDAP had DNA-binding activity (targeting to DSBs), we carried out an *in vitro* binding assay using HSF2BP and MIDAP proteins expressed in a transcription and translation coupled reticulocyte system (TNT) in which there are no nuclear protein and chromatin (Melton et al., 1984) and used RPA as positive control. The results show that both proteins lack DNA-binding abilities in contrast to the strong activity of RPA (Figure S11c-d).

To determine the role of MIDAP in recombination and DNA repair, we analyzed its cytological pattern of distribution in different synaptic/recombination mutants (REC8, SIX6OS1, RNF212, HEI10 and PSMA8 (from more severe to less severe phenotypes) and in DSBs-deficient spermatocytes (SPO11 mutant; (*Spo11^-/-^, Rnf212^-/-^* and *Hei10^-/-^* described in this work, see methods and Figures S12 and S13)). HSF2BP staining was also carried out for a direct comparison. The results showed that none of the recombination-deficient mutants abrogate MIDAP labelling at zygotene, in contrast to its absence of loading in SPO11-deficient mice (Figure S14, left). These results are very similar to those obtained for HSF2BP in these mutants (Figure S14, right) and indicate that SPO11-dependent DSBs are essential for targeting HSF2BP/MIDAP to the RNs and both can be genetically positioned at the early events very soon after DSBs generation.

To functionally analyse the role of MIDAP in HSF2BP signalling and fertility, we generated a null mutant by genome editing. *Midap^-/-^* mice displayed an absence of protein signal in Western blot and IF of chromosome spreads (Figure S15). Females, despite being fertile, showed a strong reduction of the follicle pool (Figure 7a). Male *Midap^-/-^* mice were infertile, the average size of their testes were severely reduced (WT 89,2 mg ±7,72 vs KO 30,2mg ±4,77; 66% reduction; n=4), and lacked spermatozoa (Figure 7b-c). Histological analysis showed a meiotic arrest at epithelial stage IV with apoptotic spermatocytes (Figure 7c). Double immunolabeling of SYCP3 and SYCP1 revealed that spermatocytes were partially synapsed but showed a partner-switch phenotype in which synapsis is not restricted to homologous pairs (Figure 7d). The arrest corresponds to a zygotene-like stage. *Midap^-/-^* oocyte spreads analysis revealed the presence of a subset of fully-synapsed pachynemas but an increased number of cells with different synaptic defects (47,9%±1,6 vs 12,2%±4,1 in the WT, Figure 7e). Given the relationship between HSF2BP and MIDAP, we asked whether HSF2BP localization depends on MIDAP by immunolabeling of HSF2BP in *Midap^-/-^* spermatocytes and oocytes. Our results showed an almost absence of HSF2BP staining in MIDAP-null meiocytes (Figure 8a-b).

**Figure 7.**
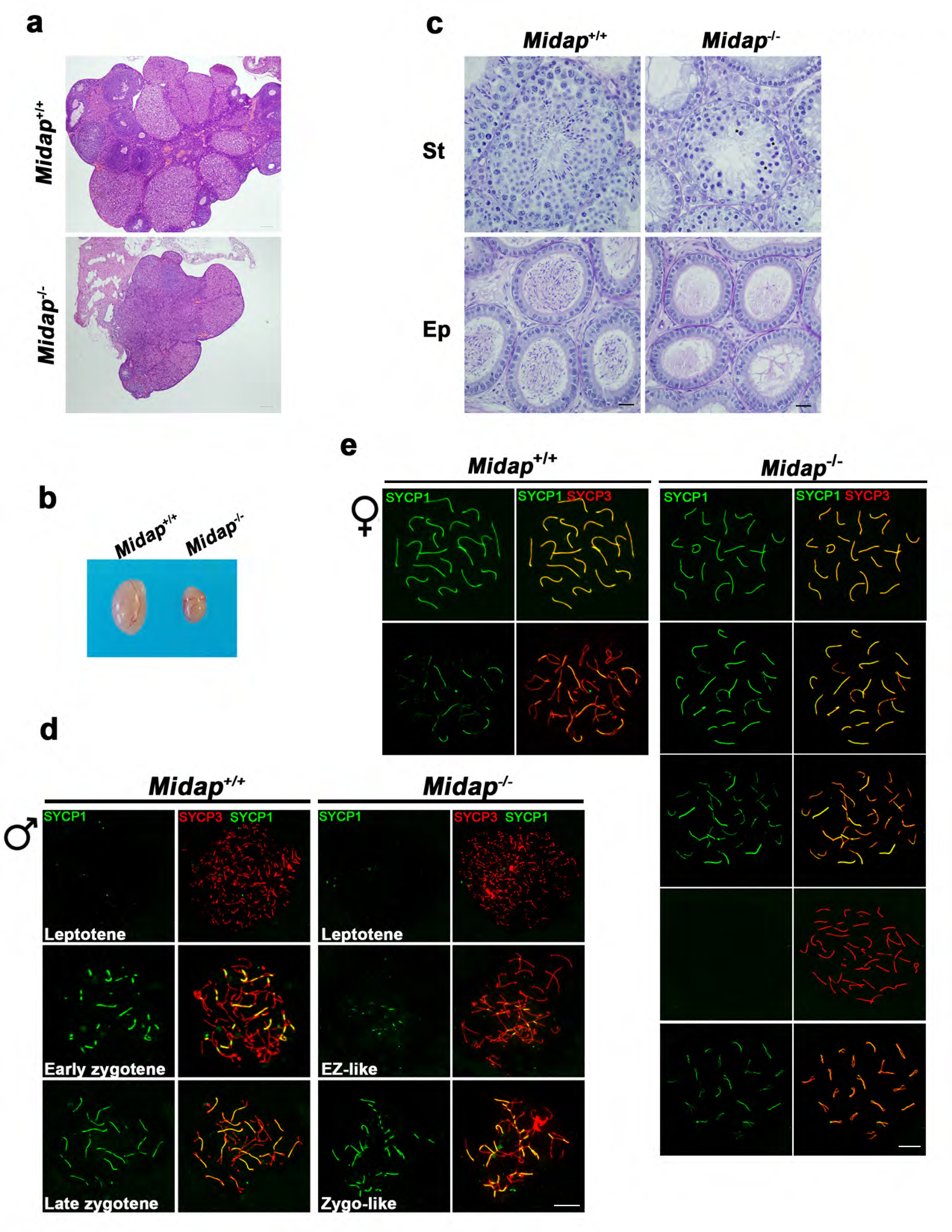
MIDAP-deficient mice show ovaries with fewer follicles and testis without spermatozoa. (a) Ovaries from adult *Midap^-/-^* females show a strong but not total depletion of follicles. Bar in panels, 50μm. (b) Testes from adult *Midap^-/-^* males show a reduction of the testis size and (c) a complete arrest of spermatogenesis in epithelial stage IV as shown in PAS+hematoxylin stained testis sections. Massive apoptosis of spermatocytes is indicated (asterisks). The spermatogenic arrest leads to empty epididymides and non-obstructive azoospermia. (**St**) Seminiferous tubules. (**Ep**) Epididymides. Bar in panels, 10μm. (d-e) Double labelling of (d) spermatocyte and (e) oocyte spreads from wild-type and *Midap^-/-^* mice with SYCP3 (red) and SYCP1 (green). *Midap^-/-^* spermatocytes showed an arrest in a zygotene-like stage establishing synapsis between non-homologous chromosomes (partner switch phenotype). (e) *Midap^-/-^* females showed a subset of fully-synapsed pachynemas (17,5 dpp) but increased numbers of different synapsis-defective cells. Bar in panels, 10μm.

**Figure 8.**
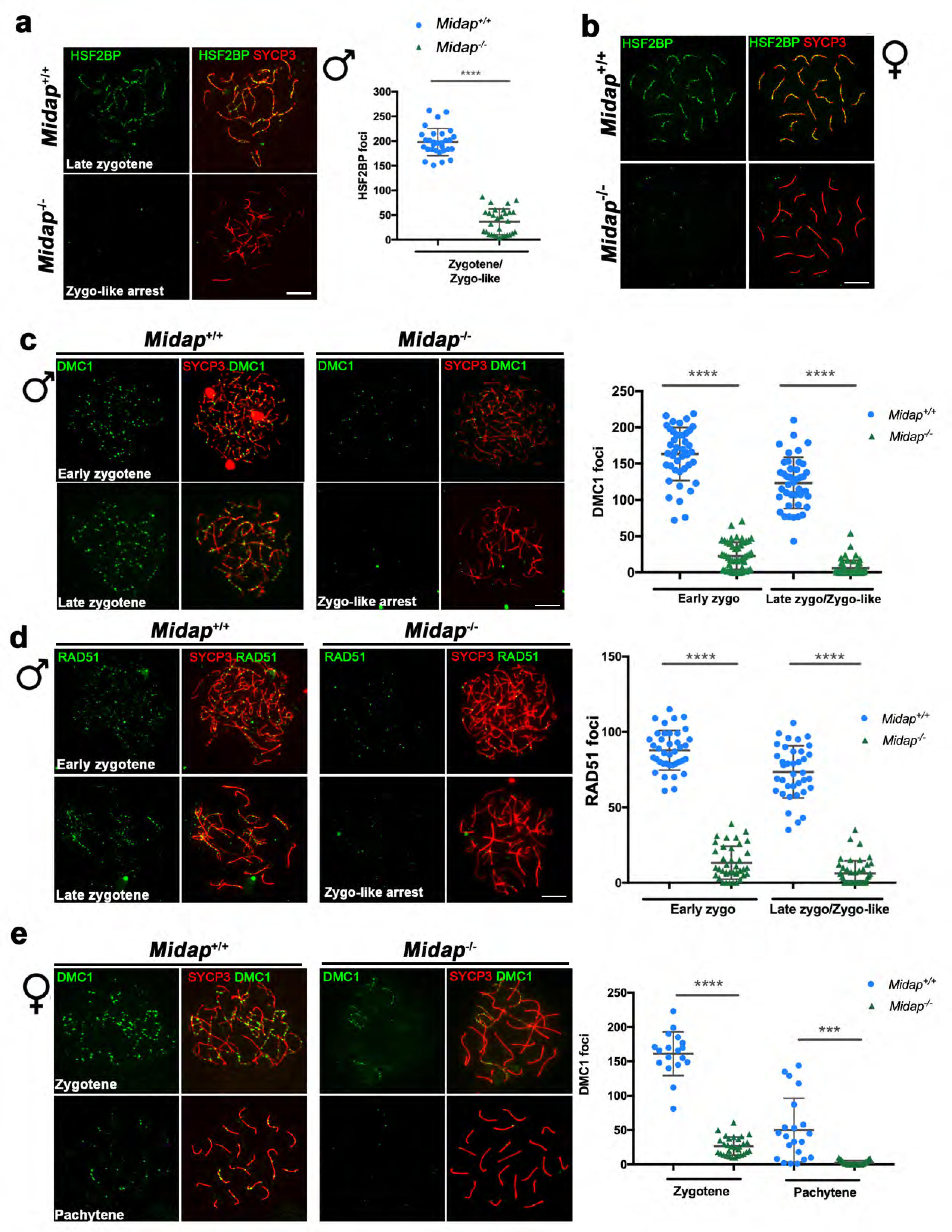
HSF2BP and recombinases loading is dependent of MIDAP. (a-b) Double labelling with HSF2BP (green) and SYCP3 (red) of (a) spermatocyte and (b) oocyte spreads showing faint HSF2BP labelling in spermatocytes and absence of labelling in oocytes. Plot on the right of (a) panel represents de quantification of MIDAP foci in *Midap^-/-^* spermatocytes. (c-d) Double immunofluorescence of (c) DMC1 or (d) RAD51 (both in green) and SYCP3 (red) in *Midap^+/+^* and *Midap^-/-^* spermatocytes showing a strongly reduced labelling in the *Midap^-/-^* in comparison to the WT. Plot right to the panels represents the quantification of DMC1 and RAD51 foci on each genotype and stage. (e) Double labelling of DMC1 (green) and SYCP3 (red) on oocyte spreads from *Midap^+/+^* and *Midap^-/-^* females also showing a strong reduction in the MIDAP-null oocytes. Plot right to the panel shows the quantification of DMC1 foci in zygotene and pachytene oocytes. The quantification of HSF2BP, DMC1 and RAD51 foci was done in parallel in the WT, *Hsf2bp*^S167L/S167L^, *Hsf2bp*^-/-^ and *Midap^-/-^* mice. For this reason, the data for the WT are the same on this figure and the figures 2a, 4b, 5a, 5c and S8c). Bar in panels, 10μm. Welch’s t-test analysis: *** p<0.001, **** p<0.0001.

Immunostaining of *Midap^-/-^* spermatocytes for H2AX, RPA and the recombinases RAD51 and DMC1 revealed an accumulation of H2AX and RPA on zygonema-like spermatocytes and a drastic reduction of RAD51/DMC1 foci in early and late zygonema (Figure S16a-b and Figure 8c-d). Accordingly with the arrest, MLH1 staining revealed absence of COs (Figure S17a). These results are similar to the phenotypes described for HSF2BP mutants (Figures 3, 4, 5, S8 and (Brandsma et al., 2019)). We also analyzed the localization of SPATA22, another ssDNA binding protein complexed to RPA during resection, and found a strong accumulation in both HSF2BP and MIDAP null mutants and milder staining in the *Hsf2bp^S167L/S167L^* spermatocytes supporting early meiotic defects before strand invasion (Figure S18). Similar to the HSF2BP, oocytes from *Midap^-/-^* females show a reduced staining of DMC1 and consequently a reduced number of MLH1 foci (Figure 8e and S17b). Thus, both HSF2BP and MIDAP mouse mutant show a meiotic phenotype with a strong sexual dimorphism in the mouse.

In order to study HSF2BP signalling through MIDAP, we analyzed its localization in *Hsf2bp^-/-^* and *Hsf2bp^S167L/S167L^* spermatocytes and oocytes (Figure 9a-b). Our results showed a total absence of MIDAP loading in *Hsf2bp^-/-^* meiocytes and a drastic reduction as a consequence of the POI variant (Figure 9a-b), strongly suggesting a causative relationship.

**Figure 9.**
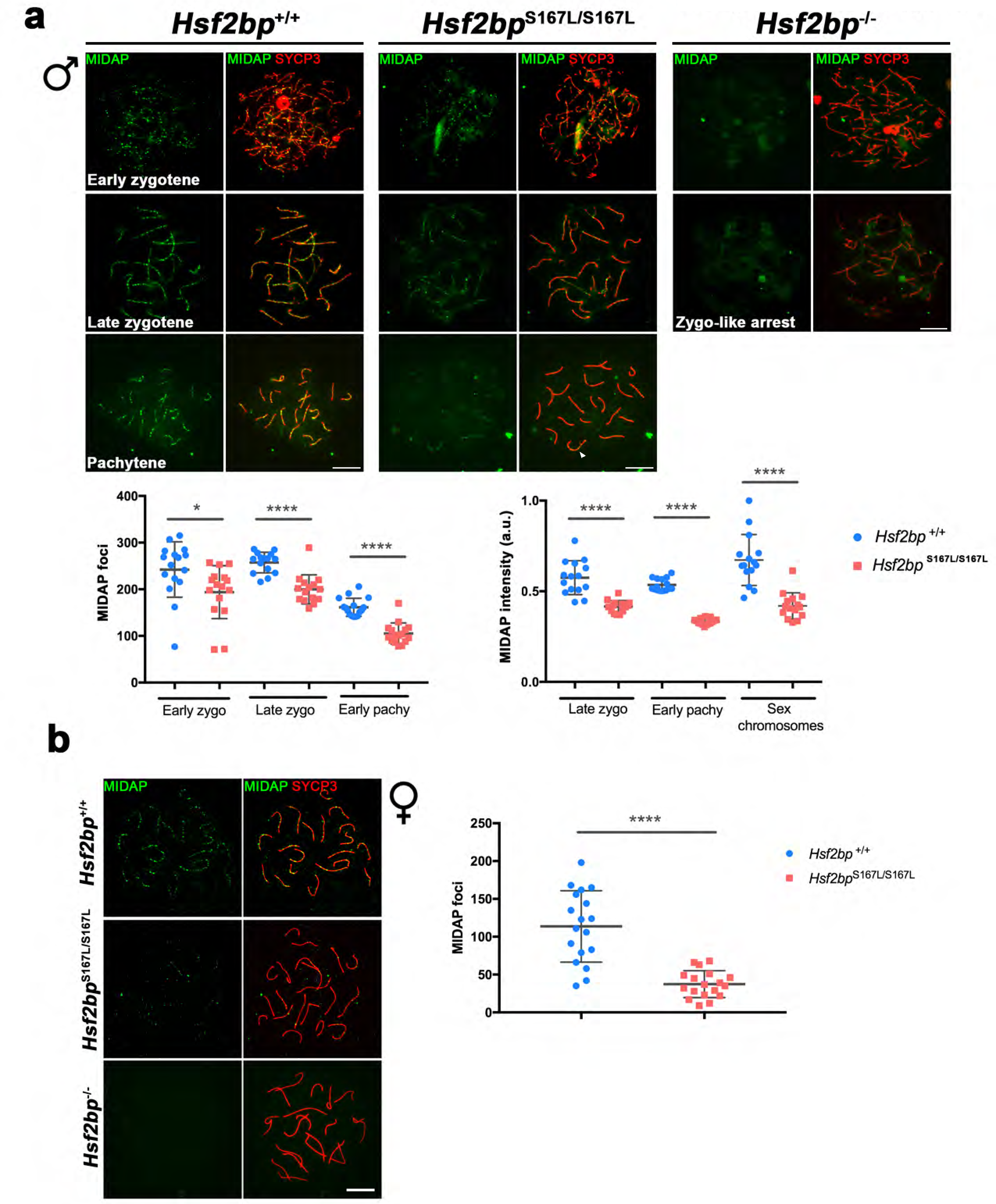
MIDAP loading is also dependent on HSF2BP. (a-b) Double immunofluorescence of MIDAP (green) and SYCP3 (red) in (a) spermatocytes and (b) oocytes from *Hsf2bp*^+/+^, *Hsf2bp*^S167L/S167L^ and *Hsf2bp*^-/-^ showing a strong reduction of MIDAP staining in the S167L mutant and a total absence in the *Hsf2bp* knock-out. Plots under the panel represent the quantification of the number of MIDAP foci or intensity on each genotype and stage. Bar in panels, 10μm. Welch’s t-test analysis: * p<0.05,**** p<0.0001.

### MIDAP and HSF2BP form a multimeric complex with PALB2 and BRCA2

To further delineate the interactome of MIDAP, we immuno-precipitated MIDAP from mouse testis extracts coupled to mass spectrometry (IP-MS). Among the list of putative interactors, we identified HSF2BP as the main hit but also BRCA2, PALB2, RAD51 and RPA, strongly suggesting that they form a large multimeric complex (Table S4 and S5 for extended data). To confirm this result, we transfected the corresponding expression plasmids in HEK293T cells for co-immunoprecipitation analysis. MIDAP co-immunoprecipitated with BRCA2 and HSF2BP when they were all co-transfected but not when co-transfected only with BRCA2 (Figure 10a and S19a). The reciprocal co-IP of BRCA2 with HSF2BP and MIDAP was also positive. However, co-immunoprecipitation of MIDAP with BRCA2 was stronger in the presence of HSF2BP-S167L than in the presence of HSF2BP-WT (Figure 10a). In addition, we observed modest but positive co-immunoprecipitations of HSF2BP with RPA, PALB2 and RAD51; and of MIDAP with RAD51 and RPA but not with PALB2 (Figure S19b).

**Figure 10.**
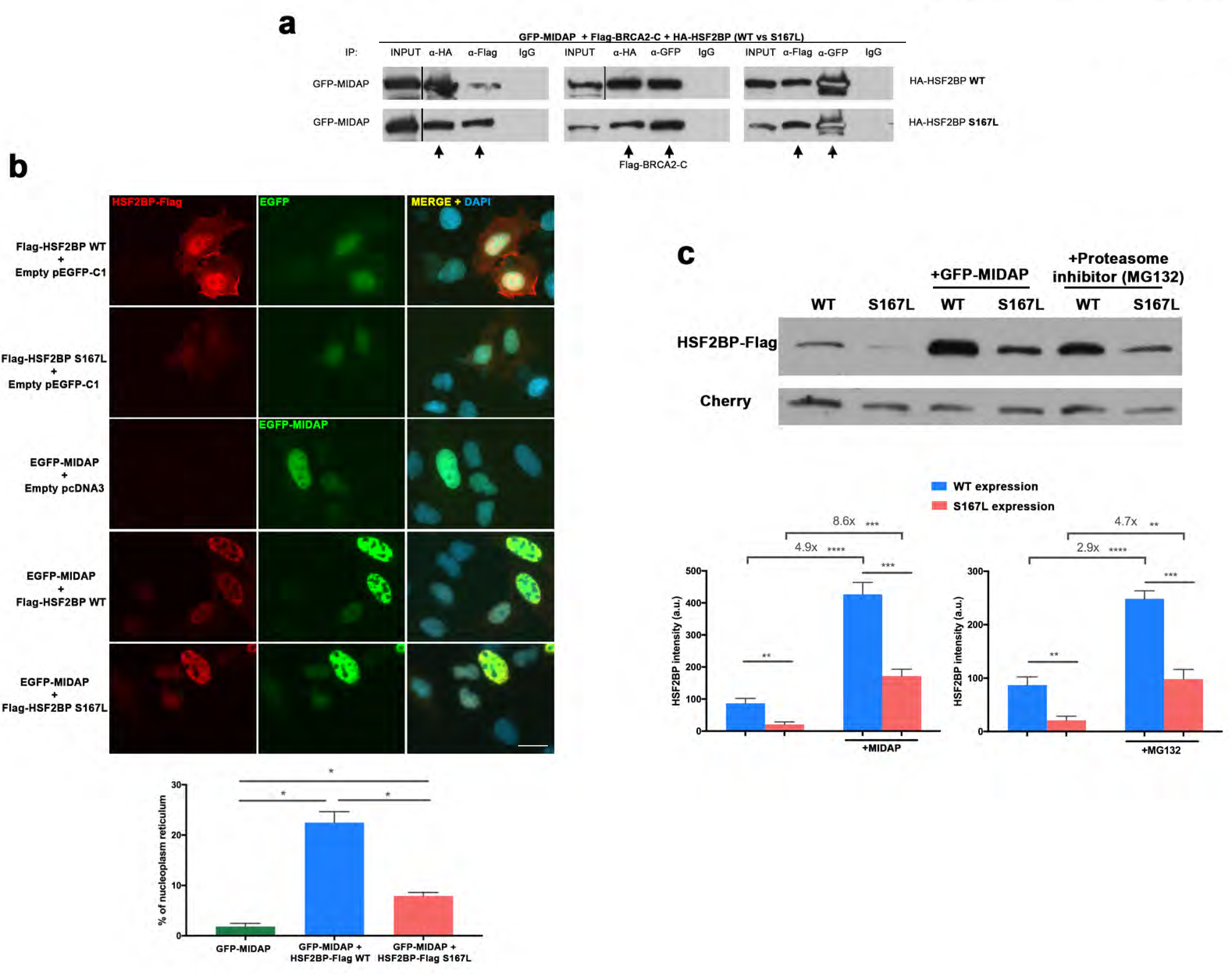
MIDAP forms a complex with BRCA2 and HSF2BP and stabilizes HSF2BP. (a) HEK293T cells were co-transfected with GFP-MIDAP, different Flag-BRCA2-C and HA-HSF2BP (WT and S167L variants). Protein complexes were immunoprecipitated overnight with either an anti-Flag, anti-EGFP, anti-HA or IgGs (negative control), and were analysed by immunoblotting with the indicated antibody. MIDAP does not co-immunoprecipitates with BRCA2-N, BRCA2-M or BRCA2-C (See Figure S19a) but when co-transfected with HSF2BP-HA (triple co-transfection) we detected co-immunoprecipitation between BRCA2-C and MIDAP. (b) Double immunofluorescence of transfected U2OS cells with plasmids encoding Flag-HSF2BP (WT or S167L) and EGFP-MIDAP alone or together and immuno-detected with antibodies against Flag (red) and EGFP (green). Transfected HSF2BP alone (both WT and S167L version) is delocalized and labels the whole cell (S167L less intense) whereas MIDAP shows nuclear localization. When co-transfected MIDAP and HSF2BP their localization patterns change and form nuclear invaginations that resemble nucleoplasmic reticulum. This phenotype is less severe in presence of HSF2BP-S167L than with the WT (graph under the panel: quantification of the number of cells with nucleoplasmic reticulum). Bar in panel, 20μm. (c) HEK293T cells were transfected with Flag-HSF2BP (WT and S167L) alone or with GFP-MIDAP. Additionally, cells transfected with Flag-HSF2BP were treated with the proteasome inhibitor (MG132, 10μM) during 4 hours and analysed by immunoblotting with a mouse anti-Flag antibody. Cherry was used as transfection efficiency control. HSF2BP-WT was expressed at higher levels than HSF2BP-S167L suggesting a reduced stability of the S167L variant. The level of HSF2BP expression (both the WT and the S167L variant) was increased when co-transfected with MIDAP. This increase of expression was greater in the HSF2BP-S167L variant in comparison with the WT (4,9 times in WT vs 8,6 times in S167L, see plots under the blot). After incubation with the proteasome inhibitor MG132, the expression levels of transfected HSF2BP were also increased mimicking the effect of MIDAP. This effect is also greater in the S167L variant than in the WT (2,9 times in WT and 4,7 in the S167L, see plot on the right). Welch’s t-test analysis: * p<0.05, ** p<0.01, *** p<0.001, **** p<0.0001.

To test more precisely their direct interactions, MIDAP, HSF2BP, PALB2, RPA and RAD51 were produced in a TNT and co-immunoprecipitated. The results showed an absence of direct interaction between any of them with the exception of MIDAP and HSF2BP (as expected from the Y2H analysis) (Figure S19c). Overall, these results suggest that despite the absence of direct interactions, these proteins are complexed *in vivo* likely through BRCA2 as observed in immunoprecipitations of testis extracts and provides clues on how the HSF2BP variant could be mechanistically operating by modulating MIDAP interaction with partners of major BRCA2-containing recombination complexes.

Given the close interaction of MIDAP and HSF2BP, we analyzed their interdependence in the heterologous system U2OS. We overexpressed MIDAP, HSF2BP (WT and S167L) and analyzed their cellular localization. Transfected HSF2BP localized to the cytoplasm and nucleus in a diffuse manner (Figure 10b). However, when HSF2BP was co-overexpressed with MIDAP, its pattern changed drastically to an intense nucleoplasm staining that also revealed the appearance of nuclear invaginations that resemble nucleoplasmic reticulum (Figure 10b) (Malhas et al., 2011). Interestingly, such invaginations were reduced when MIDAP was co-transfected with HSF2BP-S167L instead of WT (Figure 10b). In addition, the intensity of the fluorescence signal of the transfected HSF2BP-S167L was lower than for HSF2BP-WT. The fluorescence intensity of HSF2BP (both S167L and WT) was increased when co-transfected with MIDAP (Figure 10b). To further explore this observation, we analyzed by Western blot the expression levels of HSF2BP-WT and HSF2BP-S167L alone or in combination with MIDAP (Figure 10c). The results clearly showed that HSF2BP**-**S167L was expressed at lower amounts than the WT, indicating a reduced protein stability of the S167L variant and are in accordance also with the reduced expression of HSF2BP-S167L in the knock-in mouse *in vivo* (Figure 2). Interestingly, the expression of HSF2BP in transfected HEK293T cells is increased when co-transfected with MIDAP (greater effect in the S167L variant) and was partially dependent on proteasome-dependent degradation (MG132 treatment, Figure 10c), indicating a direct role of MIDAP in stabilizing HSF2BP. Taken altogether and given the low expression level of MIDAP in the *Hsf2bp^S167L/S167L^*, these results suggest that a strong functional interdependence between MIDAP and HSF2BP promotes their lower stability / expression in meiocytes, which might lead to the RPA accumulation, reduced RAD51 and DMC1 loading and consequently reduced COs.

## Discussion

Using exome sequencing, we identified the S167L missense variant in *HSF2BP* in a consanguineous family with three cases of POI with secondary amenorrhea. All affected family members are homozygous for the variant, and the healthy relatives are heterozygous carriers. The causality of the HSF2BP-S167L variant is supported by the meiotic phenotype and the subfertility observed in *Hsf2bp^S167L/S167L^* female mice. Furthermore, the DNA repair defects in murine KI spermatocytes, displayed by the accumulation of H2AX, the accumulation of RPA and the loss of loading of the RAD51/DMC1 recombinases on DSBs, provide direct evidence that this missense variant alters recombination. This conclusion was further supported by the comparative analysis of the POI allele with the *Hsf2bp* null allele that revealed that the missense variant can be considered as an hypomorphic allele of HSF2BP. This is in agreement with the secondary POI observed in the patients, and the residual (medically-assisted) fertility in one of the affected sisters. Our identification of *HSF2BP* as a gene implicated in POI is in line with recent reports of POI-causing variants in genes that are required for DNA repair and recombination, such as *MCM8, MCM9, SYCE1, MSH4, PSMC3IP, FANCM* or *NBN* (AlAsiri et al., 2015; Carlosama et al., 2017; de Vries et al., 2014; Fouquet et al., 2017; He et al., 2018; Tenenbaum-Rakover et al., 2015; Tucker et al., 2018; Wood-Trageser et al., 2014; Zangen et al., 2011).

We have shown by IP-MS that HSF2BP together with its novel interactor MIDAP forms a supramolecular complex with PALB2, RAD51, RPA and BRCA2. These interactions possibly are possibly mediated by the multidomain hub protein BRCA2 (Siaud et al., 2011) as we only detected direct interactions of HSF2BP with BRCA2 and MIDAP. In this regard, BRCA2 directly interacts with the DSBs recruiter PALB2, with the recombinases RAD51 and DMC1 (though through different specific domains), with DNA, with the DNA-mimicking protein and proteasome subunit DSS1 (which binds RPA complexes), and with HSF2BP (which binds MIDAP) (Siaud et al., 2011). This supramolecular complex participates in the orderly recruitment of essential players to the DSBs such as the initial binding of RPA to the resected DNA, exchange of RPA by RAD51 in a DSS1-dependent manner, and loading of the complex MEIOB-SPATA22 to the RPA complexes (Martinez et al., 2016; Zhao et al., 2015). Interestingly, genes with recently-identified variants in POI patients are implicated in the repair of induced DSBs at the early stages of meiosis and encode BRCA2-interacting factors, such as *MEIOB, DMC1* and *HSF2BP*, or *BRCA2* itself (Caburet et al., 2019a; He et al., 2018). This highlights the crucial importance and the high sensitivity of this particular meiotic step, and the hub role of BRCA2 as a tightly-regulated platform for correct meiotic recombination.

We have shown that genetic depletion of HSF2BP and MIDAP leads to meiotic arrest at zygotene-like stage. The arrested spermatocytes are not able to load the recombinases RAD51/DMC1, impairing the repair of DSBs and the generation of Cos, as shown by the absence of MLH1 foci). As a consequence, zygonema-like spermatocytes accumulate the SSB RPA. RPA, as part of a trimeric replication protein complex, binds and stabilizes ssDNA intermediates that forms during DNA repair. In meiosis, RPA is forming a heterodimer with the essential meiotic players MEIOB (homologue of RPA) and SPATA22. However, the loading of the complex formed by MEIOB/SPATA22 to DSBs is RPA-independent (Shi et al., 2019). It is postulated that RPA functions in meiosis at two different stages; i) during the early recombination stages when the DSBs ends are resected by the MRN complex and ii) during the strand invasion into the homologue duplex that is carried out by RAD51 and DMC1, ssDNA is generated at the displacement loops (Shi et al., 2019). The accumulation of RPA observed in the *Hsf2bp^-/-^* and *Midap*^-/-^ zygonema-like and in the *Hfs2bp^S167L/S167L^* spermatocytes is likely to occur at the early stages of recombination, because SPATA22 loading to the DSBs is also increased in the HSF2BP and MIDAP KOs and also to a lesser extent in the *Hsf2bp^S167L/S167L^*. We propose that this accumulation is not directly mediated by either HSF2BP or MIDAP, given the absence of direct interaction of HSF2BP/MIDAP to RPA, RAD51 and PALB2. In addition, the observed lack of DNA-binding ability of HSF2BP/MIDAP also points towards a model in which the absence of the heterodimer HSF2BP/MIDAP through a direct interaction with BRCA2 impairs the replacement of RPA by RAD51/DMC1 in the foci that form on the DSBs. Similarly, the POI variant, which does not affect the heterodimerization of HSF2BP/MIDAP, promotes a lower expression of both proteins of the heterodimer. This reduced amount of complex is thus less proficient in replacing RPA by the recombinases RAD51/DMC1 leading to a lower frequency of COs. The observed sexual dimorphism of HSF2BP and MIDAP mutants has also been described for several other meiotic genes in the mouse (Cahoon and Libuda, 2019). These differences can have a structural basis, given that the organization of the axial elements is known to be different between sexes. This is supported by the difference in length of the axes and by the essential role that the meiotic cohesin subunit RAD21L plays in males but not females (Herran et al., 2011). This distinct axial organization might generate different recombination landscapes in spermatocytes and oocytes. Accordingly, and similar to MIDAP/HSF2BP, the early meiotic recombination protein TEX15 is critical for loading recombinases in males but not in females, strongly suggesting a parallelism in the early recombination pathway where HSF2BP/MIDAP work (Yang et al., 2008). In general, meiotic recombination mutants seem to proceed further in female than in male meiosis perhaps because of the presence of less stringent checkpoints (Hunt and Hassold, 2002). In addition, human female meiosis seems to be especially error-prone (Bennabi et al., 2016) in comparison with other organisms (including the mouse), explaining the increased incidence of aneuploidy in the oocytes of aging women. It has recently been shown that this high level of metaphase I mis-segregation in women is due to a “female-specific maturation inefficiency” (Wang et al., 2017). This idea is based on the paradox that despite the total number of COs is higher in females than in males the frequency of bivalents without CO is also higher in women, and could also explain the relative milder phenotype of the mouse *Hsf2bp*^S167L/S167L^ oocytes compared with that of POI patients.

We have shown that the phenotype of female mice with S167L variants does not show a severe meiotic phenotype but only a slight reduction of fertility and a trend towards a reduction in the number of COs that can lead to univalents at metaphase I. Conversely, male mice bearing the mutation S167L did not show a statistically significant reduction in fertility but did show a reduction in the number of spermatozoa in the epididymis, the presence of apoptotic spermatocytes and a clear meiotic phenotype that resembles the KO phenotype but in a milder manner (reduction of COs that causes the presence of univalents at metaphase I). Interestingly, it is known that a reduction of the spermatozoa count (up to 60%) does not affect male mouse fertility which can be used to understand the molecular mechanism underlying a mutation (Schurmann et al., 2002). This can explain the normal fertility of male mice bearing the HSF2BP-S167L genetic variant despite a strong meiotic alteration.

Very recently, a high resolution genome-wide recombination map revealed novel loci involved in the control of meiotic recombination and highlighted genes involved in the formation of the synaptonemal complex (SYCE2, RAD21L, SYCP3, SIX6OS1) and the meiotic machinery itself as determinants of COs (Halldorsson et al., 2019). Within the second category, variants of the SUMO ligase RNF212 and the ubiquitin ligase HEI10 have been largely documented as genetic determinants of the recombination rate in humans, and importantly also of HSF2BP. Consequently, gene dosage of RNF212 and HEI10 affect CO frequency through their activity in CO designation and maturation (Lake and Hawley, 2013; Reynolds et al., 2013). We found that both MIDAP and HSF2BP localization are unaffected in the loss-of-function mutants of *Rnf212* and *Hei10*. This significant observation together with the proper co-localization of MIDAP/HSF2BP with RPA allows us to map these proteins upstream in the recombination pathway. Interestingly, a genetic variant of HSF2BP (Gly224Ter) affects recombination rate in males (Halldorsson et al., 2019). Two siblings homozygous for this HSF2BP variant in the analysed Icelandic population were healthy but none of them had descendants, suggesting they were infertile (Halldorsson et al., 2019).

It is interesting to note that some of the genes affecting the recombination rate have also been described as ‘fertility genes’, such as *SYCP3, HFM1* and *HSF2BP* ((Geisinger and Benavente, 2016; Wang et al., 2014) and this work). Altogether, we propose that different variants of the same meiotic gene (alleles responsible for mild or strong phenotypes) can give rise to either an altered genome-wide recombination rate with no detrimental effect, or cause infertility when the decreased recombination rate falls below the lower limit of one COs per bivalent. In the present POI family, the S167L variant in HSF2BP seems to be under that limit. To our knowledge, HSF2BP is one of the very few human genes with variants known to affect both the genome-wide recombination rate in the human population and meiotic chromosome missegregation (fertility) through a reduction of the recombination rate. Along similar lines, it is conceivable that variants with additive effects (Schimenti and Handel, 2018) can lead to a genome-wide reduction of the recombination rate and thus to aneuploidy and infertility. Specifically, variants in genes involved in meiotic recombination and SC constituents could be responsible for a large fraction of genetic idiopathic infertilities. These variants should be under purifying selection and would be removed or substantially reduced from the population. However, this is not the case for genes with sexual phenotypic dimorphism (Gershoni and Pietrokovski, 2014) as is apparent for a wide number of meiotic genes, including HSF2BP and MIDAP (Cahoon and Libuda, 2019), where individuals of one of the sexes are fertile carriers.

In summary, we describe for the first time a human family where POI co-segregates with a genetic variant in *HSF2BP* (S167L) in a Mendelian fashion and reveal that this variant promotes a lower expression of the heterodimer formed by HSF2BP and its novel stabilizer and interactor MIDAP. This hypomorphic expression of HSF2BP/MIDAP phenocopy in a mild manner the meiotic arrest observed in mice lacking either of HSF2B or MIDAP.

## MATERIAL AND METHODS

### Whole Exome Sequencing

Written informed consent was received from participants prior to inclusion in the study and the institutions involved. Genomic DNA was extracted from blood samples by standards protocols.

For individuals III-3 and III-10, library preparation, exome capture, sequencing and initial data processing were performed by Beckman Coulter Genomics (Danvers, USA). Exon capture was performed using the hsV5UTR kit target enrichment kit. Libraries were sequenced on an Illumina HiSEQ instrument as paired-end 100bp reads. For individual III-2, library preparation, exome capture, sequencing and data processing were performed by IntegraGen SA (Evry, France) according to their in-house procedures. Target capture, enrichment and elution were performed according to manufacturer’s instructions and protocols (SureSelect Human All Exon Kits Version CRE, Agilent). The library was sequenced on an Illumina HiSEQ 2500 as paired-end 75bp reads. Image analysis and base calling was performed using Illumina Real Time Analysis (RTA 1.18.64) with default parameters.

### Bioinformatic analysis

For the 3 individuals, sequence reads were mapped onto the human genome build (hg38 / GRCh38) using the Burrows-Wheeler Aligner (BWA) tool. Duplicated reads were removed using sambamba tools. WES metrics are provided in Table S1.

Variant calling, allowing the identification of SNV (Single Nucleotide Variations) and small insertions/deletions (up to 20bp) was performed via the Broad Institute GATK Haplotype Caller GVCF tool (3.7). Ensembl’s VEP (Variant Effect Predictor, release 87) program was used for initial variant annotation. This tool considers data available in dbSNP (dbSNP147), the 1000 Genomes Project (1000G_phase3), the Exome Variant Server (ESP6500SI-V2-SSA137), the Exome Aggregation Consortium (ExAC r3.0), and IntegraGen in-house databases. Additional annotation data was retrieved using dbNSFP (version 3.5, https://sites.google.com/site/jpopgen/dbNSFP) and Varsome (https://varsome.com/). Minor allele frequencies were manually verified on GnomAD (http://gnomad.broadinstitute.org), ISB Kaviar (http://db.systemsbiology.net/kaviar/), and Great Middle Eastern variant database GME Variome (http://igm.ucsd.edu/gme/).

Variant filtering was performed on the following criteria:

– minimum depth at variant position of 10,
– correct segregation in the family, on the basis of homozygosity by descent: variants should be homozygous in both affected sisters III-2 and III-3, and heterozygous or homozygous for Reference allele in the fertile sister III-10,
– absence in unrelated in-house fertile controls,
– Minor Allele Frequency (MAF) below 1% in global and in each population in the GnomAD database,
– presence in the coding sequence (i.e not in UTRs, introns, intergenic, …)
– high predicted functional impact on the protein. Impact was evaluated based on the predictors included in dbNSFP3.5 [14–16] (considered as pathogenic when the majority of the predictors agreed).

The number of variants fulfilling those criteria is provided in Table S2. Visual inspection of the variant was performed using the IGV viewer (Figure S1).

### Sanger Sequencing Analysis

To confirm the presence and segregation of the variant, direct genomic Sanger DNA sequencing of *HSF2BP* was performed in the patients, the parents and non-affected siblings using specific primers: HSF2BP-EX6F: 5’-CTAGAATCTTCTGTATCCTGCA-3’ and HHSF2BP-EX6R2: 5’-GGTCTGGAAGCAAACAGGCAA-3’. The resulting chromatograms are shown in Figure S2.

### Predictions of pathogenicity and sequence conservation

The S167L variant was predicted to be pathogenic or deleterious and highly conserved by 11 out of the 18 pathogenicity predictors available in dbNSFP 3.5 (Table S3). Upon verification, it appears that the conflicting interpretation of this variant might stem from the single occurrence of a Leu at this position in zebrafish. As the change in zebrafish is the variant that we have in the human family, we checked all the available sequences (Ensembl Release 99, January 2020, removing the one-to-many relationships). Ser167 is very highly conserved in mammals, birds and reptiles and fish and is present in 208 of 212 orthologous sequences (Figures S3 and S4).

### Generation of CRISPR/Cas9-Edited Mice

For developing all the mutant mice models (*Hsf2bp*^-/-^, *Hsf2bp*^S167L/S167L^, *Midap*^Δ/Δ^, *Midap*^-/-^, *Spo11*^-/-^, *Rnf212*^-/-^ and *Hei10*^-/-^) the different crRNAs were predicted at https://eu.idtdna.com/site/order/designtool/index/ <u>CRISPR_SEQUENCE.</u> The crRNAs, the tracrRNA and the ssODNs were produced by chemical synthesis at IDT (crRNAs and ssODNs sequences are listed in Tables S6 and S7). For the *Hsf2bp*^S167L^ we introduced a mutation in the mouse counterpart residue (p.Ser171Leu) of the POI mutation found in the clinical case (p.Ser167Leu). However, on this manuscript we will refer to the mutant allele by the acronym of the human mutation (S167L) to simplify. The ssODN contains the mutation on the corresponding position of the mouse sequence (c.512C>T, p.Ser171Leu, see character in red in Table S7) and the PAM mutations avoiding amino acid changes (see characters in bold in the Table S7). For the *Spo11*^-/-^ mice generation, the ssODN contains the mutations in the active site (TACTAC>TTCTTC p.YY137-138FF, see Table S7) and the PAM mutations (bold characters in Table S7). In all cases the crRNA and tracrRNA were annealed to obtain the mature sgRNA. A mixture containing the sgRNAs, recombinant Cas9 protein (IDT) and the ssODN (30 ng/μl Cas9, 20 ng/μl of each annealed sgRNA and 10 ng/μl ssODN) were microinjected into B6/CBA F2 zygotes (hybrids between strains C57BL/6J and CBA/J) (Singh et al., 2015) at the Transgenic Facility of the University of Salamanca. Edited founders were identified by PCR amplification (Taq polymerase, NZYtech) with primers flanking the edited region (see Table S8 for primer sequences). The PCRs products were direct sequenced or subcloned into pBlueScript (Stratagene) followed by Sanger sequencing, selecting the founders carrying the desired alelles. The selected founders were crossed with wild-type C57BL/6J to eliminate possible unwanted off-targets. Heterozygous mice were re-sequenced and crossed to give rise to edited homozygous. Genotyping was performed by analysis of the PCR products produced from genomic DNA extracted from tail biopsies. The primers and the expected amplicon sizes are listed in the Table S8. Mouse mutants for *Rec8, Six6os1* and *Psma8* have been previously described (Bannister et al, 2004; Gomez et al, 2019; Gomez et al, 2016).

### Ethics statement

Mice were housed in a temperature-controlled facility (specific pathogen free, spf) using individually ventilated cages, standard diet and a 12 h light/dark cycle, according to EU laws at the “Servicio de Experimentación Animal, SEA”. Mouse protocols have been approved by the Ethics Committee for Animal Experimentation of the University of Salamanca (USAL). We made every effort to minimize suffering and to improve animal welfare. Blinded experiments were applied when possible but for the *Hsf2bp*^-/-^ and *Midap*^-/-^ mutants it was not possible since the phenotype was obvious between wild-type and these mutant mice for all of the experimental procedures used. No randomization methods were applied since the animals were not divided in groups/treatments. The minimum size used for each analysis was two animals/genotype. The mice analyzed were between 2 and 4 months of age, except in those experiments where is indicated.

### Histology

For histological analysis, after the necropsy of the mice their testes or ovaries were removed and fixed in Bouińs fixative or formol 10%, respectively. They were processed into serial paraffin sections and stained with haematoxylin-eosin (ovaries) or Periodic acid–Schiff (PAS) and hematoxylin (testes). The samples were analysed using a microscope OLYMPUS BX51 and images were taken with a digital camera OLYMPUS DP70.

### Fertility assessment

*Hsf2bp*^+/+^ and *Hsf2bp*^S167L/S167L^ males and females (8 weeks old) were mated with wild type females and males, respectively, over the course of 4-12 months. 6 mice per genotype (7 mice for *Hsf2bp* ^S167L/S167L^ females) were crossed. The presence of copulatory plug was examined daily and the number of pups per litter was recorded.

### Immunocytology and antibodies

Testes were detunicated and processed for spreading using a conventional “dry-down” technique. Oocytes from fetal ovaries (E16.5, E17.5 and E19.5 embryos) were digested with collagenase, incubated in hypotonic buffer, disaggregated and fixed in paraformaldehyde. Rabbit polyclonal antibodies (R1 and R2 generated from two different host rabbits) against HSF2BP and MIDAP were developed by ProteintechTM against a fusion protein of poly-His with full length HSF2BP or MIDAP (pUC57 vector) of mouse origin. Rabbit polyclonal antibody against DMC1 was developed by ProteintechTM against a DMC1 peptide (EESGFQDDEESLFQDIDLLQKHGINMADIKKLKSVGICTIKG).

The primary antibodies used for immunofluorescence were rabbit αHSF2BP R2 (1:30, ProteintechTM), rabbit αMIDAP R2 (1:100, ProteintechTM), mouse αSYCP3 IgG sc-74569 (1:100, Santa Cruz), rabbit α-SYCP3 serum K921 (provided by Dr. José Luis Barbero, Centro de Investigaciones Biológicas, Spain), rabbit αSYCP1 IgG ab15090 (1:200, Abcam), rabbit anti-γH2AX (ser139) IgG #07-164 (1:500, Millipore), mouse αMLH1 51-1327GR (1:20, BD Biosciences), mouse αCDK2 (1:20; Santa Cruz Sc-6248) rabbit αRAD51 PC130 (1:50, Calbiochem), rabbit αRPA serum “Moll” (1:30, provided by Dr. Edyta Marcon, Medical Research University of Toronto, Canada), rabbit αDMC1 (1:500, ProteintechTM), rabbit αSPATA22 16989-1-AP (1:60, Proteintech), mouse αFlag IgG (1:100; F1804, Sigma-Aldrich).

### Generation of plasmids

Full-length cDNAs encoding HSF2BP, MIDAP (full length and delta constructs), RPA1, BRCA2 (N, M and C constructs), TEX11, PALB2, RAD51, and PSMA8 were RT-PCR amplified from murine testis RNA. The cDNAs were cloned into the EcoRV pcDNA3-2XFlag, SmaI pcDNA3-2XHA or SmaI pEGFP-C1 expression vectors under the CMV promoter. In frame cloning was verified by Sanger sequencing.

### Y2H assay and screening

Y2H assay was performed using the Matchmaker Gold Yeast Two-Hybrid System (Clontech) according to the manufacturers’ instructions. Mouse *Hsf2bp* cDNA was subcloned into the vector pGBKT7 and was used as bait to screen a mouse testis Mate & Plate cDNA library (Clontech Laboratories Inc.). Positive clones were initially identified on double dropout SD (synthetic dropout)/–Leu/–Trp/X-α-Gal/Aureobasidin A plates before further selection on higher stringency quadruple dropout SD/–Ade/–His/–Leu/–Trp/X-α-Gal/Aureobasidin A plates. Pray plasmids were extracted from the candidate yeast clones and transformed into *Escherichia coli*. The plasmids from two independent bacteria colonies were independently grown, extracted and Sanger sequenced.

### DNA pull down assay

ssDNA/dsDNA pull down assays were performed using the protocol previously described by (Souquet et al., 2013). A HPLC-purified biotinylated oligonucleotide was used for the DNA pull down assays: ss60-mer F: 5′-GAT CTG CACGACGCACACCGGACGTATCTGCTATCGCTCATGTCAACCGCTCAAGCTGC/3’ BiotinTEG/ (IDT) and ss60-mer R (No biotinylated): 5′-GCAGCTTGAGCGGTTGACATGAGCGATAGCAGATACGTCCGGTGTGCGTCGTGCAGATC-3’. Double-stranded DNA annealing was carried out in 50 mM NaCl, 25 mM Tris-HCl, pH 7.5 buffer with complementary sequences at molecular equivalence by a denaturing step (5 minutes at 95°C) and a slow return to room temperature. DNA was immobilized onto Dynabeads M-280 Streptavidin (Dynal) following the manufacturer instructions (0.2 pmol per 1 µg of beads). Protein extracts were obtained from *in vitro* coupled transcription/translation systems (TNT® T7 Coupled Reticulocyte Lysate Systems, Promega). 15 µl of Flag-tagged proteins from TNT assays were pre-incubated on ice for 10 minutes in modified DBB (DBB: 50 mM Tris HCl, 100mM NaCl, 10% (w/v) glycerol,

Complete Protease inhibitor, 1mM 2-mercaptoethanol pH 7,4 modified with 25 mM Tris-HCl, 1 mM EDTA plus 5 mg/ml BSA). After this preincubation 500 µg Dynabeads with immobilized ss- or ds-DNA were added and incubated for 1 hour at 4°C under agitation. Then the beads were washed three times (5 min rotating at RT) in 700 µl of modified DBB without BSA, before being washed once in 700 µl of rinsing buffer (modified DBB with 150 mM NaCl). Finally, DNA binding proteins were eluted by resuspending the beads in 30 µl of Laemmli buffer boiling the samples for 5 min. The samples were analyzed by western blot.

### Cell lines and transfections

HEK293T and U2OS cell lines were obtained from the ATCC and transfected with Jetpei (PolyPlus) according to the manufacturer protocol. Cell lines were tested for mycoplasma contamination using the Mycoplasma PCR ELISA (Sigma).

### Immunoprecipitation and western blotting

HEK293T cells were transiently transfected and whole cell extracts were prepared in a 50mM Tris-HCl pH 7,4, 150mM NaCl, 1mM EDTA, 1% Triton X-100 buffer supplemented with protease inhibitors. Those extracts were cleared with protein G Sepharose beads (GE Healthcare) for 1 h. The corresponding antibodies were incubated with the extracts for 2 h and immunocomplexes were isolated by adsorption to protein G-Sepharose beads o/n. After washing, the proteins were eluted from the beads with 2xSDS gel-loading buffer 100mM Tris-Hcl (pH 7), 4% SDS, 0.2% bromophenol blue, 200mM β-mercaptoethanol and 20% glycerol, and loaded onto reducing polyacrylamide SDS gels. The proteins were detected by western blotting with the indicated antibodies. Immunoprecipitations were performed using mouse αFlag IgG (5µg; F1804, Sigma-Aldrich), mouse αGFP IgG (4 µg; CSB-MA000051M0m, Cusabio), ChromPure mouse IgG (5µg/1mg prot; 015-000-003). Primary antibodies used for western blotting were rabbit αFlag IgG (1:2000; F7425 Sigma-Aldrich), goat αGFP IgG (sc-5385, Santa Cruz) (1:3000), rabbit αMyc Tag IgG (1:3000; #06-549, Millipore), rabbit αHSF2BP R2 (1:2000, ProteintechTM), rabbit αMIDAP R1 (1:3000, ProteintechTM), rat αRPA2 (1:1000, Cell Signaling (Cat 2208S)). Secondary horseradish peroxidase-conjugated α-mouse (715-035-150, Jackson ImmunoResearch), α-rabbit (711-035-152, Jackson ImmunoResearch), α-goat (705-035-147, Jackson ImmunoResearch) or α-rat (712-035-150, Jackson ImmunoResearch) antibodies were used at 1:5000 dilution. Antibodies were detected by using Immobilon Western Chemiluminescent HRP Substrate from Millipore.

### Testis immunoprecipitation coupled to MS/MS analysis

200 µg of antibodies R1 and R2 (against HSF2BP and MIDAP) were crosslinked to 100 ul of sepharose beads slurry (GE Healthcare). Testis extracts were prepared in 50mM Tris-HCl (pH8), 500mM NaCl, 1mM EDTA 1% Triton X100. 20 mg of protein extracts were incubated o/n with the sepharose beads. Protein-bound beads were packed into columns and washed in extracting buffer for three times. Proteins were eluted in 100 mM glycine pH3 and analysed by Lc-MS/MS shotgun in LTQ Velos Orbitrap at the Proteomics facility of Centro de Investigación del Cáncer (CSIC/University of Salamanca).

### MS/MS data analysis

Raw MS data were analyzed using MaxQuant v 1.6.2.6 (Cox and Mann, 2008) against SwissProt Mouse database (UP000000589, Oct, 2019) and MaxQuant contaminants. All FDRs were of 1%. Variable modifications taken into account were oxidation of M and acetylation of the N-term, while fixed modifications included considered only carbamidomethylation of C. The maximum number of modifications allowed per peptide was of 5. Proteins were quantified using iBAQ (Schwanhausser et al., 2011). Potential contaminants, reverse decoy sequences and proteins identified by site were removed. Proteins with less than two unique peptides in the Ab1 and Ab2 groups were not considered for ulterior analysis. Proteins with less than two unique peptides in the control group and more than two in both groups Ab1 and Ab2 were selected as high-confidence candidates (group Ab1 and Ab2 only). An additional group of putative candidates was selected for those proteins with two or more unique peptides in one of the Ab1 or Ab2 groups and no unique peptides in the control sample (groups Ab1 only and Ab2 only, respectively).

### Statistics

In order to compare counts between genotypes, we used the Welch’s t-test (unequal variances t-test), which was appropriate as the count data were not highly skewed (i.e., were reasonably approximated by a normal distribution) and in most cases showed unequal variance. We applied a two-sided test in all the cases. Asterisks denote statistical significance: *p-value <0.05, **p-value <0.01, ***p-value<0.001 and ****p-value<0.0001.

## Online Supplemental material

Table S1 shows the WES and mapping metrics for the 3 genomic samples. Table S2 shows the numbers of variants from the WES analysis and passing the various filters. Table S3 shows the predictions of pathogenicity and conservations by 18 computational predictors.

Tables S4 and S5 show the candidate MIDAP interactors identified by IP-MS. Table S6 and S7 show respectively the crRNAs and the ssODN employed in the generation of the various mouse models. Table S8 shows the primers and expected product sizes for genotyping mouse models. Figure S1 shows the pedigree for the consanguineous family with 3 POI cases. Figure S2 shows the chromatograms obtained by Sanger sequencing of the HSF2BP-S167L variant in the family. Figure S3 and S4 show the conservation of the S167 position in mammals, birds and reptiles, and fish. Figure S5 shows the generation and genetic characterization of *Hsf2bp*^Ser167Leu^ and *Hsf2bp*-deficient mouse models. Figure S6 shows the absence of synaptic defects in *Hsf2bp*^S167L/S167L^ meiocytes. Figure S7 shows apoptotic metaphases I in *Hsf2bp*^S167L/S167L^ males. Figure S8 shows the defective DNA repair in *Hsf2bp*^-/-^ mice. Figure S9 shows Comparative interaction of HSF2BP-S167L and HSF2BP-WT with BRCA2. Figure S10 shows the absence of MIDAP and HSF2BP loading defects in the *Midap* Δ142-472 mutant. Figure S11 shows the localization of C19orf57/MIDAP at meiotic recombination nodules and the absence of DNA binding abilities in both HSF2BP and MIDAP. Figure S12 and S13 show the generation and genetic characterization of *Spo11^-/-^*

, *Rnf212^-/-^*and *Hei10^-/-^* mice. Figure S14 shows the loading of MIDAP and HSF2BP on in different synapsis/recombination-defective mutant mice. Figure S15 shows the generation and genetic characterization of *Midap*^-/-^ mice. Figure S16 shows the accumulation of H2AX and RPA in *Midap*^-/-^ males. Figure S17 shows the defect on MLH1 localization in *Midap*^-/-^ mice. Figure 18 shows the accumulation of SPATA22 in *Midap*^-/-^, *Hsf2bp^S167L/S167L^* and *Hsf2bp*^-/-^ mice. Figure S19 shows the co-immunoprecipitations between HSF2BP, MIDAP, RPA, RAD51 and PALB2.

### Author contributions

Conceptualization: RAV and AMP

Formal analysis: SC, ALT, RAV, RGV, DDR, NFM, FSS, YBC, ELC, PD, LGH, AMP

Resources: SAS, NFM, MSM.

Investigation: SC, ELC, ALT, RAV, NFM, AMP Writing – original draft & Visualization: SC, RAV, AMP. Writing – review & editing: SC, RAV, AMP.

Funding acquisition: RAV, AMP

## Acknowledgements

We thank Dr. Barbero and Dr. Edyta Marcon, for providing antibodies against SYCP3 and RPA, respectively, Dr. Emmanuelle Martini for helpful advice with the DNA binding assay and Dr. Fabien Fauchereau for the genetic analysis of the family. This study was supported by Université Paris Diderot and the Fondation pour la Recherche Médicale (Labelisation Equipes DEQ20150331757, S. C., A-L. T and R. A. V.). This work was supported by MINECO (BFU2017-89408-R) and by Junta de Castilla y León (CSI239P18). NFM, FSS and LGH are supported by European Social Fund/JCyLe grants (EDU/310/2015, EDU/556/2019 and EDU/1083/2013). YBC is funded by a grant from MINECO (BS-2015-073993). The proteomic analysis was performed in the Proteomics Facility of Centro de Investigación del Cáncer, Salamanca, Grant PRB3(IPT17/0019 -ISCIII-SGEFI / ERDF). CIC-IBMCC is supported by the Programa de Apoyo a Planes Estratégicos de Investigación de Estructuras de Investigación de Excelencia cofunded by the Castilla–León autonomous government and the European Regional Development Fund (CLC–2017–01). The funders had no role in study design, data collection and analysis, decision to publish, or preparation of the manuscript.

## Legends to Supplementary figures

**Figure S1.**
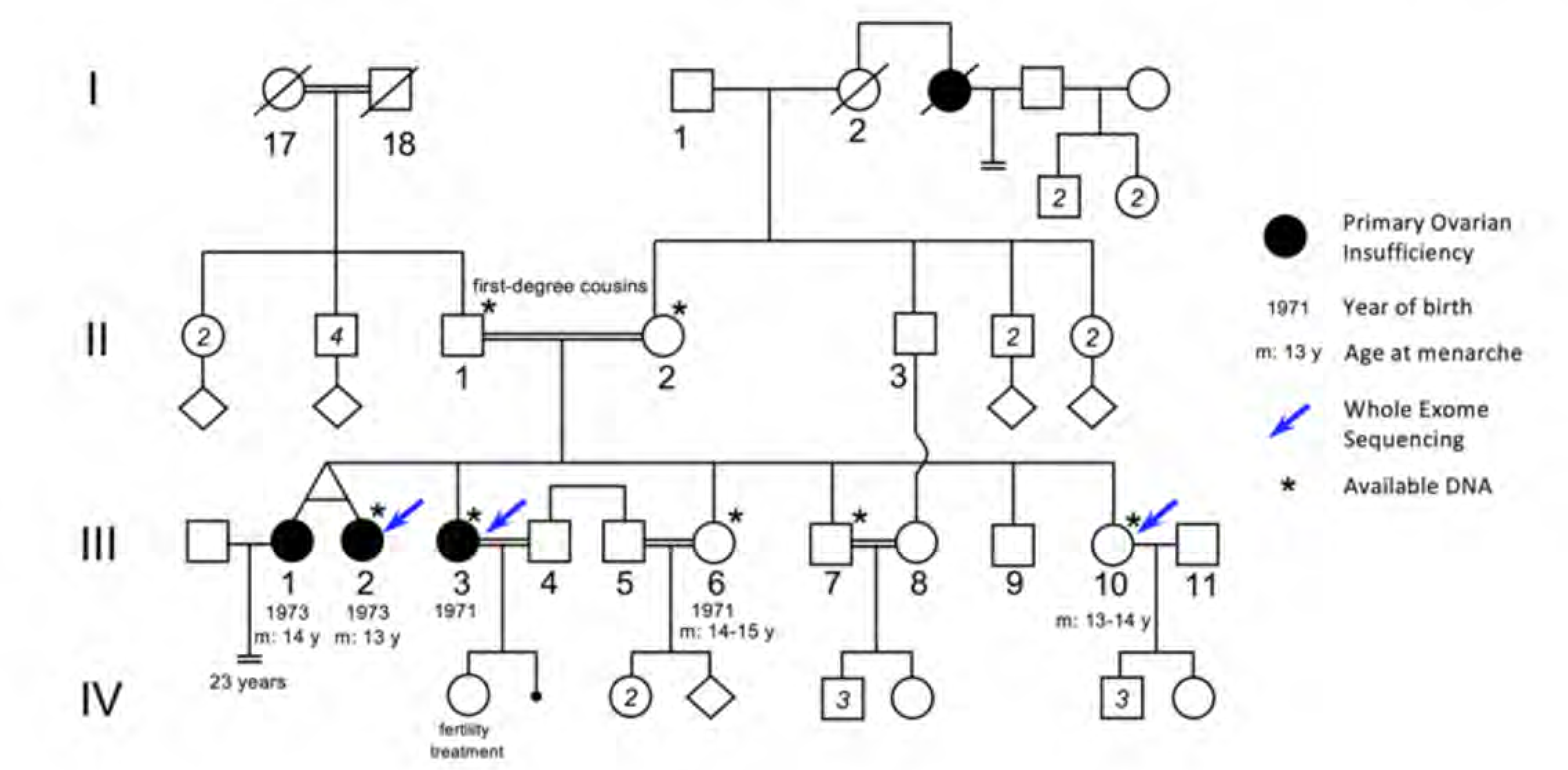
Pedigree of the consanguineous family with the variant HSF2BP-S167L. III-1 and III-2 are monozygotic twins, who appear phenotypically dizygotic. Clinical investigation confirmed POI, with normal 46, XX karyotype (500 bands and SKY spectral karyotyping). Year of birth and age of menarche are indicated when known. III-1 became amenorrheic at age 24 and III-2 at age 25, both after irregular menstruations since menarche. III-1 presents with a short stature (152 cm, within the 3-5 percentile), a normal neck, cubitus valgus and metacarpal shortening of 4-5. Ultrasound investigation showed normal uterus and ovaries. Her g-banding karyotyping was normal 46, XX (500 bands) and variants in FMR1 gene were ruled out. III-2 displays a normal secondary sexual development with no dysmorphic sign. Clinical investigation confirmed POI, with normal 46, XX karyotype (500 bands and SKY spectral karyotyping). The elder sister III-3 was also diagnosed with POI, with no further clinical information. She is 160.5 cm. She had one normal pregnancy with the help of “fertility treatment”, and a second unsuccessful attempt. The two fertile sisters, III-6 and III-10 had their menarche at 14-15 and 13-14 respectively, with regular menstruations ever since. They are respectively 150 cm and 151 cm, with no clinical sign, and each one had several children without difficulties. The fertile brother III-7 is 171 cm, and shows no health or fertility problem. He developed frontal baldness since the age of 30.

**Figure S2.**
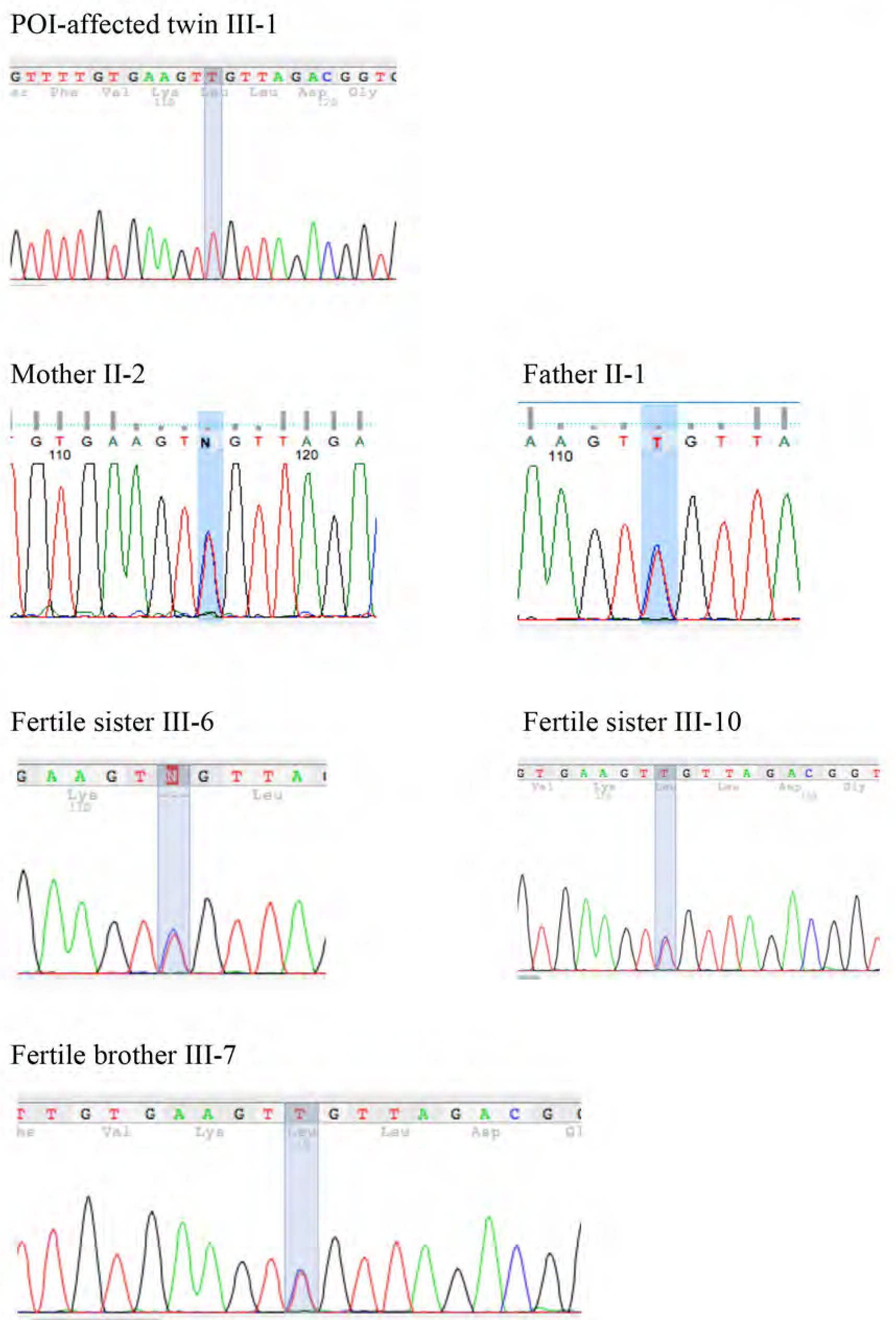
Segregation of the Ser167Leu Variant in HSF2BP in the consanguineous family.

**Figure S3.**
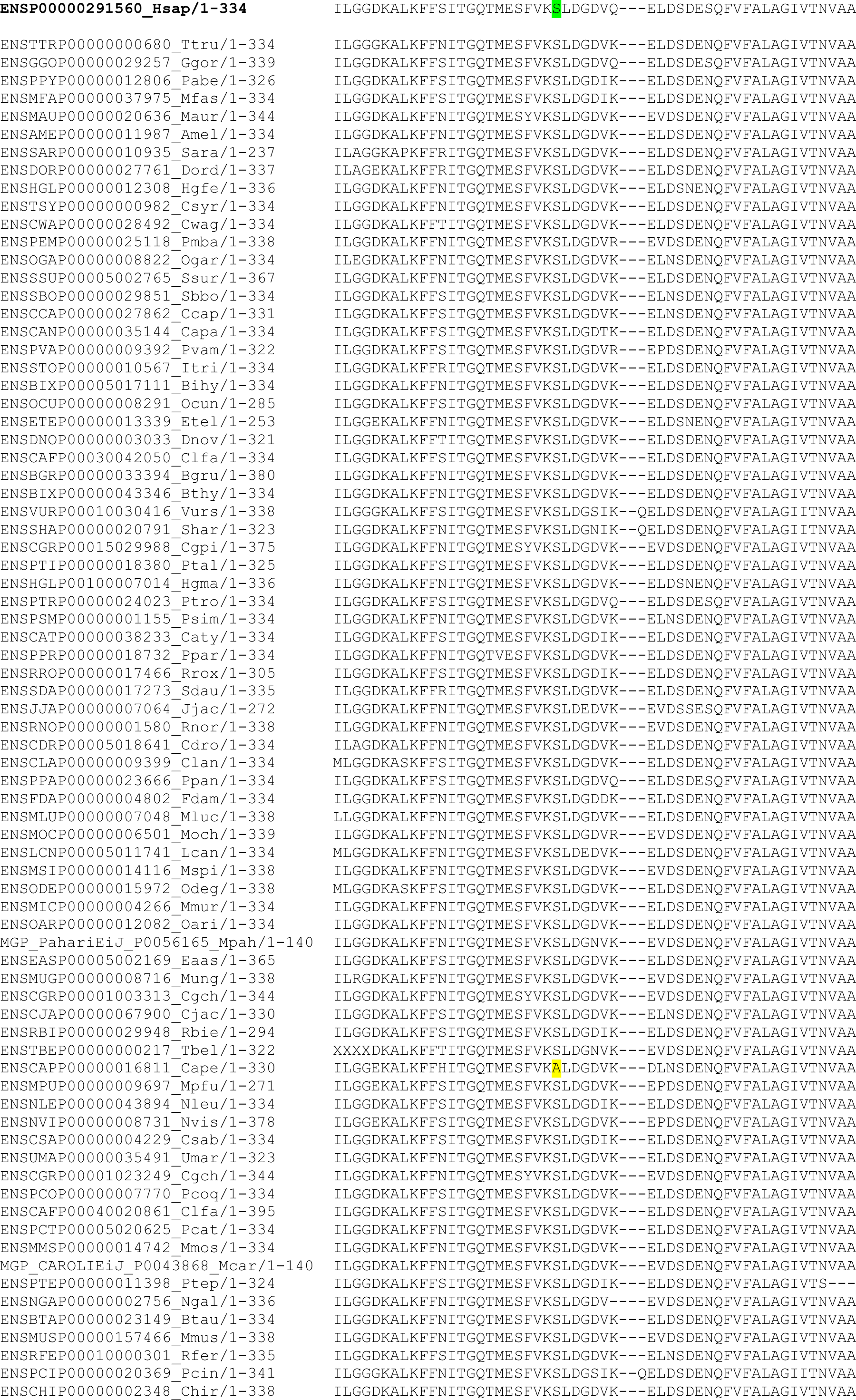
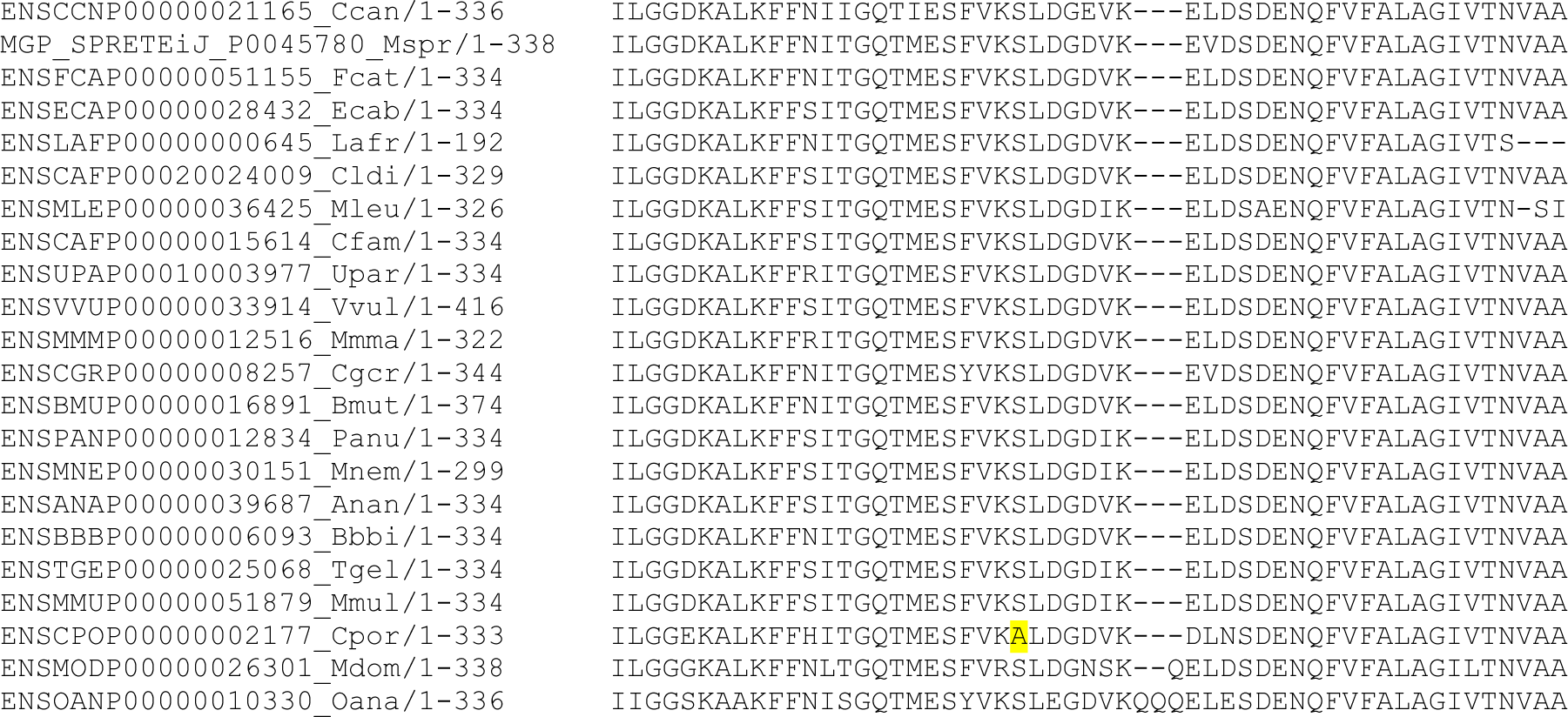
Strong conservation of the Ser167 residue in HSF2BP protein in 99 mammals. CLUSTAL W 2.0 multiple sequence alignment of one-to-one orthologues. The human Ser167 (green) is conserved in 97 of the 99 available sequences in mammals. The only two changes are highlighted in yellow in *Cavia aperea* (Cape, Brazilian guinea pig) and *Cavia porcellus* (Cpor, guinea pig).

**Figure S4.**
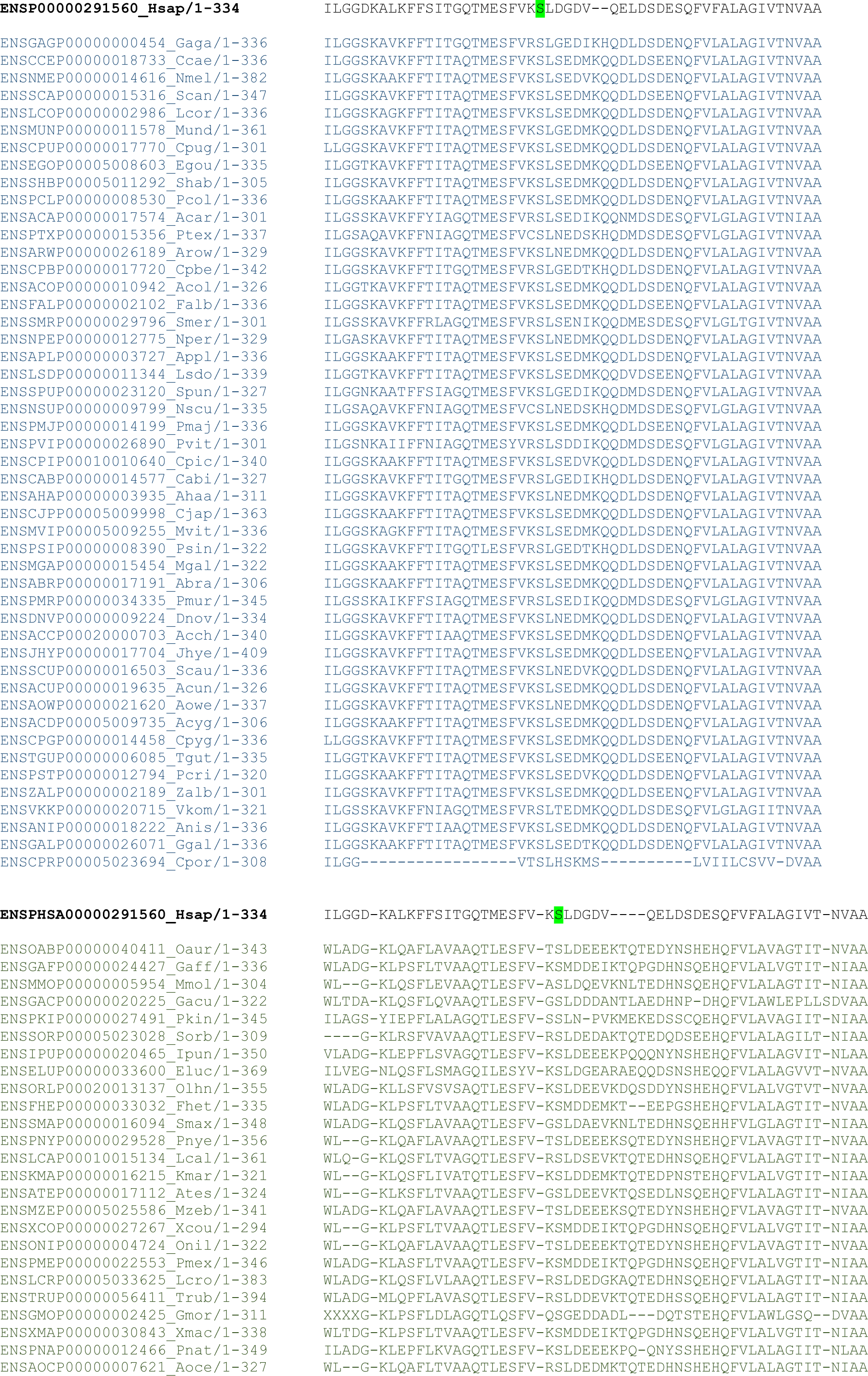
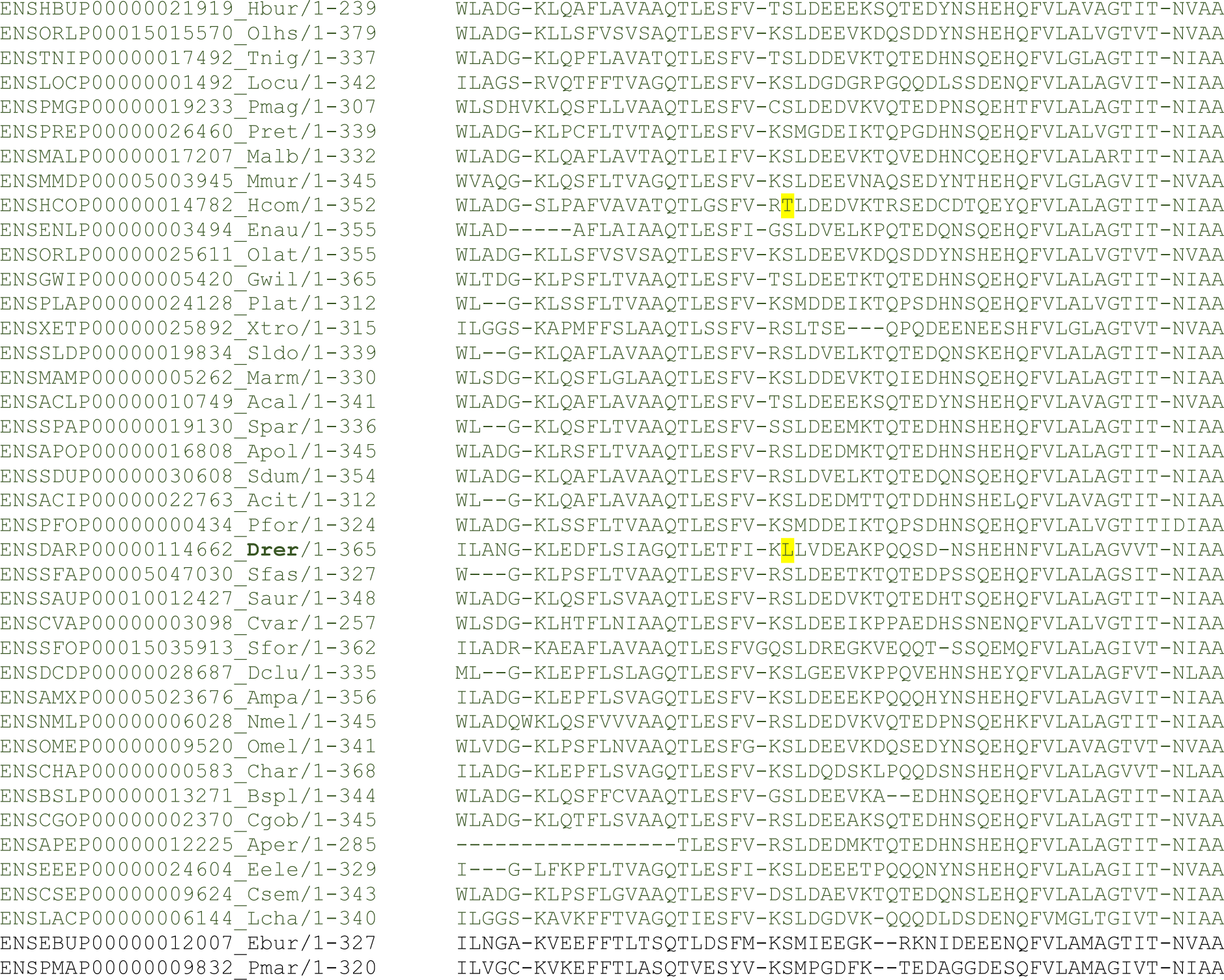
Strong conservation of the Ser167 residue in HSF2BP in 48 birds and reptiles and 64 fish species. CLUSTAL W 2.0 multiple sequence alignment of one-to-one orthologues. The human Ser167 position (green) is strictly conserved in the 48 available sequences from birds and reptiles (sequences in dark blue) and is conserved in 63 of the 65 available sequences in fish and jawless fish (sequences in dark green). The only two changes in fish species are highlighted in yellow in *Hippocampus comes* (Hcom, Tiger tail seahorse) and *Danio rerio* (Drer, zebrafish). All other fish have a Ser in this position, including the Coelacanth and the more distantly-related jawless vertebrates, the Lamprey and the Hagfish.

**Figure S5.**
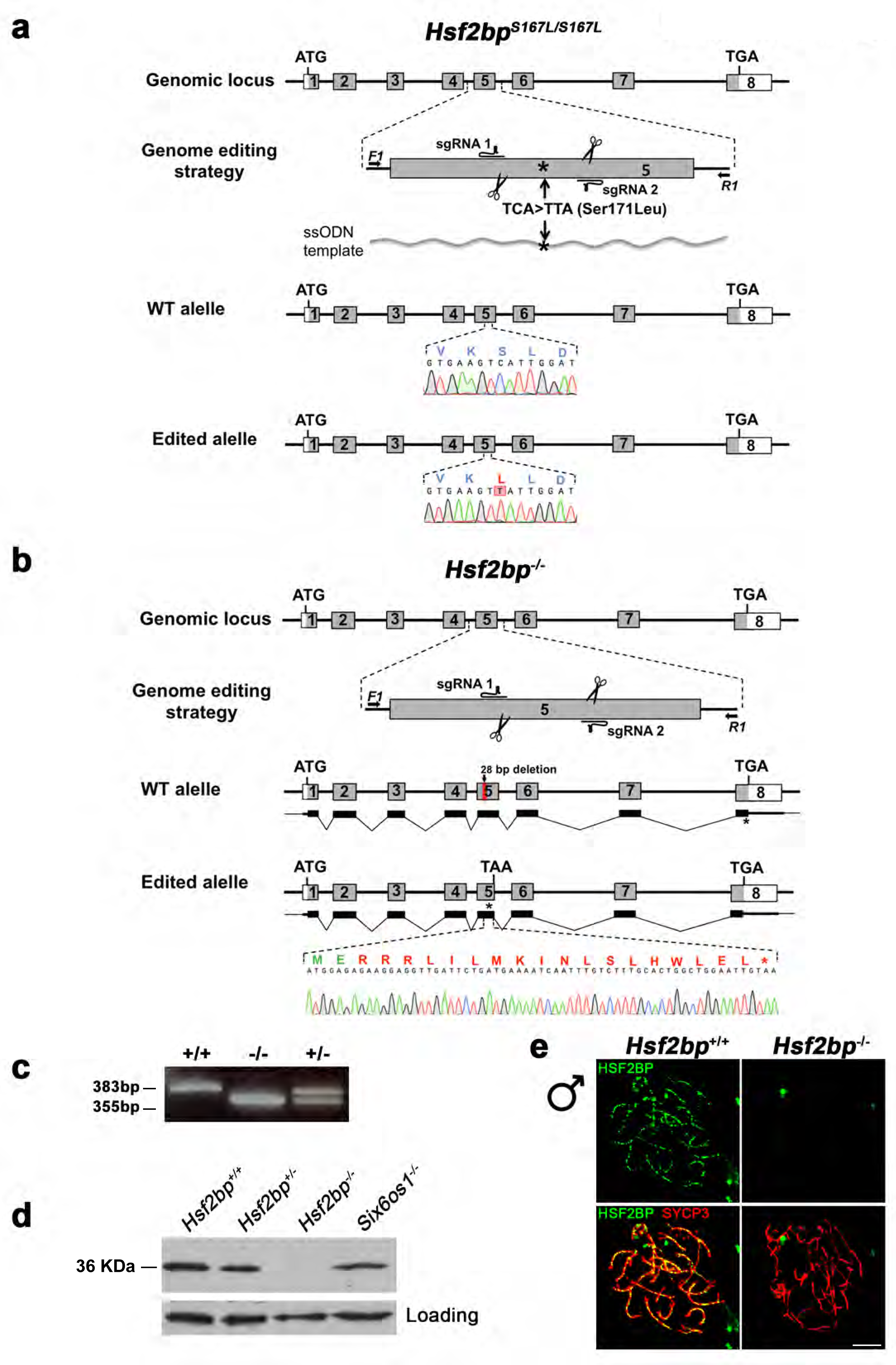
Generation and genetic characterization of *Hsf2bp* Ser167Leu and *Hsf2bp-*deficient mice. (a) Diagrammatic representation of the mouse *Hsf2bp* locus (WT) and the genome editing strategy (see methods) showing the sgRNAs located on exon 5 and the ssODN bearing the desired mutation (TCA>TTA, Ser171Leu) on the homologous residue of the variant identified in the family affected with POI (Ser167Leu). The murine allele is refered with the human variant (S167L). The corresponding coding exons (light grey) and non-coding exons (open boxes) are represented. Thin (non-coding) and thick (coding sequences) lines under exons represent the expected transcript derived from wild-type and *Hsf2bp* edited allele. ATG, initiation codon; TGA and *, stop codon. The nucleotide sequence of the WT and edited allele derived from PCR amplification of DNA from *Hsf2bp*^+/+^ and *Hsf2bp*^S167L/S167L^ mice is indicated. Arrows represent genotyping primers (F1 and R1). (b) Diagrammatic representation of the mouse *Hsf2bp* locus (WT) and the genome editing strategy showing the sgRNAs located on exon 5. ATG, initiation codon; TGA/TAA and *, stop codon. The nucleotide sequence of the edited allele derived from PCR amplification of DNA from the *Hsf2bp^-/-^* is indicated. Arrows represent genotyping primers (F1 and R1). **(c)** PCR analysis of genomic DNA from three littermate progeny of *Hsf2bp^+/-^* heterozygote crosses. The PCR amplification with primers F1 and R1 revealed 383 and 355 bp fragments for wild-type and disrupted alleles respectively. Wild-type (+/+), heterozygous (+/-), and homozygous knock-out (-/-) animals. (d) Western blot analysis of protein extracts from wild type and *Hsf2bp^-/-^* testis with a specific antibody against whole recombinant HSF2BP protein. An unspecific band was used as loading control. (e) Double immunofluorescence of spermatocytes at zygotene/zygotene-like stage obtained from *Hsf2bp^+/+^* and *Hsf2bp^-/-^* mice using SYCP3 (red) and a rabbit polyclonal antibody against HSF2BP (green, see methods). Bar in panel, 10 µm.

**Figure S6.**
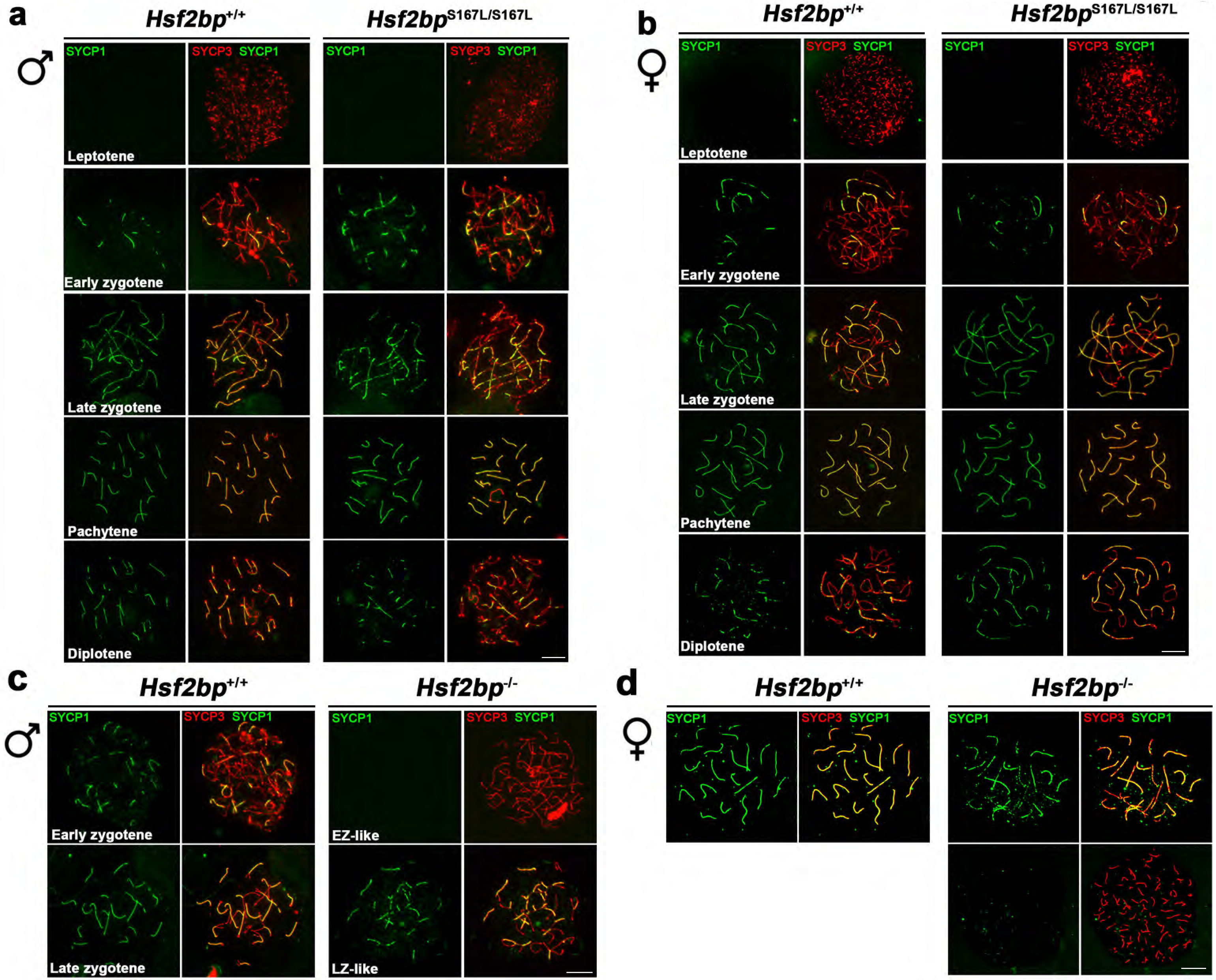
*Hsf2bp*^S167L/S167L^ mice do not show synapsis defects. (a-b) Double immunolabelling of spermatocyte (a) and oocyte (b) spread preparations with SYCP1 (green) and SYCP3 (red), showing that meiotic prophase proceeds with no defects in synapsis/desynapsis in *Hsf2bp*^S167L/S167L^ mutants. (c) Immunofluorescence of spermatocyte spreads of wild-type and *Hsf2bp^-/-^* with SYCP3 (red) and SYCP1 (green). *Hsf2bp^-/-^* males show a spermatogenic arrest at zygotene-like displaying synapsis between non-homologous chromosomes (partner switch phenotype). (d) Immunofluorescence of oocyte spreads (17,5 dpp) of wild-type and *Hsf2bp^-/-^* with SYCP3 (red) and SYCP1 (green). *Hsf2bp^-/-^* females show a delay in prophase I progression with an increase of cells with synapsis defects. Bar in panels, 10 µm.

**Figure S7.**
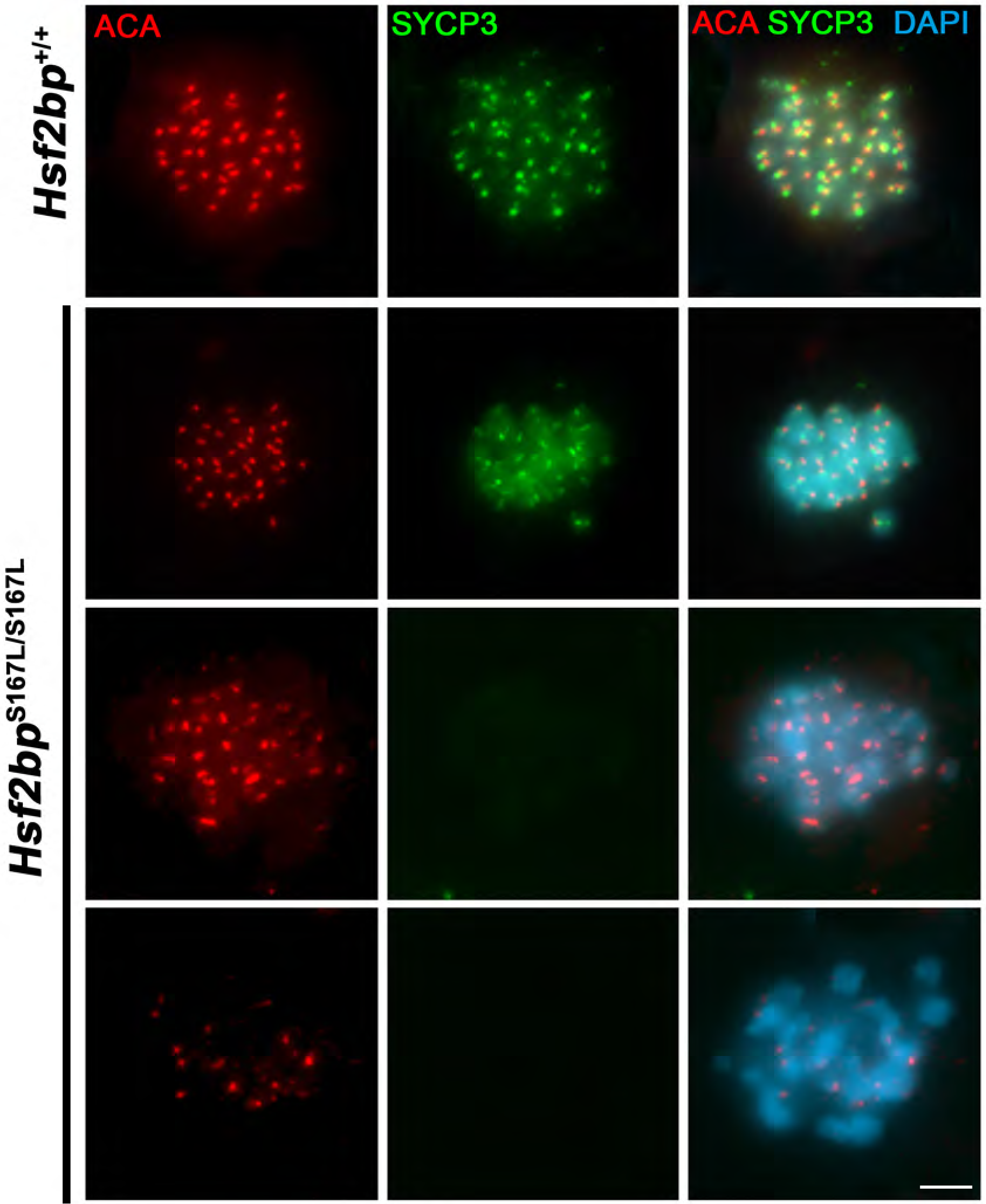
Metaphase I defects in *Hsf2bp*^S167L/S167L^ males. Co-labeling of SYCP3 (green) and ACA (red) in squashed preparations of tubules from WT and S167L mice. In the mutant mice, a large number of metaphases I were apoptotic with partial or total loss of immunoreactivity for the labelled proteins. See quantification of apoptotic cells in Figure 1h. Bar in panel, 10 µm.

**Figure S8.**
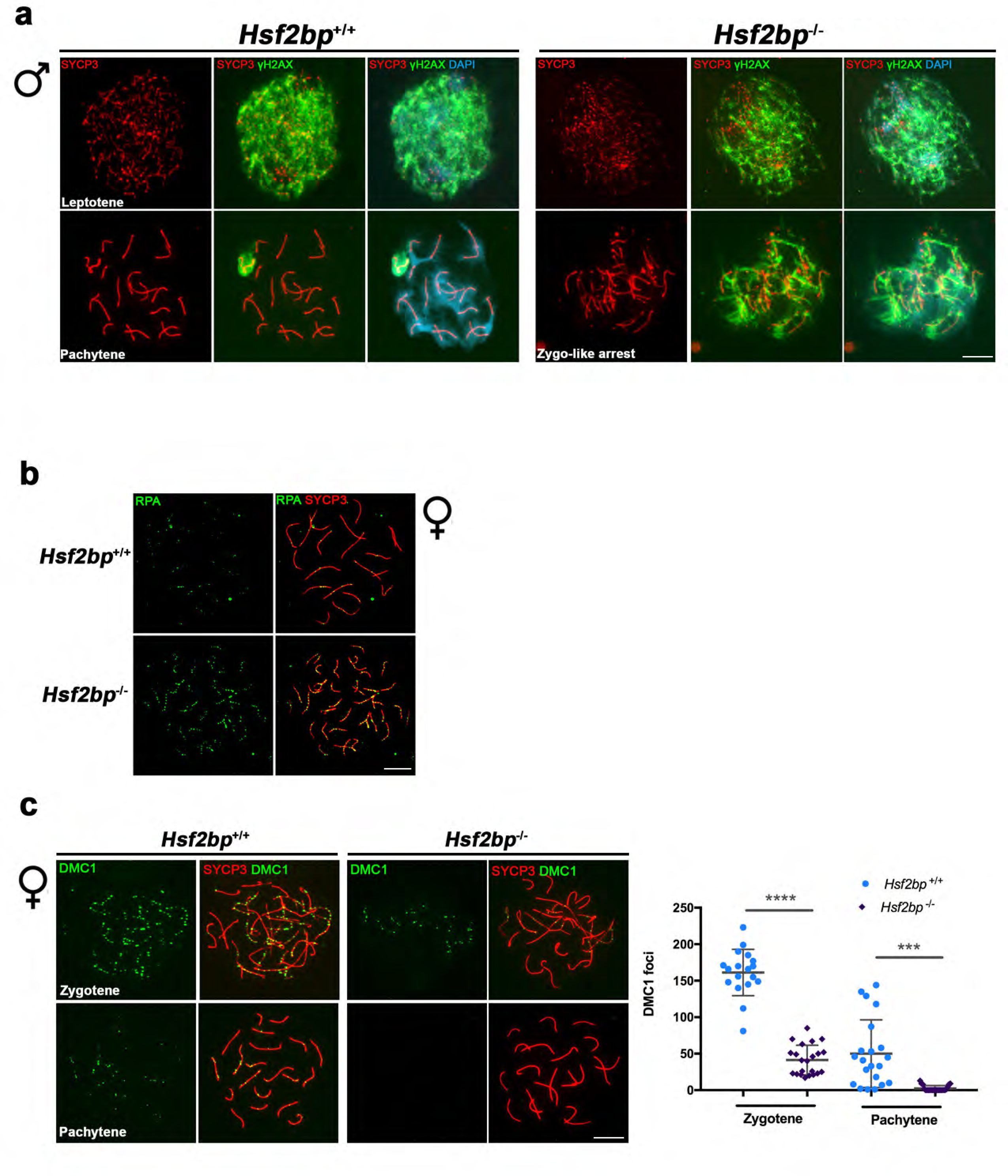
Defective DNA repair in *Hsf2bp*^-/-^ mice. (a) Double immunolabelling of SYCP3 (red) with γH2AX (green) in spermatocyte spreads from *Hsf2bp^+/+^*and *Hsf2bp^-/-^* mice. The strong labelling of γH2AX at leptotene indicates a correct induction of DSBs by SPO11 in both WT and KO mice. However, in the absence of HSF2BP, DSBs are not completely repaired and γH2AX signal is accumulated in the zygonema-like cells whereas in the WT the signal disappears from autosomes and remains restricted to the sexual chromosomes at pachytene. (b) Double immunofluorescence of RPA (green) and SYCP3 (red) in oocyte spreads from WT and *Hsf2bp*^-/-^ females showing an accumulation in the synapsis-defective cells of the knock-out. Bar in panels, 10 µm.

**Figure S9.**
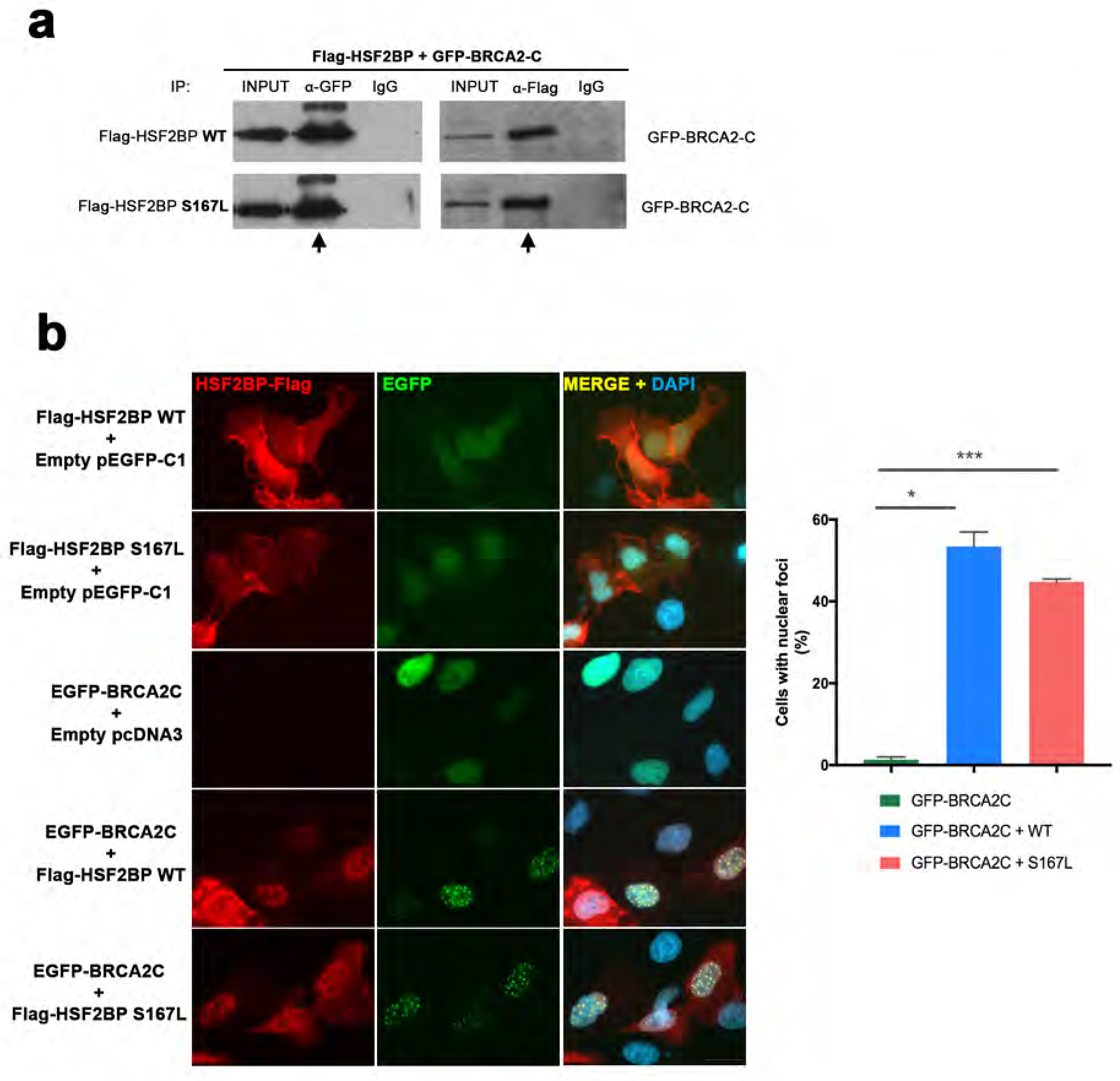
Comparative interaction of HSF2BP-S167L and HSF2BP-WT with BRCA2. (a) HEK293T cells were transfected with Flag-HSF2BP (WT, upper panel. S167L, panel below) and GFP-BRCA2-C (Cterm). Protein complexes were immunoprecipitated overnight with either an anti-Flag or anti-EGFP or IgGs (negative control), and were analysed by immunoblotting with the indicated antibody. Both HSF2BP variants (WT and S167L) co-immunoprecipitate (co-IP) with BRCA2-C similarly. (b) Double immunofluorescence of U2OS cells transfected with plasmids encoding Flag-HSF2BP (WT or S167L) and EGFP-BRCA2-C alone or together and immuno-detected with antibodies against Flag (Flag-HSF2BP, red) and against EGFP (EGFP-BRCA2-C, green). Transfected HSF2BP alone (both WT and S167L version) is delocalized and labels the whole cell (S167L less intense) whereas BRCA2-C shows a nuclear localization. When we co-transfected BRCA2-C with HSF2BP or HSF2BP-S167L they both gave rise to a similar nuclear punctate pattern (see quantification in the graph on the right of the panel). Bar in panel, 20µm. Welch’s t-test analysis: * p<0.05; *** p<0.001.

**Figure S10.**
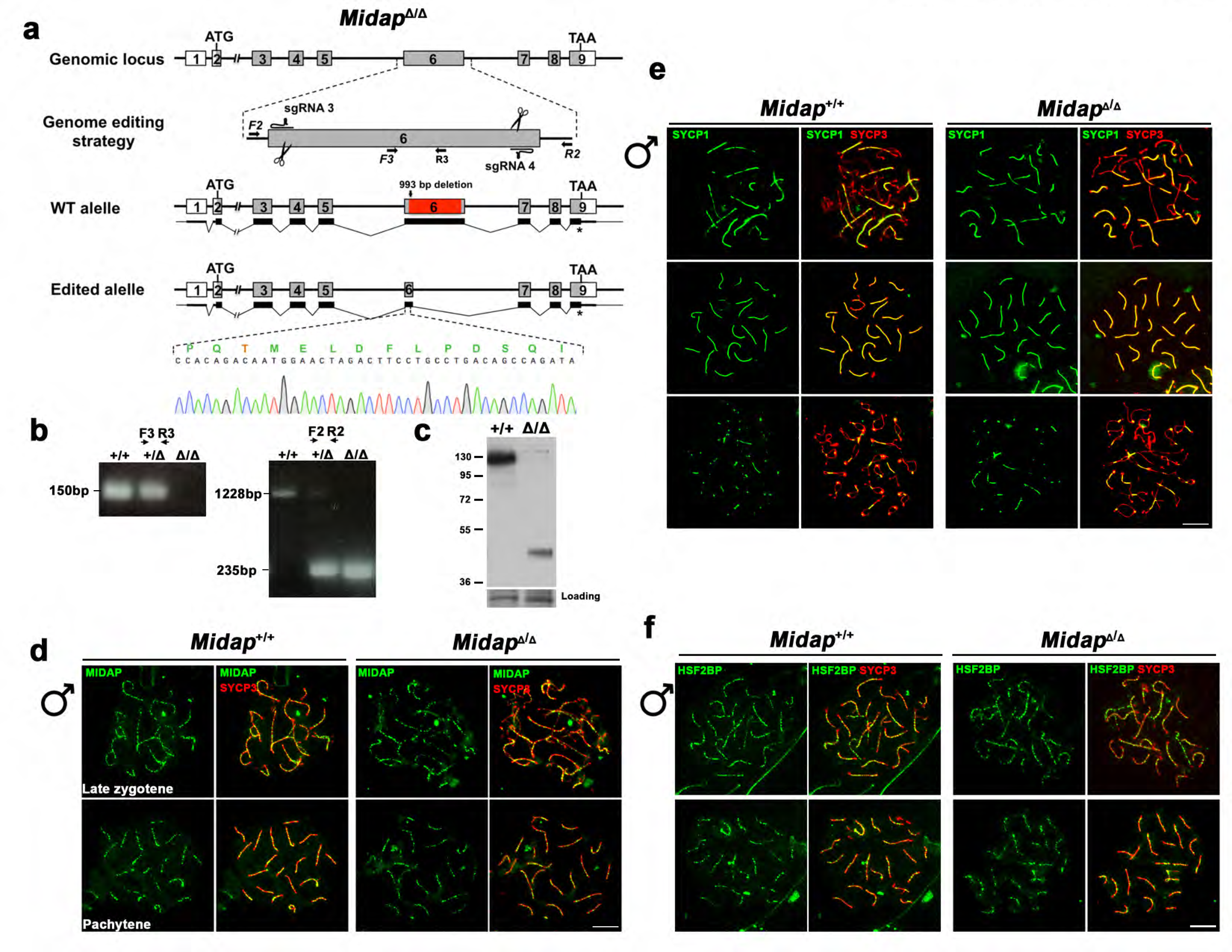
*Midap* Δ142-472 mutants do not show MIDAP and HSF2BP loading defects. (a) Diagrammatic representation of the mouse *Midap* locus (WT) and the genome editing strategy showing the sgRNAs located on exon 6 (see methods), the corresponding coding exons (light grey) and non-coding exons (open boxes). Thin (non-coding) and thick (coding sequences) lines under exons represent the expected transcript derived from wild-type and *Midap* edited allele. ATG, initiation codon; TAA and *, stop codon. The nucleotide sequence of the edited allele derived from PCR amplification of DNA from the *Midap***^Δ/Δ^** is indicated. Primers (F2, R2, F3 and R3) are represented by arrows. (b) PCR analysis of genomic DNA from three littermate progeny of *Midap^+/^*^Δ^ heterozygote crosses. The PCR amplification with primers F2 and R2 revealed 1228 and 235bp fragments for wild-type and delta alleles respectively. Wild-type (+/+), heterozygous (+/Δ), and homozygous delta (Δ/Δ) animals. The PCR amplification with the primers F3 and R3 revealed a 150bp fragment for wild-type and no PCR product for the delta allele. (c) Western blot analysis of protein extracts from *Midap^+/+^* and *Midap* ^Δ */*Δ^ mouse testis with a specific antibody against MIDAP. Note that the corresponding MIDAP (full-length) protein band migrates at 130 KDa (expected 65 KDa) due very likely to its high acidic content. The expected Δ142-472 band would be about 30 KDa but migrates higher for the same reason. An unspecific band was used as loading control. (d) Double immunofluorescence of spermatocytes at late zygotene and pachytene stage obtained from *Midap^+/+^* and *Midap* ^Δ/Δ^ mice using SYCP3 (red) and a rabbit polyclonal antibody against MIDAP (green). The Δ142-472 MIDAP localizes at recombination sites in the same manner than the full length MIDAP *in vivo*. (e) Double immunolabeling of SYCP1 (green) and SYCP3 (red) in *Midap^+/+^*and *Midap* ^Δ/Δ^ (Δ142-472) spermatocyte spreads showing no synapsis defects in presence of a MIDAP protein that lacks half of its total length. (f) Double immunolabeling of HSF2BP (green) and SYCP3 (red) in *Midap^+/+^*and *Midap* ^Δ/Δ^ (Δ142-472) spermatocyte spreads showing a normal localization of HSF2BP in absence of the central region of MIDAP. Bar in panels, 10 µm.

**Figure S11.**
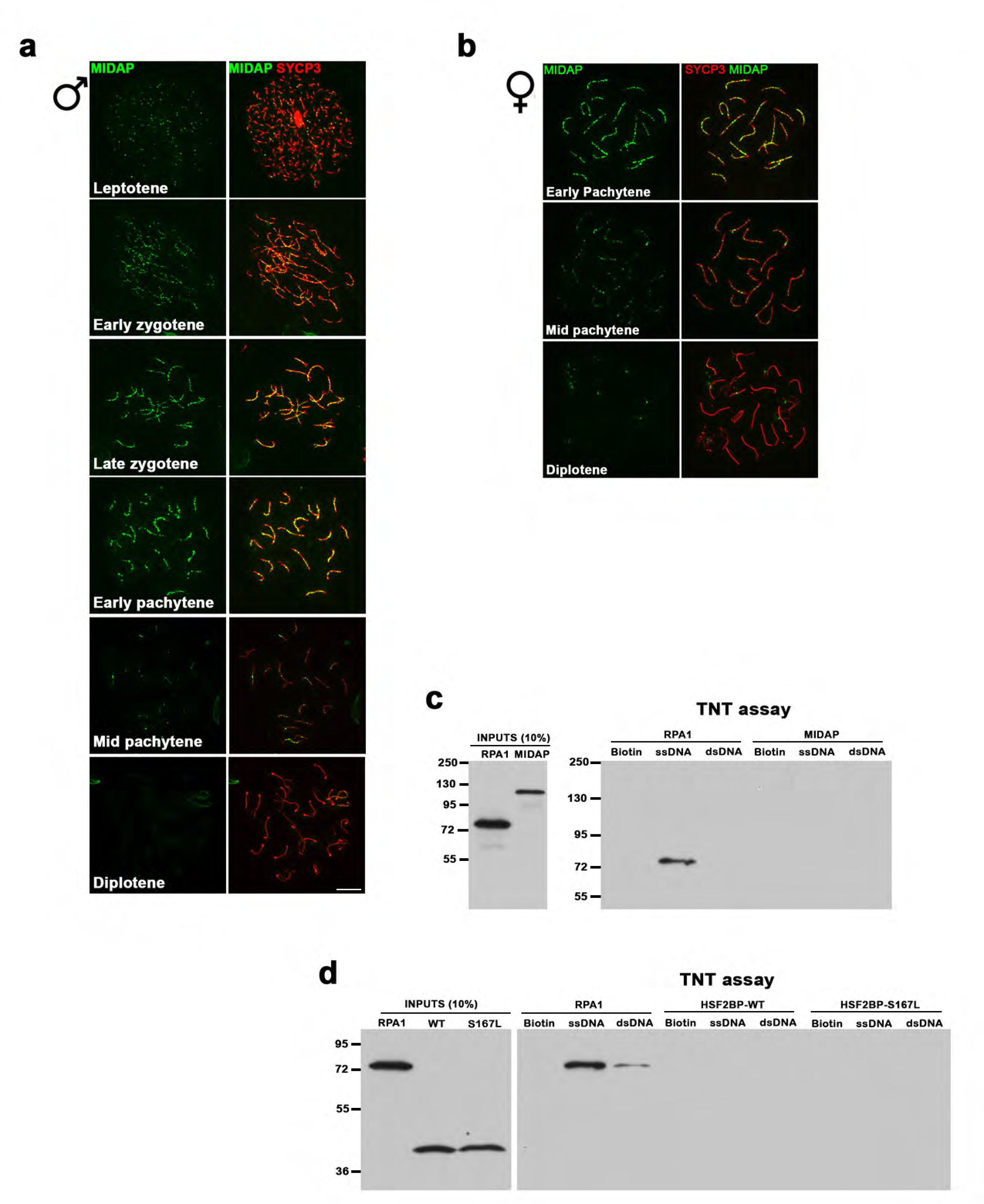
C19ORF57/MIDAP localizes at meiotic recombination nodules but has no DNA binding abilities. (a-b) Double immunolabeling of spermatocyte (a) and oocyte (b) spread preparations with a polyclonal antibody raised against a full-length recombinant MIDAP protein (green) and SYCP3 (red), showing that MIDAP localizes to meiotic recombination sites from leptotene to mid pachytene peaking at late zygotene. Bar in panels, 10 µm. (c-d) Flag-tagged MIDAP (c) and HSF2BP (WT and S167L) (d), and RPA1 (positive control in both) were produced in the cell-free system TNT and were incubated with biotin-coated magnetic beads or ssDNA or dsDNA (biotinylated) coated magnetic beads. The eluted DNA binding proteins were analysed by western blot. RPA1 was used as a positive control with ssDNA binding abilities.

**Figure S12.**
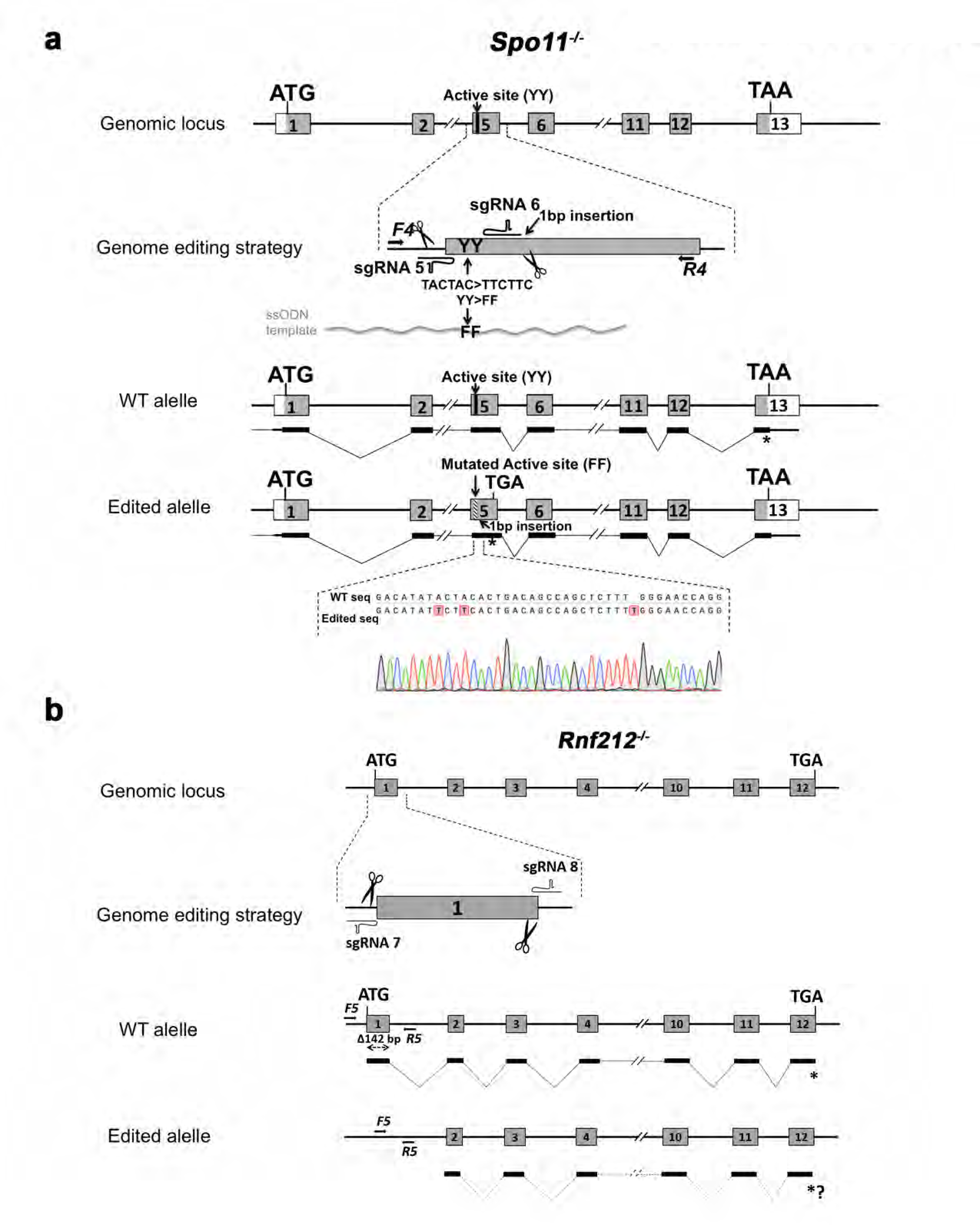
Generation and genetic characterization of *Spo11^-/-^* and *Rnf212^-/-^* mice. **(a)** Diagrammatic representation of the mouse *Spo11* locus (WT) and the genome editing strategy (see methods) showing the sgRNAs located on exon 5 and the ssODN carrying the desired mutation (TACTAC>TTCTTC p.YY137-138FF). The corresponding coding exons (light grey) and non-coding exons (open boxes) are represented. Thin (non-coding) and thick (coding sequences) lines under exons represent the expected transcript derived from wild-type and *Spo11* edited allele. ATG, initiation codon; TAA/TGA and *, stop codon. The nucleotide sequence of the WT and edited allele derived from PCR amplification of DNA from *Spo11*^+/+^ and *Spo11*^-/-^ mice is indicated. Arrows represent genotyping primers (F4 and R4). The mutant mice mimics the previously described null-mutants for *Spo11.* (b) Schematic representation of the mouse *Rnf212* locus (WT) and the genome editing strategy (see methods) showing the sgRNAs located upstream of the beginning of the ORF and in the exon 1. The corresponding coding exons (light grey) and non-coding exons (open boxes) are represented. Thin (non-coding) and thick (coding sequences) lines under exons represent the expected transcript derived from wild-type and *Rnf212* edited alleles. ATG, initiation codon; TGA and *, stop codon. Arrows represent genotyping primers (F5 and R5). This mutant behaves like the previously described null-mutants of *Rnf212*.

**Figure S13.**
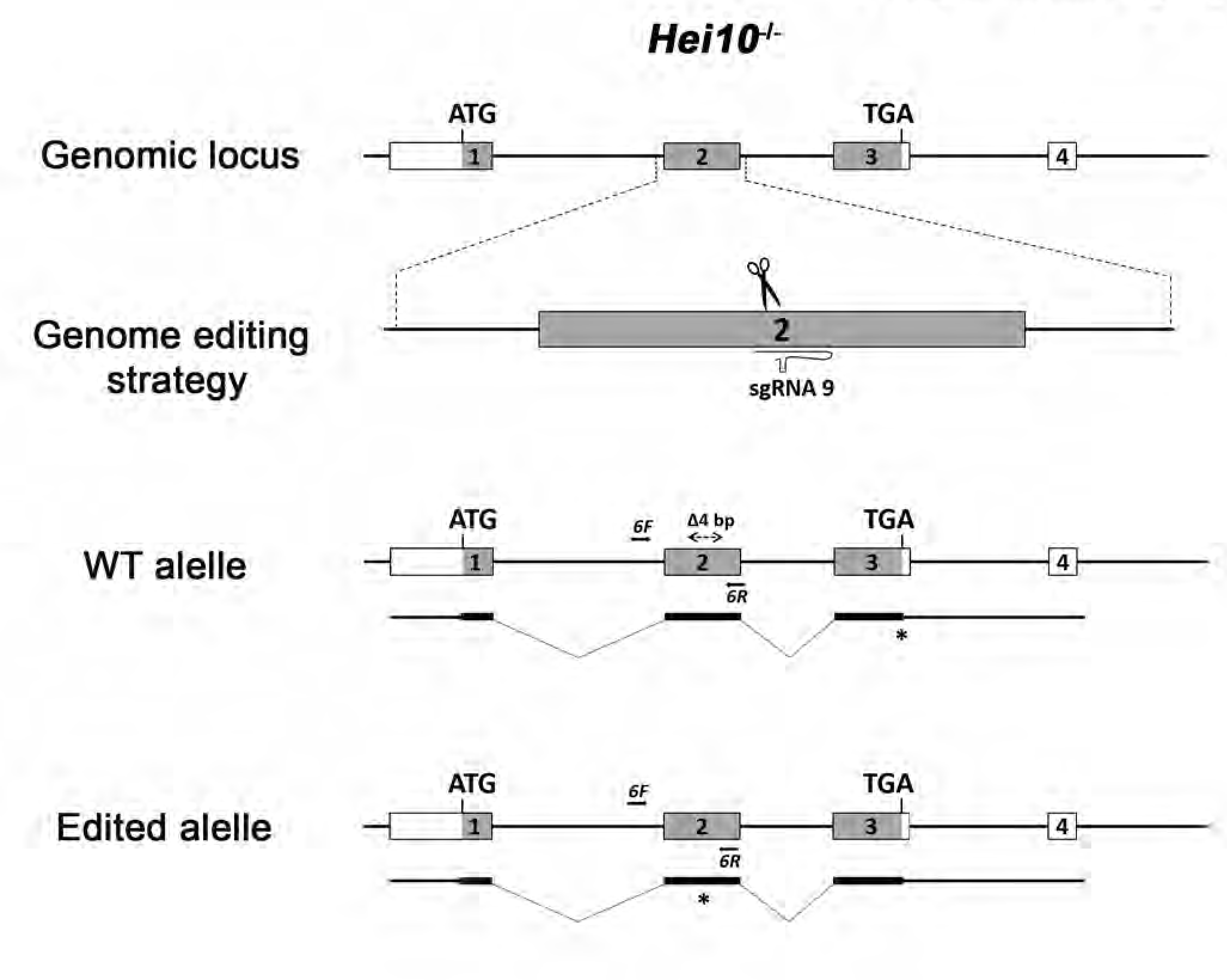
Generation and genetic characterization of *Hei10^-/-^* mice. (a) Schematic representation of the mouse *Hei10* locus (WT) and the genome editing strategy (see methods) showing the sgRNA located on the exon 2. The corresponding coding exons (light grey) and non-coding exons (open boxes) are represented. Thin (non-coding) and thick (coding sequences) lines under exons represent the expected transcript derived from wild-type and *Hei10* edited allele. ATG, initiation codon; TGA and *, stop codon. Arrows represent genotyping primers (F6 and R6). This mutant behaves like the previously described null-mutants of *Hei10*.

**Figure S14.**
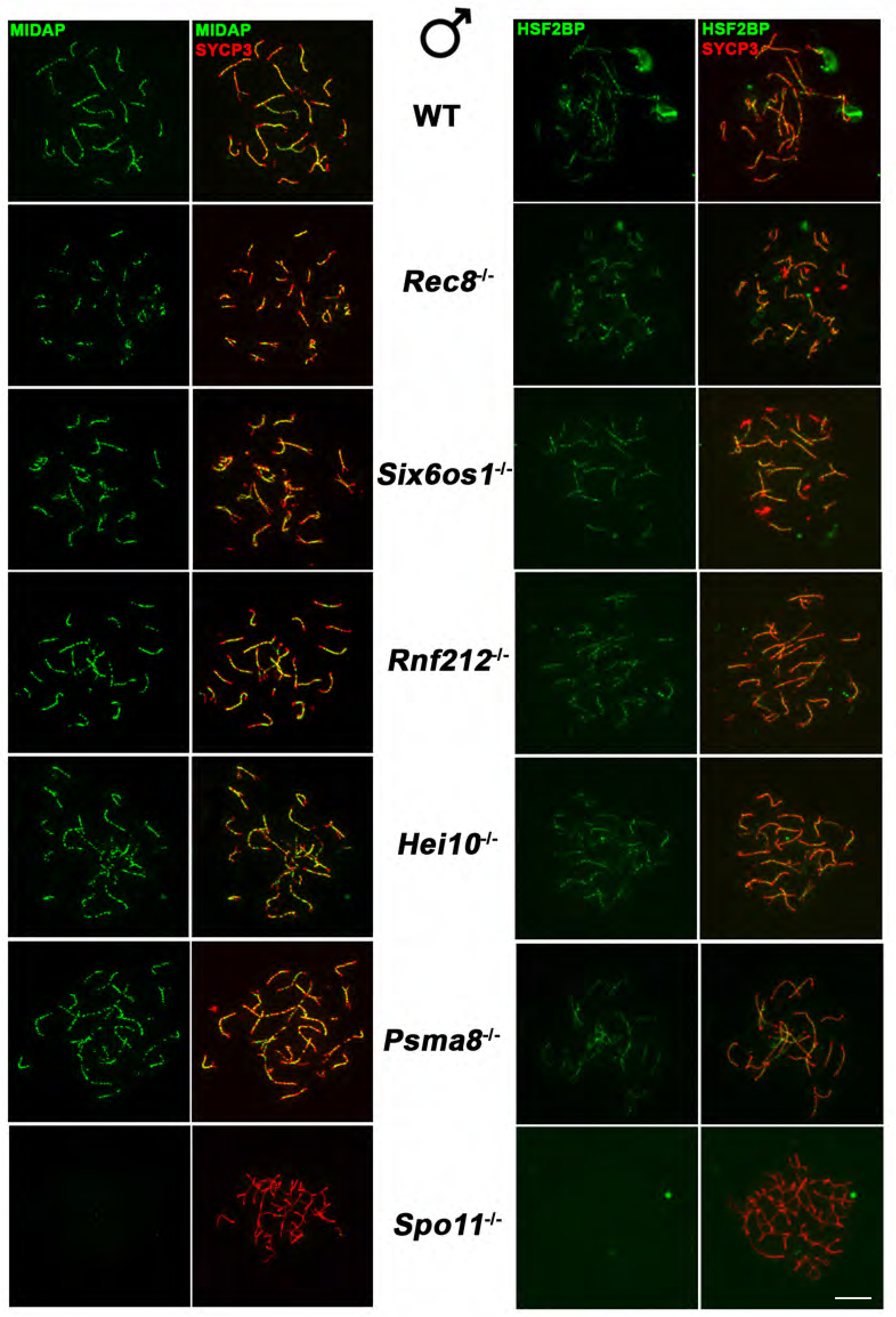
MIDAP loading depends on DSBs generation but not on synapsis. Double labelling of MIDAP (left)/HSF2BP (right) (green) and SYCP3 (red) in *Rec8*^−/−^, *Six6os1*^−/−^, *Rnf212*^−/−^, *Hei10*^−/−^, *Psma8*^−/−^ and *Spo11*^−/−^ mice showing a correct loading of MIDAP and HSF2BP in synapsis and recombination-defective mutants (*Six6os1*^−/−^, *Rec8*^−/−^*, Hei10*^−/−^ and *Rnf212*^−/−^,) but is absent in the *Spo11*^−/−^ spermatocytes indicating a total dependence on DSBs formation. These results indicate that both MIDAP and HSF2BP can be genetically positioned at the early events after DSB generation. Bar in panels, 10µm.

**Figure S15.**
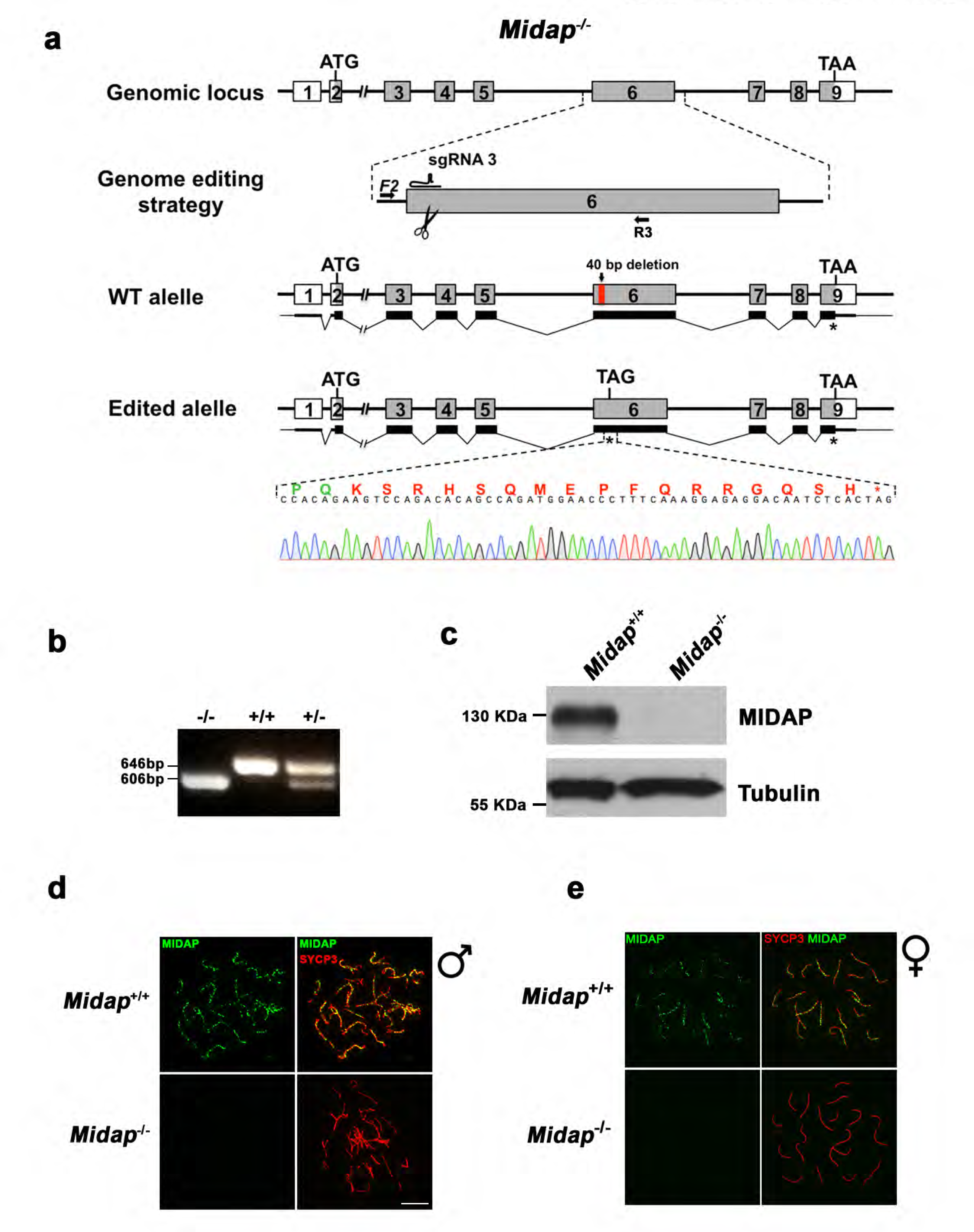
Generation and genetic characterization of *Midap* knock-out mice. (a) Diagrammatic representation of the mouse *Midap* locus (WT) and the genome editing strategy showing the sgRNA located on exon 6 (see methods), the corresponding coding exons (light grey) and non-coding exons (open boxes). Thin (non-coding) and thick (coding sequences) lines under exons represent the expected transcript derived from wild-type and *Midap* edited allele. ATG, initiation codon; TAA/TAG and *, stop codon. The nucleotide sequence of the edited allele derived from PCR amplification of DNA from the *Midap***^-/-^** is indicated. Primers (F2 and R3) are represented by arrows. (b) PCR analysis of genomic DNA from three littermate progeny of *Midap^+/-^* heterozygote crosses. The PCR amplification with primers F2 and R3 revealed 646 and 606 bp fragments for wild-type and disrupted alleles respectively. Wild-type (+/+), heterozygous (+/-), and homozygous knock-out (-/-) animals. (c) Western blot analysis of protein extracts from wild type and *Midap^-/-^* testis with a specific antibody against whole recombinant MIDAP protein. Tubulin was used as loading control. (d-e) Double immunofluorescence of spermatocytes at late zygotene-like arrested stage (d) and pachytene oocytes (e) obtained from *Midap^+/+^* and *Midap* ^-/-^ mice using SYCP3 (red) and a rabbit polyclonal antibody against MIDAP (green). No signal is detected in the *Midap* ^-/-^ demonstrating that is a null allele. Bar in panels, 10µm.

**Figure S16.**
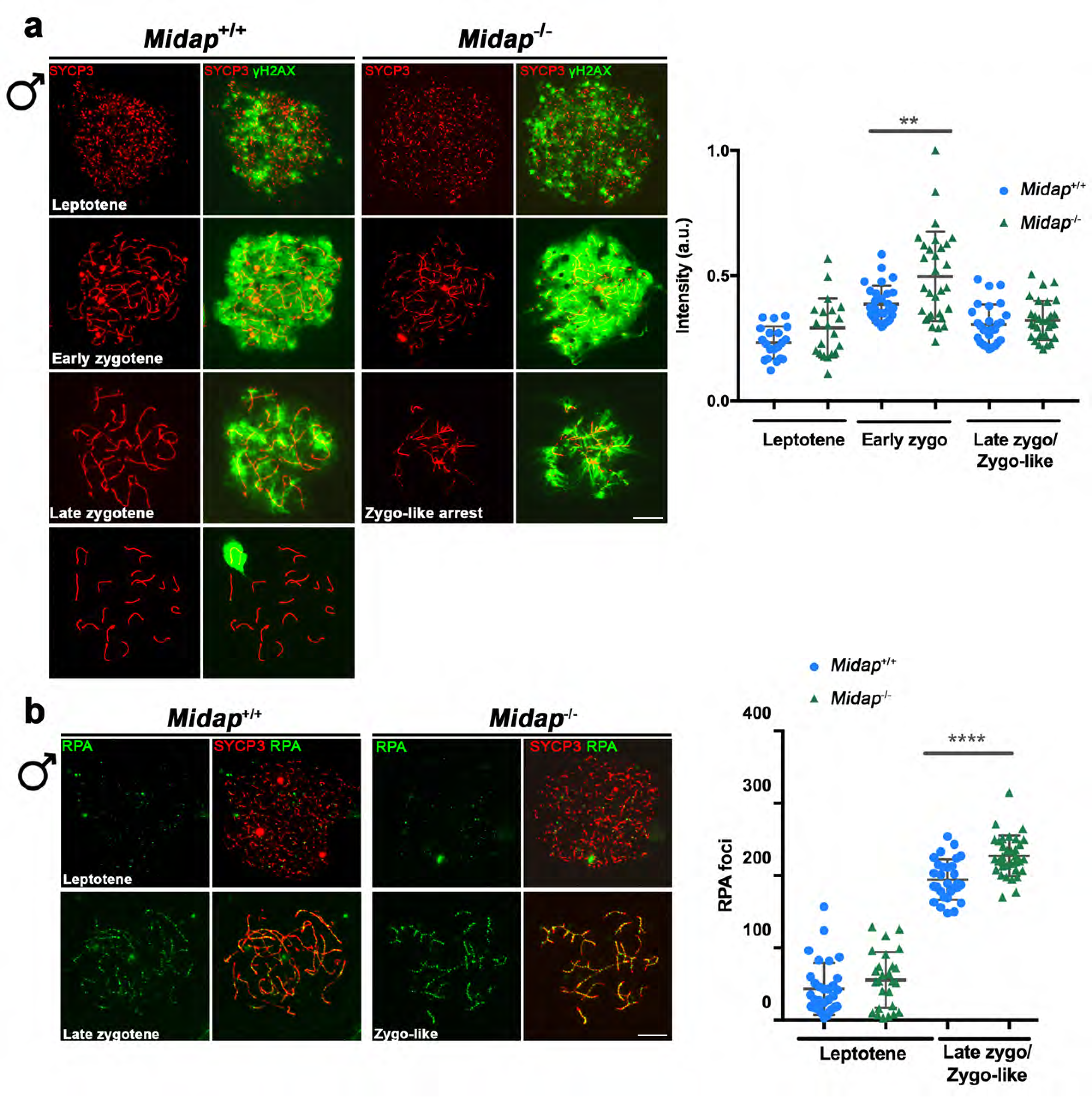
DSBs are formed but not properly repaired in *Midap*-deficient mice and mimic the phenotype of *Hsf2bp*-deficient mice. (a) Double labeling of γH2AX (green) and SYCP3 (red) in spermatocyte spreads from WT and *Midap*-deficient mice. γH2AX labeling appears in leptotene in both WT and *Midap* KO spermatocytes and increase during zygotene. However, whereas in the wild-type the signal decreases in pachytene, *Midap*-deficient mice show an accumulation of γH2AX in the zygotene-like arrest. The plot on the right represents the quantification of γH2AX intensity in both WT and KO spermatocytes. (b) Double immunolabeling of RPA (green) and SYCP3 (red) in spermatocyte spreads from *Midap^+/+^* and *Midap^-/-^* males. RPA accumulates at the late zygotene arrest of the *Midap*-deficient mouse indicating a defective DNA repair in these mutants. The plot on the right of the panel represents the quantification of RPA foci on each genotype and stage. The quantification of RPA foci was done in parallel in the WT, *Hsf2bp*^S167L/S167L^, *Hsf2bp*^-/-^ and *Midap^-/-^* mice. For this reason the data for the WT are the same on this figure and the figures 4a). Bar in panels, 10µm. Welch’s t-test analysis: ** p<0.01; **** p<0.0001.

**Figure S17.**
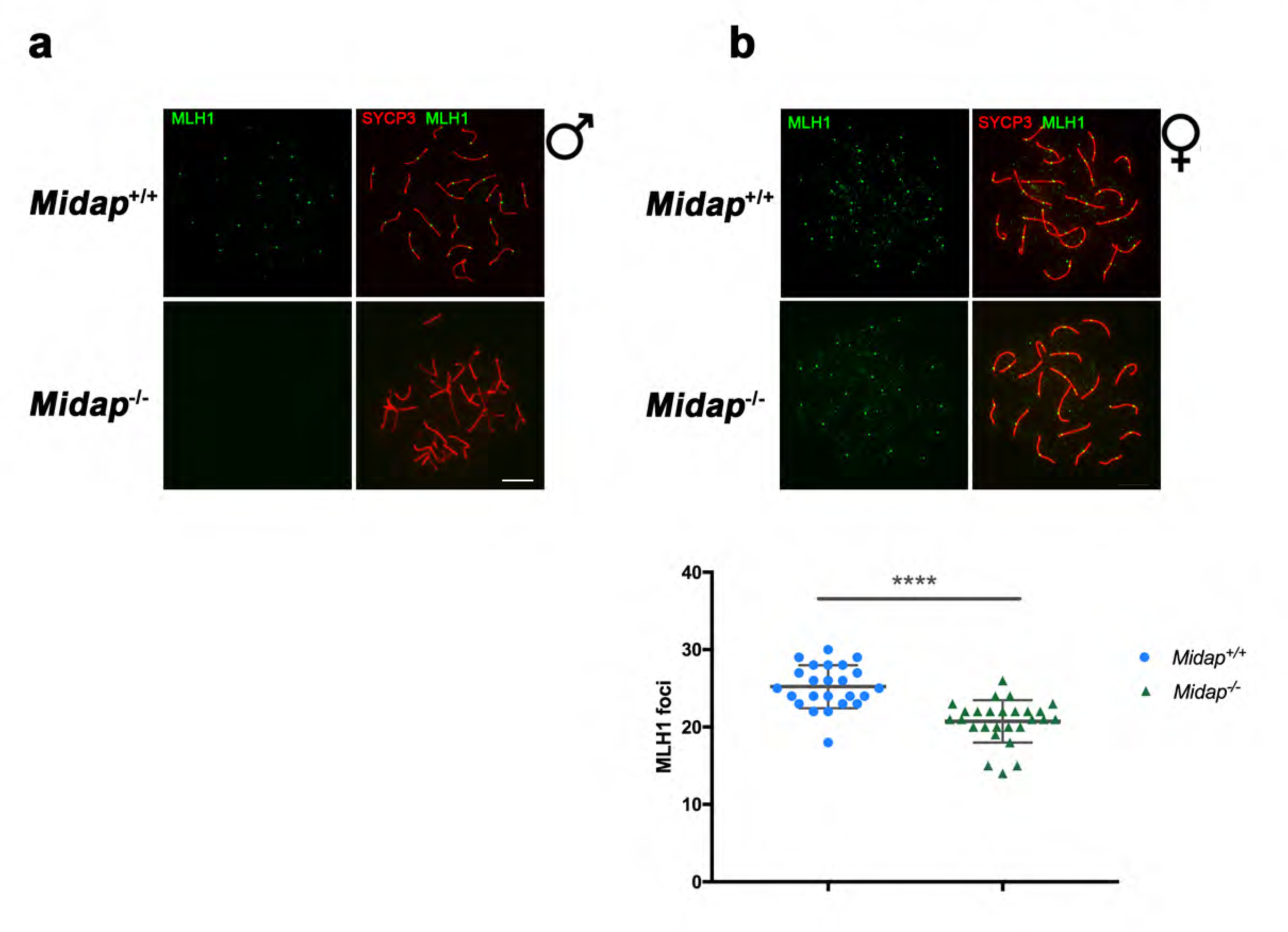
Cross-over formation is abolished in *Midap^-/-^* males and reduced in females. (a-b) Double immunofluorescence of MLH1 (green) and SYCP3 (red) in *Midap^+/+^* and *Midap^-/-^* (a) spermatocytes and (b) oocytes showing the absence and reduction of MLH1 labelling, respectively. Plot under the panel in (b) shows the quantification of MLH1 foci in oocytes. The quantification of MLH1 foci in *Midap^+/+^*, *Midap^-/-^* and *Hsf2bp*^S167L/S167L^ was done in parallel. For this reason the data for the WT are the same on this figure and the figure 5d. Welch’s t-test analysis: **** p<0.0001.

**Figure S18.**
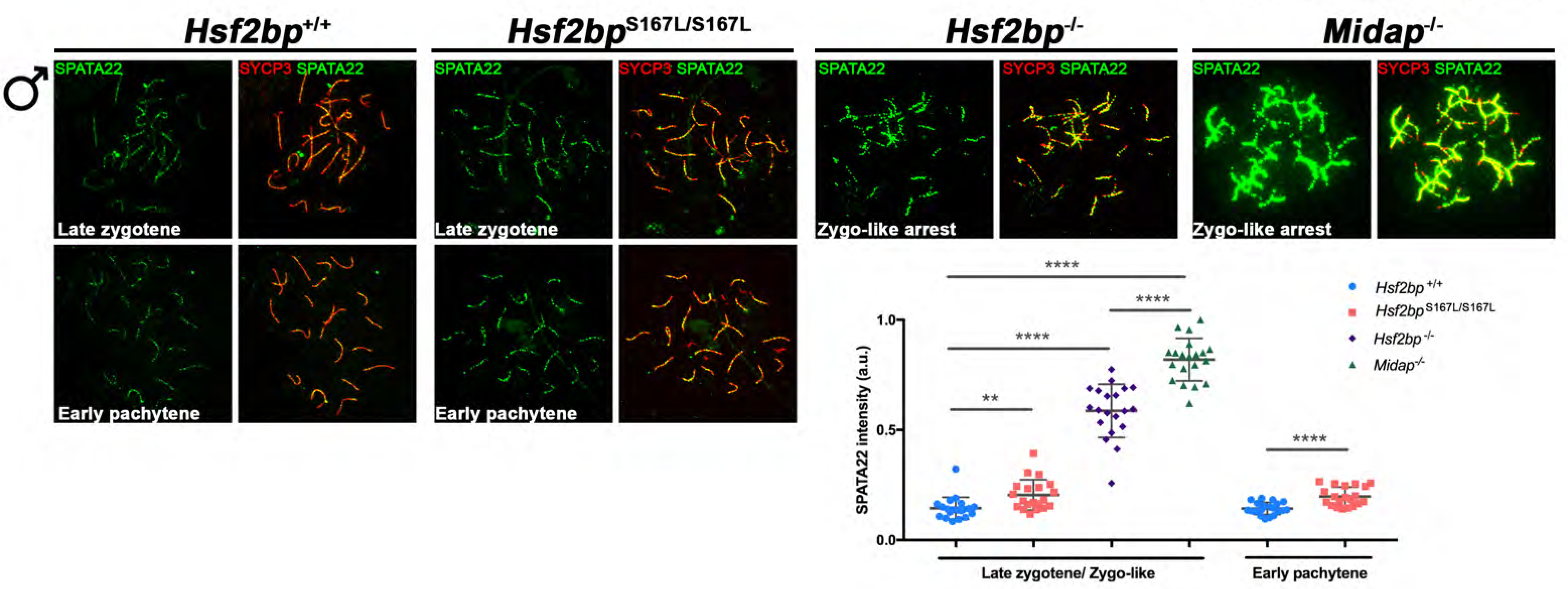
The ssDNA binding protein SPATA22 is accumulated in *Hsf2bp^-/-^, Hsf2bp^S167L/S167L^* and *Midap^-/-^* mutants. Double labelling of SPATA22 (green) and SYCP3 (red) in spermatocyte spreads from WT, *Midap*- and *Hsf2bp*-deficient mice and in the *Hsf2bp^S167L/S167L^* mutant. SPATA22 is highly accumulated in both knock-out spermatocytes and shows a milder accumulation in the *Hsf2bp^S167L/S167L^* spermatocytes.

**Figure S19.**
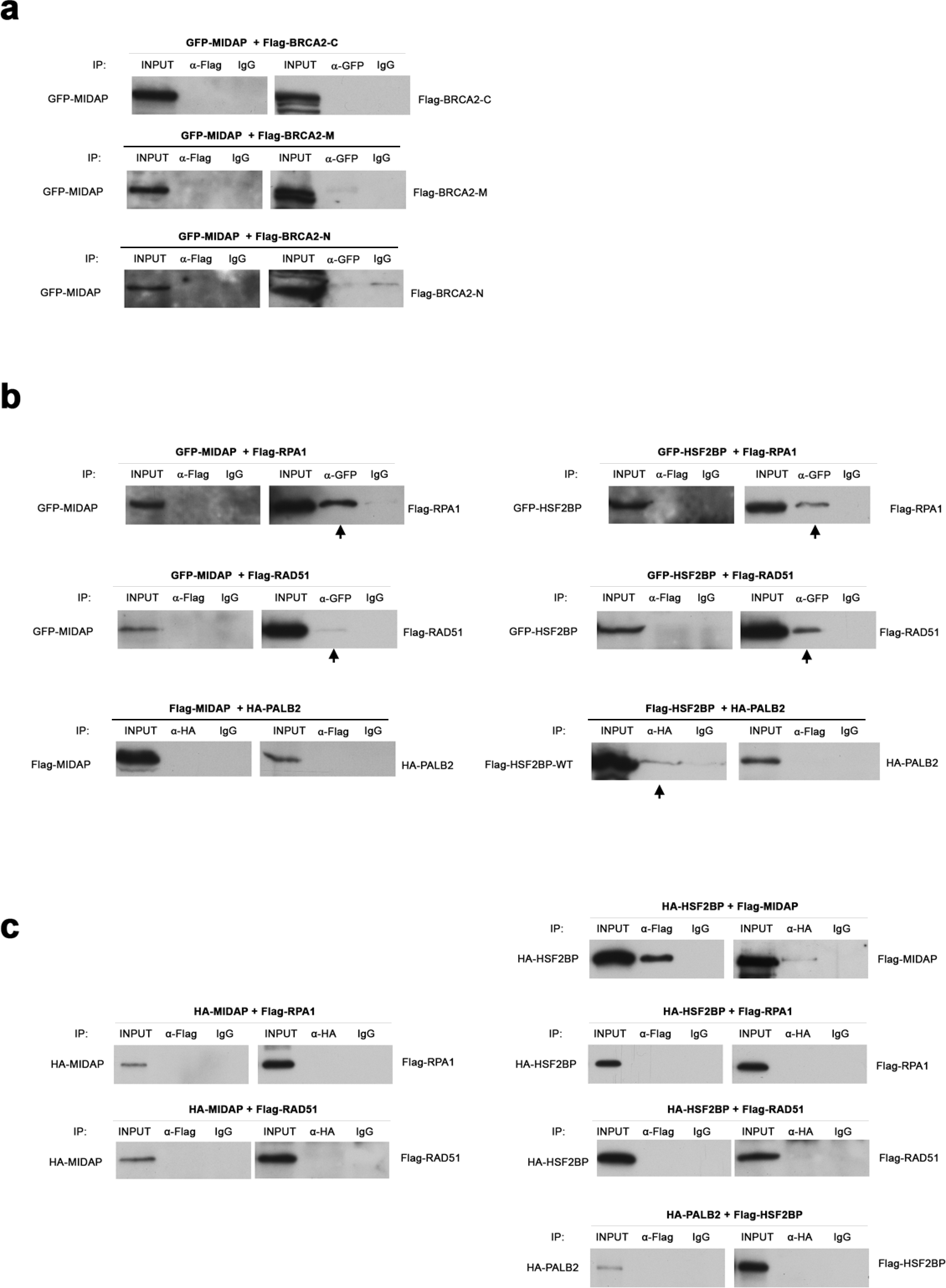
Co-immunoprecipitations of MIDAP, HSF2BP, RPA, RAD51 and PALB2 expressed from transfected HEK293 cells and from TNT assays. (a-b) HEK293T cells were transfected with GFP- or Flag-tagged MIDAP/HSF2BP and different tagged potential interactors. Protein complexes were immunoprecipitated overnight with an anti-Flag, anti-HA or anti-EGFP or IgGs (negative control), and were analysed by immunoblotting with the indicated antibody. (a) MIDAP does not co-immunoprecipitates with BRCA2-N, BRCA2-M or BRCA2-C. (b) MIDAP (left panels) does not co-immunoprecipitates with PALB2 but co-immunopricpitates with RPA1 and RAD51 (mildly). HSF2BP (right panels) co-immunoprecipitates with RPA1, RAD51 and PALB2 (mildly). (c) Flag or HA-tagged HSF2BP and MIDAP and their potential interactors (RPA1, RAD51 and PALB2) were expressed in a cell-free TNT system. Proteins were complexed during 30 minutes at RT an immunoprecipitated overnight with either an anti-Flag, anti-HA or IgGs (negative control), and analysed by immunoblotting with the indicated antibody. The unique positive co-immunoprecipitation was between HSF2BP with MIDAP.

## Supplementary Tables

**Table S1.**
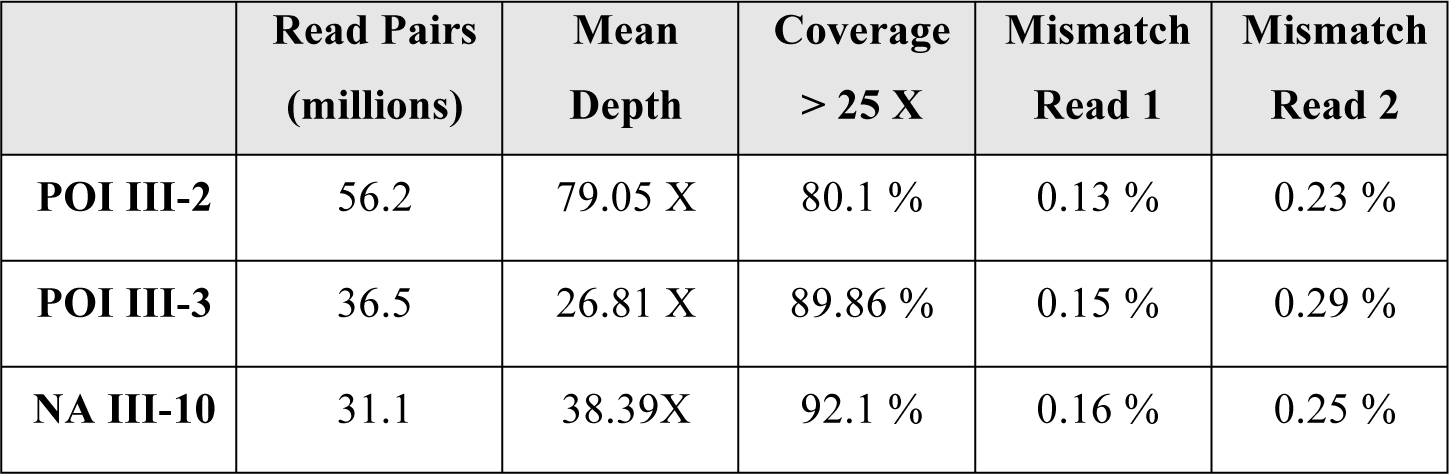
Whole Exome Sequencing and mapping data for the POI patients and the unaffected sister.

|  | Read Pairs<br>(millions) | Mean<br>Depth | Coverage<br>> 25 X | Mismatch<br>Read 1 | Mismatch<br>Read 2 |
| --- | --- | --- | --- | --- | --- |
| **POI III-2** | 56.2 | 79.05 X | 80.1 % | 0.13 % | 0.23 % |
| **POI III-3** | 36.5 | 26.81 X | 89.86 % | 0.15 % | 0.29 % |
| **NA III-10** | 31.1 | 38.39X | 92.1 % | 0.16 % | 0.25 % |

**Table S2:**
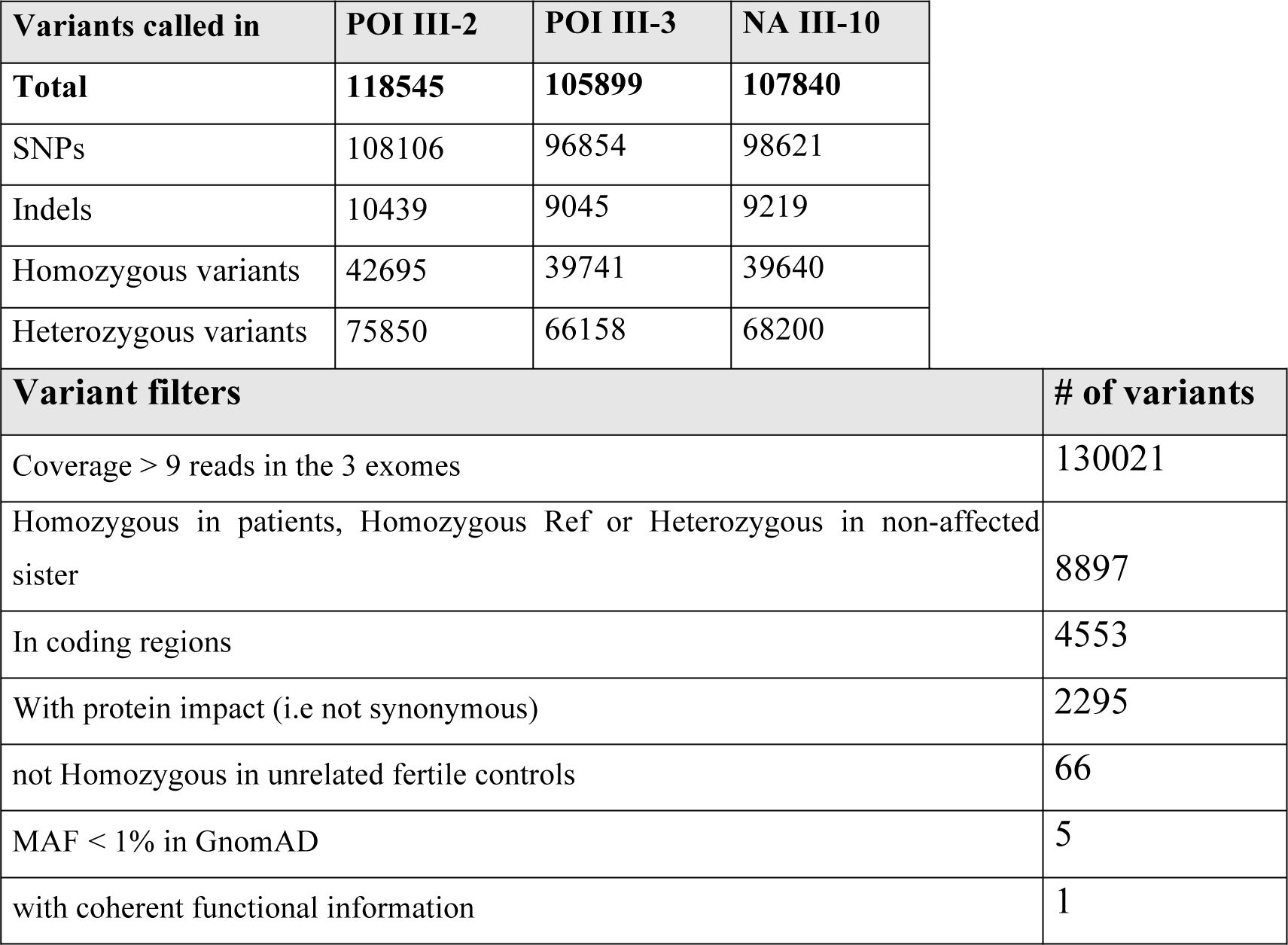
Filtering of the variants identified by Whole Exome Sequencing

**Table S3:**
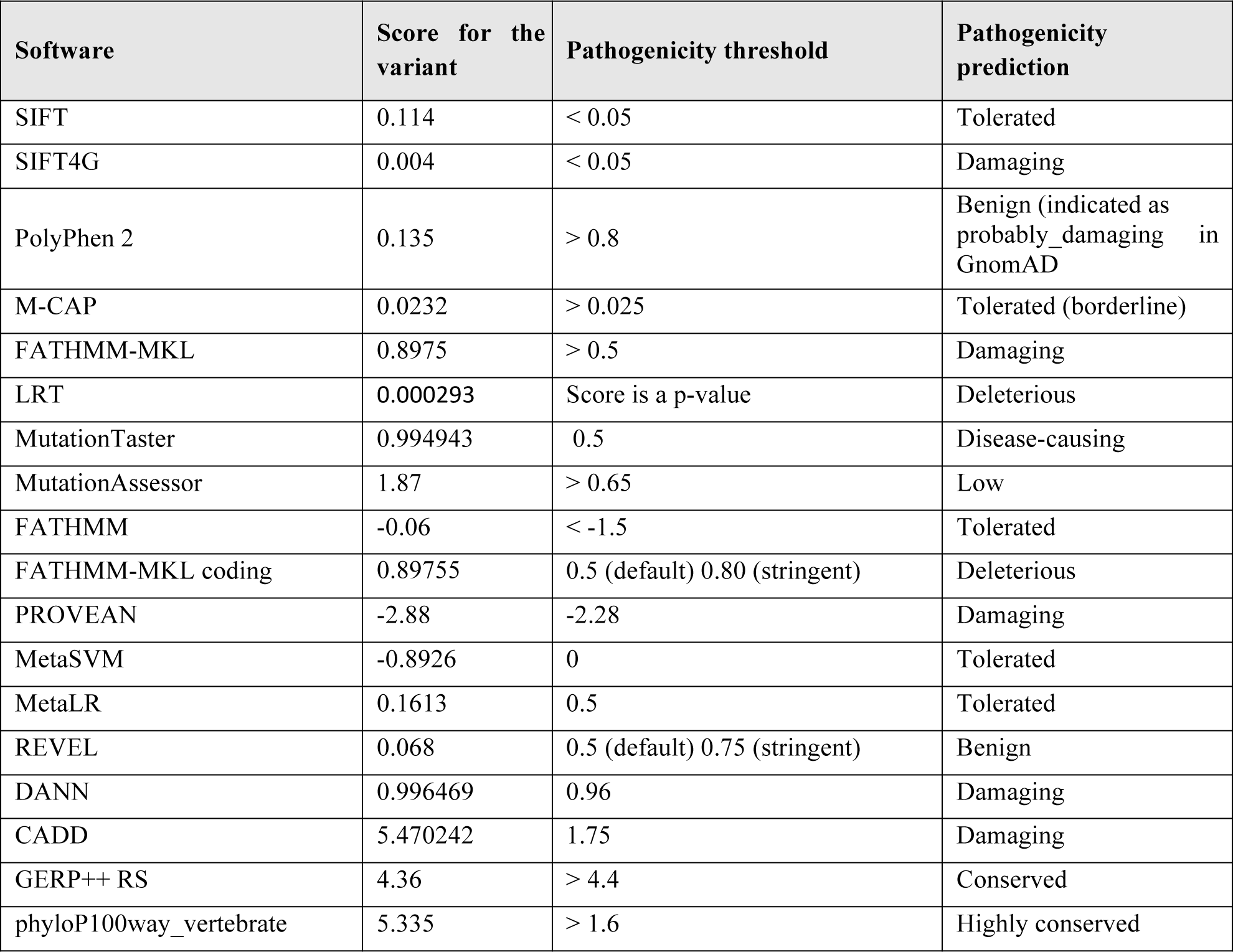
Pathogenicity predictions for the S167L variant in *HSF2BP*

**Table S4.**
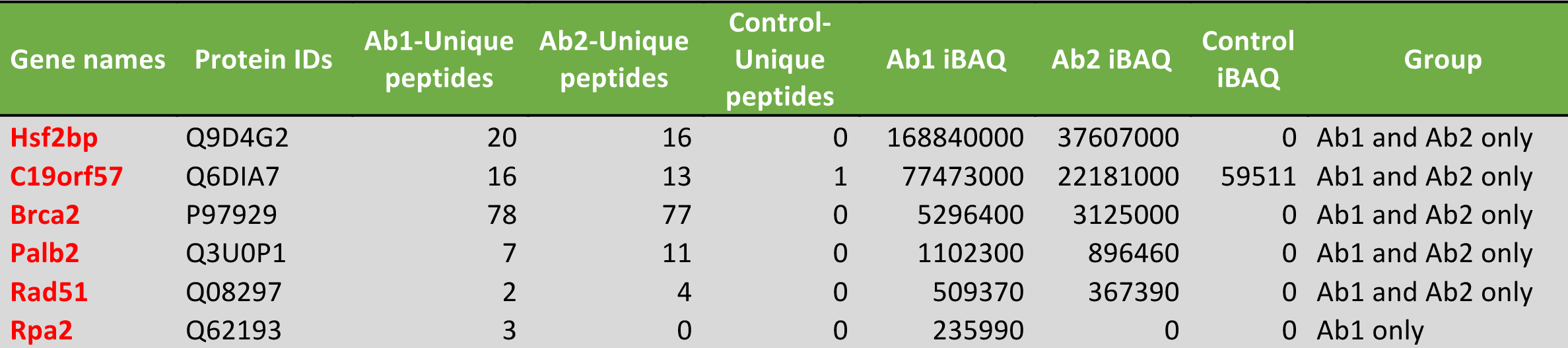
Meiotic recombination players identified as putative MIDAP interactors by IP-MS of whole testis extracts with two antibodies (Ab1 and Ab2) against MIDAP. See extended data on Table S5.

| Gene names | Protein IDs | Ab1-Unique peptides | Ab2-Unique peptides | Control-Unique peptides | Ab1 iBAQ | Ab2 iBAQ | Control iBAQ | Group |
| --- | --- | --- | --- | --- | --- | --- | --- | --- |
| Hsf2bp | Q9D4G2 | 20 | 16 | 0 | 168840000 | 37607000 | 0 | Ab1 and Ab2 only |
| C19orf57 | Q6DIA7 | 16 | 13 | 1 | 77473000 | 22181000 | 59511 | Ab1 and Ab2 only |
| Brca2 | P97929 | 78 | 77 | 0 | 5296400 | 3125000 | 0 | Ab1 and Ab2 only |
| Palb2 | Q3U0P1 | 7 | 11 | 0 | 1102300 | 896460 | 0 | Ab1 and Ab2 only |
| Rad51 | Q08297 | 2 | 4 | 0 | 509370 | 367390 | 0 | Ab1 and Ab2 only |
| Rpa2 | Q62193 | 3 | 0 | 0 | 235990 | 0 | 0 | Ab1 only |

**Table S6.**
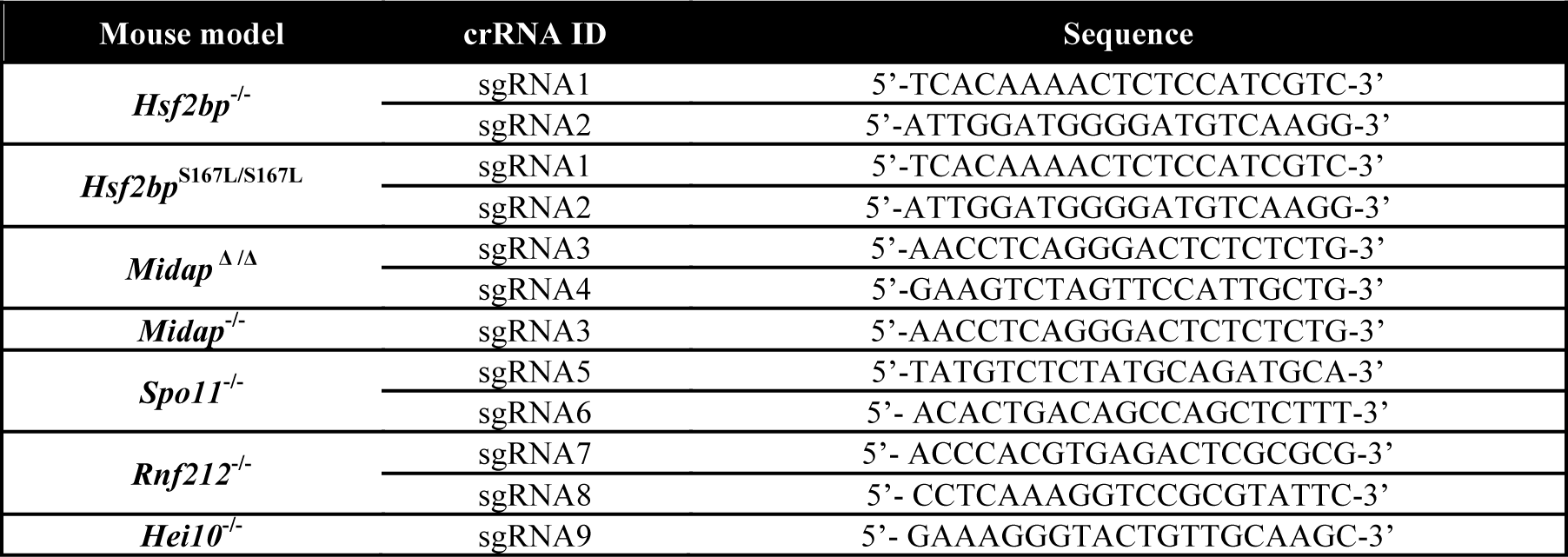
crRNAs employed for the generation of the different mouse models.

**Table S7.**
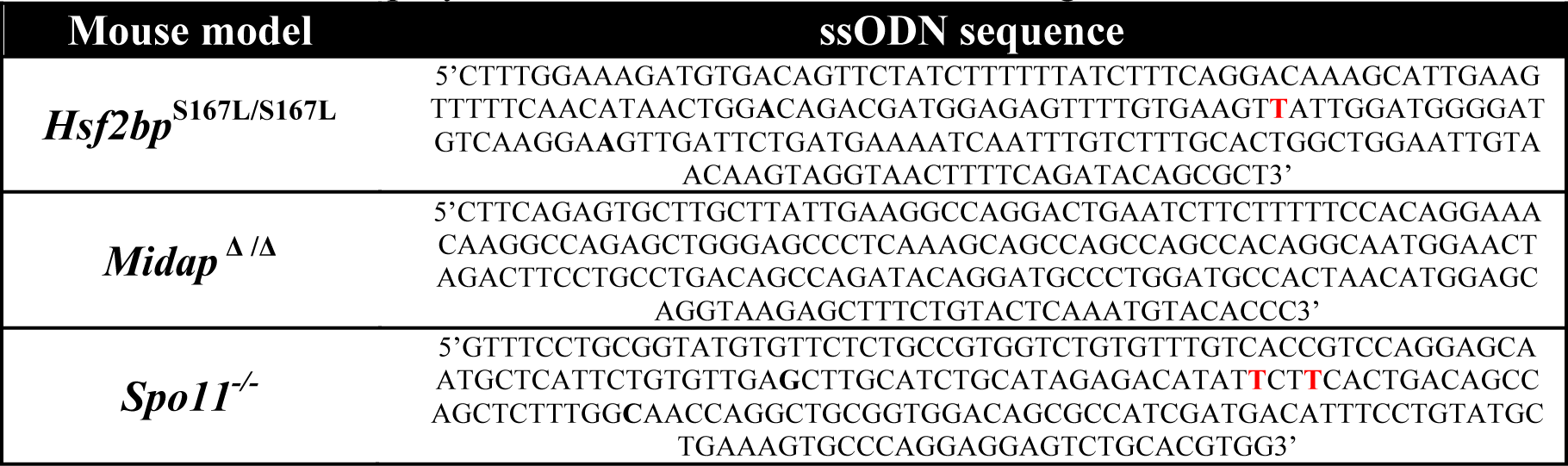
ssODN employed for the different mouse models generation.

**Table S8.**
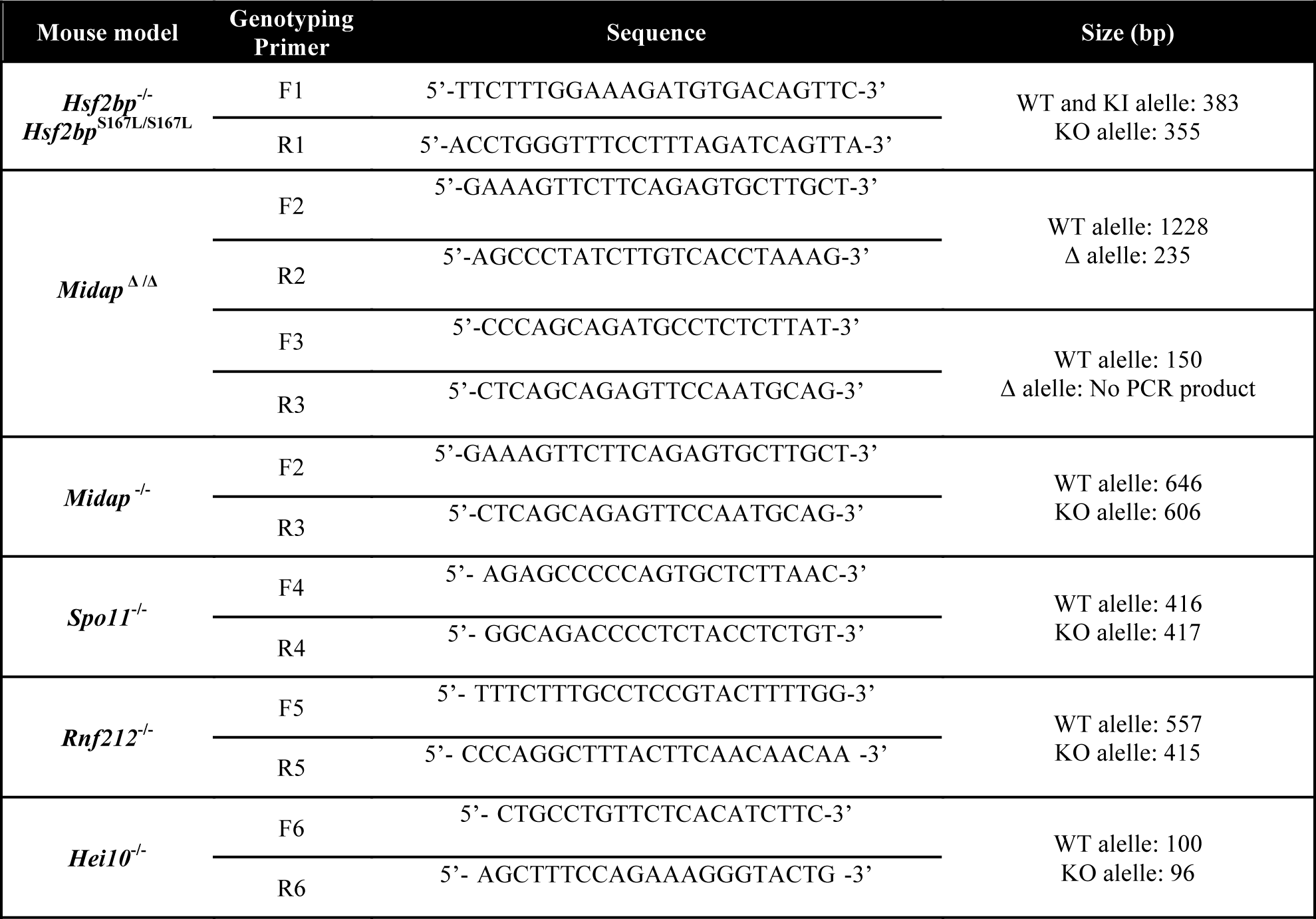
Primers and expected product size for genotyping each mutant mouse model.

